# Saponin Nanoparticle Adjuvants Incorporating Toll-Like Receptor Agonists Drive Distinct Immune Signatures and Potent Vaccine Responses

**DOI:** 10.1101/2023.07.16.549249

**Authors:** Ben S. Ou, Julie Baillet, Maria V. Filsinger Interrante, Julia Z. Adamska, Xueting Zhou, Olivia M. Saouaf, Jerry Yan, John H. Klich, Carolyn K. Jons, Emily L. Meany, Adian S. Valdez, Lauren Carter, Bali Pulendran, Neil P. King, Eric A. Appel

## Abstract

Over the past few decades, the development of potent and safe immune-activating adjuvant technologies has become the heart of intensive research in the constant fight against highly mutative and immune evasive viruses such as influenza, SARS-CoV-2, and HIV. Herein, we developed a highly modular saponin-based nanoparticle platform incorporating toll-like receptor agonists (TLRas) including TLR1/2a, TLR4a, TLR7/8a adjuvants and their mixtures. These various TLRa-SNP adjuvant constructs induce unique acute cytokine and immune-signaling profiles, leading to specific Th-responses that could be of interest depending on the target disease for prevention. In a murine vaccine study, the adjuvants greatly improved the potency, durability, breadth, and neutralization of both COVID-19 and HIV vaccine candidates, suggesting the potential broad application of these adjuvant constructs to a range of different antigens. Overall, this work demonstrates a modular TLRa-SNP adjuvant platform which could improve the design of vaccines for and dramatically impact modern vaccine development.

**Teaser:** Saponin-TLRa nanoadjuvants provide distinct immune signatures and drive potent, broad, durable COVID-19 and HIV vaccine responses.

## Introduction

The development of prophylactic vaccines against infectious diseases has been at the heart of intensive scientific research for the past hundred years, but the emergence of global pandemics such as SARS-CoV-2, HIV, and influenza have further catalyzed their development (*1*). Yet, many commercial vaccines comprise traditional live-attenuated or inactivated virus technologies, both of which suffer from complex manufacturing processes and potential safety hazards (*2*). Over the last few decades, protein-based subunit vaccine approaches have been shown to offer desirable characteristics in terms of safety, cost effectiveness, scalability, and manufacturability. Subunit vaccines leverage the use of viral protein antigens paired with immune stimulating molecules, also known as adjuvants, that are essential in promoting the magnitude and durability of the immune response.

To date, only a handful of adjuvants are clinically licensed in vaccine formulations and approved by the Food and Drug Administration (FDA), including more traditional insoluble aluminum salts (Alum) and oil-in-water emulsions such as MF59 and AS03 (*3*). Numerous adjuvant candidates targeting specific molecular pathways have been in development recently, many of which offer promising improvements in the overall efficacy of subunit vaccines. A specific class of adjuvants that has been gaining attention is toll-like receptor agonists (TLRas) (*4, 5*). These agonists include Pam3CysSerLys4 (Pam3CSK4; TLR1/2a), (Polyinosinic)polycytidylic acid (Poly(I:C); TLR3a), Monophosphoryl lipid (MPL; TLR4a), imidazoquinolines such as Resiquimod or 3M-052 (TLR7/8a), and CpG-ODN (TLR9a). While these molecules differ in their physical and chemical properties, and include lipids, small molecules, single-stranded DNA, and double-stranded RNA, they all activate different toll-like receptors (TLRs) commonly expressed by antigen-presenting cells (APCs) to drive distinct immune signaling pathways. Recent FDA approvals of vaccines comprising CpG-ODN 1018 (Dynavax), AS04 (GSK; MPL/Alum), AS01 (GSK; liposomes comprising MPL), along with numerous ongoing clinical trials using other promising TLRas such as 3M-052 (3M/AAHI), reinforce their promise as adjuvants to improve the efficacy of vaccines.

While TLRa adjuvants have been the primary focus in vaccine design, other non-TLR-based adjuvants have also been of interest. For example, saponins such as Quil-A, natural triterpene glycosides from Quillaja, are leading adjuvant molecules in development. Although saponins can be toxic when delivered in a soluble form, they are well tolerated when delivered in nanoparticles such as the well-known honeycomb-like structure ISCOMATRIX (*6–12*). These well-defined nanostructures are 30-70 nm in size and are generated by spontaneous self-assembly of Quil-A saponins with cholesterol and phospholipids. These adjuvants have been successfully used for decades in vaccine formulations and have been demonstrated to be highly potent and safe. Indeed, Matrix-M (Novavax), a self-assembled saponin-based nanostructure adjuvant comparable to ISCOMATRIX, was approved by the FDA in October 2022 as part of the Novavax COVID-19 vaccine. While the mechanism of immune stimulation of these self-assembled saponin-based nanostructure adjuvants has not yet been fully elucidated, recent reports suggest they increase lymph flow and lymph node permeability to enhance antigen acquisition by B cells in draining lymph nodes (dLNs) (*7, 12*). More generally, intensive research has demonstrated the advantage of formulating adjuvants as nanoparticles, to improve their stability, solubility, cellular uptake, immunogenicity, and safety, all ultimately augmenting vaccine immune responses favorably (*13–19*).

Few attempts have been made to combine both TLRas and saponin adjuvants into the same nanostructure (*7, 20–24*). For example, AS01 adjuvant, which is part of the licensed Shingrix and Mosquirix vaccines by GSK, is a liposome containing MPLA TLR4a and QS-21 saponin (*21*). Similarly, Irvine and coworkers recently designed an ISCOMATRIX analog incorporating MPLA TLR4a, named SMNP (*7*). Despite the potential for synergistic effects associated with combining saponins with a broader array of TLRas, saponin constructs incorporating a variety of TLRas haven’t yet been reported. Here, we report a library of saponin-based nanoparticle adjuvants incorporating various single or multiple clinically relevant TLRas (TLRa-SNPs) to generate a modular platform that could be broadly applicable. We generated four formulations of TLRa-SNPs incorporating Pam3CSK4 (TLR1/2a-SNP), MPLA (TLR4a-SNP), an imidazoquinoline derivative (TLR7/8a-SNP), as well as a mixture of both MPLA and the imidazoquinoline derivative (TLR4a-TLR7/8a-SNP). Compared to plain saponin nanoparticles (SNP), comparable to Matrix-M and ISCOMATRIX, all four TLRa-SNPs elicited improved humoral immune responses when used as adjuvants in SARS-CoV-2 and HIV vaccines in mice. TLRa-SNPs led to more potent antibody titers, improved antibody durability, enhanced breadth against variants of concern (VOCs), and induced better neutralizing responses when compared to SNPs or another clinically relevant control adjuvant CpG/Alum. Notably, TLRa-SNPs induced unique acute cytokine profiles, leading to distinct Th-skewing depending on the TLRa incorporated. Overall, we describe the generation of a potent and modular TLRa-saponin nanoparticle platform enabling tuning of Th-skewed responses, a pertinent feature in preventing diseases for which different Th-skewed responses lead to better clinical outcomes, as well as enhancing over humoral immune responses to vaccines.

## Results

### Formulation and characterization of TLRa-SNPs

We sought to create a platform of lipid-based saponin nanoparticles incorporating various TLRa adjuvants. The formulation of plain saponin nanoparticles (SNPs), traditionally known as ISCOMATRIX, has been widely reported in literature: mixing Quil-A saponin, dipalmitoylphosphatidylcholine (DPPC), and cholesterol at a molar ratio of 10:10:5 in aqueous medium triggers their spontaneous self-assembly into honeycomb-like nanoparticles 30-70 nm in diameter (*6, 8, 9*). We hypothesized that lipid-based TLRas could be readily incorporated with SNPs at a specific stoichiometry via hydrophobic effects, while non-lipidated TLRas could be conjugated with a cholesterol motif to specifically interact with saponins. In this way, we successfully incorporated TLR1/2a Pam3CSK4, TLR4a synthetic MPLA (MPLAs), and cholesteryl TLR7/8a imidazoquinoline derivative with SNPs by mixing the desired TLRa adjuvant, Quil-A saponin, DPPC, and total cholesterol content at a molar ratio of 1:10:2.5:10 across all formulations (Figure 1A, Table S1, Figure S1-3, and SI for synthesis). The resulting nanoparticles showed monodisperse populations of hydrodynamic diameters between 40 and 60 nm and negative surface charges equal to or below -30 mV (Figures 1B, 1C). Cryo-electron microscopy confirmed all nanoparticles conserved the signature honeycomb-like structure after introduction of TLRa adjuvants (Figure 1D). Moreover, TLRa-SNPs maintained their colloidal stability for at least 6 weeks when stored at 4 °C and for several months when stored at -20 °C (Figure S4).

**Figure 1:**
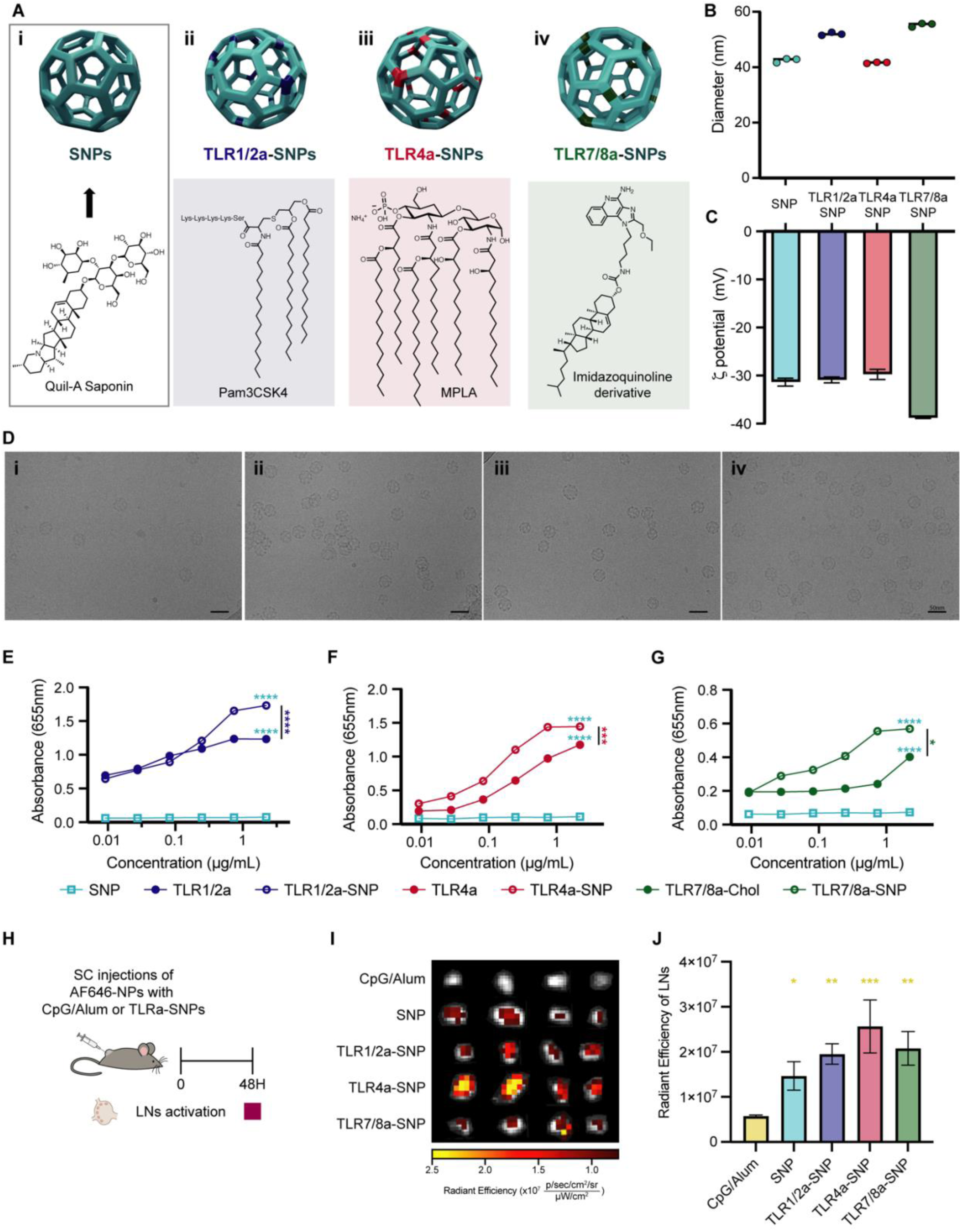
Design and characterization of TLRa-SNPs. **(A)** Schematic representation of SNP and different formulation of TLRa-SNPs: TLR1/2a-SNP incorporating Pam3CSK4, TLR4a-SNP incorporating MPLA, and TLR7/8a-SNP incorporating imidazoquinoline derivative. Hydrodynamic diameter **(B)** and surface charge **(C)** of SNP and TLRa-SNPs measured by Dynamic Light Scattering and Zetasizer. **(D)** Cryo-Electron Microscopy of **(i)** SNP, **(ii)** TLR1/2a-SNP, **(iii)** TLR4a-SNP, and **(iv)** TLR7/8a-SNP, demonstrating the maintenance of SNP structure after introduction of the TLRas. Scale bars: 50 nm. RAW-Blue macrophage cells incubated with saponin-nanoparticle (SNP) or TLRa incorporated SNPs (TLRa-SNPs). Activation curves of **(E)** soluble TLR1/2a and TLR1/2a-SNP, **(F)** soluble TLR4a and TLR4a-SNP, and **(G)** soluble cholesteryl-modified TLR7/8a and TLR7/8a-SNP across a range of TLRa concentrations (0.0091-2.22 μg/mL) with 100,000 RAW-Blue cells (n = 3). **(H)** Schematic of *in vivo* evaluation of antigen (AF647-NP) accumulation with different TLRa-SNPs and CpG/Alum control. **(I)** Fluorescence IVIS imagining of the inguinal draining lymph nodes 48h after subcutaneous injection at the tail base. **(J)** Quantification of the lymph node accumulations at 48h. Data were shown as mean +/- SEM. *p* values were determined with one-way ANOVA with Tukey test of the absorbance values at the highest TLRa concentration or logged radiant efficiency values. Complete *p* values for comparisons are shown in Table S3-4. \**p* < 0.05, \*\**p* < 0.01, \*\*\**p* < 0.001, and \*\*\*\**p* < 0.0001.

### In vitro activation of TLRa-SNPs and in vivo lymph node accumulation of NP antigen with TLRa-SNPs

To further confirm that TLRas had been incorporated with SNPs and maintained their bioactivities, we performed a RAW-Blue transgenic mouse macrophage *in vitro* cell assay to confirm TLR activation. RAW-Blue cells were incubated with different formulations of TLRa-SNPs (SNPs, TLR1/2a-SNPs, TLR4a-SNPs, or TLR7/8a-SNPs) or the corresponding soluble TLRa adjuvants. Activation of the TLRs by the TLRas would lead to downstream activation of NF-κB and AP-1 pathways, which can be assessed with QUANTI-Blue for a colorimetric output. Concentration-dependent NF-κB and AP-1 activation curves were generated by stimulating the cells with a range of soluble TLRa or TLRa-SNP concentrations (n=3; equivalent TLRas concentration of 2.22-0.0091 μg/mL; Figure 1E-1G, Figure S5, *p* values in Table S3). No activation with plain SNPs or relevant controls was observed. In contrast, all TLRa-SNPs generated concentration-dependent NF-κB and AP-1 activation. Notably, TLRa-SNPs produced more potent activation compared to their soluble counterparts, most likely due to the ability of the nanoparticle structure to multivalently display the TLRas and to enhance cellular uptake.

Since saponin-based adjuvants have been found to improve lymphatic drainage (*7*), we evaluated the accumulation of antigens in the lymph nodes with different TLRa-SNPs. Mice received 5 μg of an AlexaFluor 647-labelled I53-50 nanoparticle antigen (AF647-NP) and 10 μg of SNP or TLRa-SNPs via subcutaneous injections at the tail base (Figure 1H). I53-50 is a 28 nm-wide, 120-subunit complex that can be assembled *in vitro* by simply mixing trimeric and pentameric components forming a nanoparticle platform to allow for multivalent antigen presentation (*25–27*). This platform allows us to use AF647-labelled I53-50 nanoparticles for imagining and use the vaccine antigen-tethered I53-50 nanoparticles for immunogenicity studies. CpG/Alum was used as a clinically relevant positive control adjuvant. 48h after administration, the inguinal dLNs (n=4) were harvested for imaging using an In Vivo Imagining System (IVIS) (Figure 1I). Compared to CpG/Alum, vaccines comprising SNP or TLRa-SNPs elicited significantly higher accumulation of AF647-NPs in the dLNs (Figure 1J, *p* values in Table S4). Interestingly, TLRa-SNPs also resulted in higher accumulation compared to plain SNP, suggesting that the incorporation of TLRas improve antigen accumulation in the dLNs. Overall, the TLRa-SNP constructs maintained the bioactivity of the encapsulated TLRas *in vitro* and preserved the saponin’s ability to improve lymphatic drainage and lymph node accumulation *in vivo*.

### RBD-NP vaccines adjuvanted with TLRa-SNPs generate potent and broad humoral responses

The immune responses generated by different TLRa-SNPs were investigated using SARS-CoV-2 vaccines formulated with receptor-binding domain nanoparticles (RBD-NP) antigen and TLRa-SNPs as nanoparticle adjuvants. The RBD-NP used in this study, also called RBD-16GS-I53-50, has been extensively studied in mice, nonhuman primates (NHPs), and human clinical trials (*27–31*). It has been shown to generate potent neutralizing humoral responses when formulated with AS03 or CpG/Alum adjuvants, leading to licensure in several countries of RBD-NP adjuvanted with AS03 as a COVID-19 vaccine for adults 18 years or older (*32*). Following the same immunization regime as previous studies (*27, 30*), 7-8 week old C57BL/6 mice (n=5) were subcutaneously immunized with vaccines comprising 1.5 μg RBD-NP and 10 μg of TLRa-SNP formulations (SNP, or TLR1/2a-SNP, or TLR4a-SNP, or TLR7/8a-SNP). For comparison, a clinical control containing CpG/Alum (20 μg and 100 μg, respectively) was evaluated. Mice were immunized at Week 0 (prime) followed by a boost at Week 3, and sera were collected from Week 0 to Week 11 (Figure 2A).

**Figure 2:**
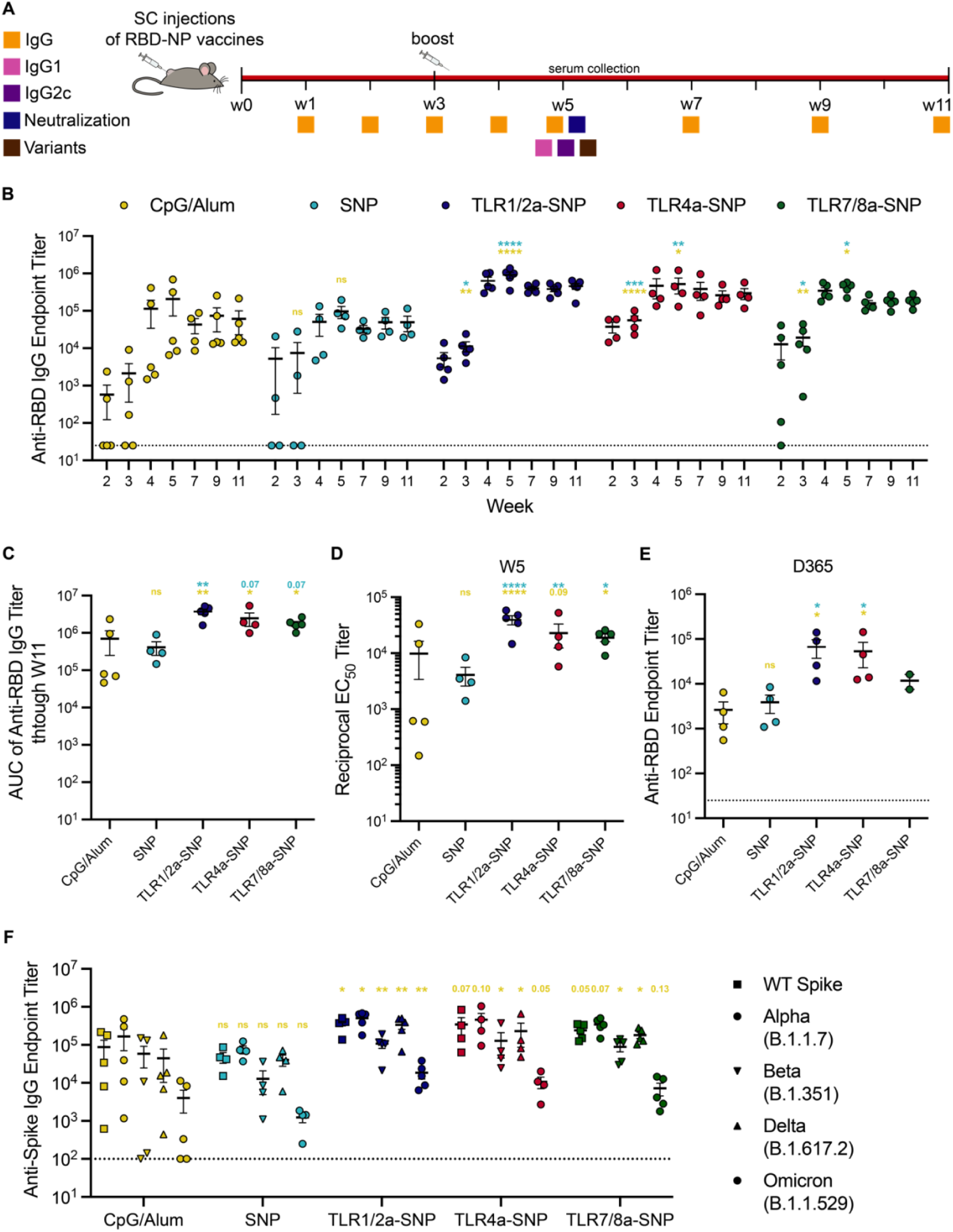
In vivo humoral response to RBD-NP vaccines adjuvanted with TLRa-SNPs. **(A)** Timeline of immunization and blood collection to determine IgG titers. Mice were immunized on Week 0 and boosted on Week 3 with RBD-NP vaccines adjuvanted with CpG/Alum, SNP, or TLRa-SNPs. IgG1, IgG2c, neutralization, and variants titers were determined on Week 5. **(B)** Anti-RBD IgG binding endpoint titers of RBD-NP vaccines adjuvanted with CpG/Alum, SNP, or TLRa-SNPs. **(C)** Area under the curves (AUCs) of anti-RBD IgG endpoint antibody titers from Week 0 to Week 11 of the different RBD-NP vaccines. **(D)** Half-maximal binding dilution (EC_50_) of anti-RBD IgG endpoint titers on Week 5 of the different RBD-NP vaccines. **(E)** Anti-RBD IgG binding endpoint titers a year (D365) after immunization. **(F)** Anti-spike IgG binding endpoint titers from sera collected on Week 5 after the initial immunization. Titers were determined for wildtype WT spike as well as Alpha (B.1.1.7), Beta (B.1.351), Delta (B.1.617.2), and Omicron (B.1.1.529) variants of the spike protein. Data (n = 4-5) are shown as mean +/- SEM. *p* values were determined using the general linear model followed by Tukey’s HSD comparison procedure on the logged titer values for IgG titer comparisons. Complete *p* values for comparisons are shown in Table S5- S9. \**p* < 0.05, \*\**p* < 0.01, \*\*\**p* < 0.001, and \*\*\*\**p* < 0.0001.

We first evaluated anti-RBD total IgG titers over time (Figure 2B). At Week 3, prior to boosting, mice immunized with RBD-NP vaccines adjuvanted with CpG/Alum or SNP elicited inconsistent antibody titers with around half the mice’s anti-RBD IgG endpoint below the limit of detection. In contrast, on Week 3, all mice immunized with vaccines adjuvanted with TLRa-SNPs (TLR1/2a-SNP, TLR4a-SNP, and TLR7/8a-SNP) seroconverted and generated antibody titers significantly higher than the CpG/Alum and SNP controls (*p* values in Table S5). Two weeks post-boost (Week 5), no significant difference in titers was observed between the two controls CpG/Alum and SNP, with an average endpoint titer of 2.1 x 10^5^ and 9.6 x 10^4^, respectively. However, TLRa-SNPs induced significantly higher titers for which the average endpoint titers were 9.1 x 10^5^ (TLR1/2a-SNP), 5.2 x 10^5^ (TLR4a-SNP), and 4.3 x 10^5^ (TLR7/8a-SNP). Additionally, TLRa-SNP vaccinated mice maintained significantly higher anti-RBD IgG endpoint titers at all timepoints post-boost and produced robust titers with little deviations across animals (*p* values in Table S5). In contrast, mice vaccinated with CpG/Alum resulted in variable titer responses spanning up to two orders of magnitude, leading to a standard error of mean (SEM) measured to be at least twice that of the rest of the treatments after boosting.

We calculated the area under the curves (AUCs) of the endpoint titers over the 11-week period to estimate the mean IgG production. No significant difference in AUC titers between CpG/Alum and SNP was observed (Figure 2C; *p* values in Table S6). While all TLRa-SNPs resulted in significantly higher AUCs than CpG/Alum, only TLR1/2a-SNP produced a significantly higher AUC than SNP. EC_50_ titers (half-maximal binding dilution) were also measured two weeks post-boost (Week 5) to ensure the endpoint titers reported correlated with the functional potency of the antibody (Figure 2D). Similar to comparisons of endpoint titers, CpG/Alum and SNP induced low and variable EC_50_ titers compared to TLRa-SNPs, which all generated significantly higher average EC_50_ (*p* values in Table S7). Furthermore, we assessed the endpoint titers of the vaccinated mice a year after immunization (D365, Figure 2E). While the average endpoint titers from Week 11 to D365 drastically dropped by 24-fold for CpG/Alum (2.6 x 10^3^) and 13-fold for SNP (3.9 x 10^3^), TLRa-SNPs’ endpoint titers remained high with only a 7-fold (6.7 x 10^4^) and 5-fold (5.4 x 10^4^) decrease for TLR1/2a-SNP and TLR4a-SNP, respectively. TLR7/8a-SNP vaccinated mice were excluded from the comparison due to an insufficient sample size that remained (n=2 at D365 due to murine dermatitis leading to euthanasia criteria; statistical power calculated using Mead’s Resource Equation) (*33, 34*). Overall, the titers measured at D365 from mice receiving vaccines adjuvanted with TLRa-SNPs were significantly higher than mice that received vaccines adjuvanted with either SNP or CpG/Alum controls. More importantly, these titers were at the same order of magnitude as the endpoint titers measured for the SNP and CpG/Alum controls on Week 11, demonstrating the durability of RBD-NP vaccines adjuvanted with TLRa-SNPs (Figure S6, *p* values in Table S8).

We further determined anti-Spike antibody titers to confirm the antibodies produced from RBD-NP adjuvanted with TLRa-SNP vaccines could bind to native spike proteins. Moreover, the breath of the antibody response was assessed by measuring the endpoint titers against different SARS-CoV-2 VOCs including Alpha (B.1.1.7), Beta (B.1.351), Delta (B.1.617.2), and Omicron (B.1.1.529) spike proteins two weeks post-boost (Week 5) compared to CpG/Alum (Figure 2F). While no differences in endpoint titers were observed between SNP and CpG/Alum regardless of variants (*p* values in Table S9), TLR1/2a-SNP led to significantly higher titers against all variants. Overall improved titers against variants were also observed for TLR4a-SNP and TLR7/8a-SNP. Notably, the ratio of titers against different variants compared to wild-type spike titer (percent drop) showed more consistent and higher breadth for TLRa-SNPs compared to CpG/Alum and SNP controls, for which more variable and larger interquartile ranges were observed (Figure S7). Overall, these findings suggested that SNP as an adjuvant did not perform better than the clinical control CpG/Alum in generating humoral responses. Only TLRa-containing SNPs elicited more potent and durable titer responses with better breadth.

### RBD-NP vaccines adjuvanted with TLRa-SNPs produce strong neutralizing antibodies

Since mice immunized with RBD-NP vaccines adjuvanted with TLRa-SNPs generated potent, durable, and broad antibody responses, we sought to measure the neutralizing activities of the sera. Week 5 sera neutralization was evaluated by utilizing a SARS-CoV-2 spike pseudotyped lentivirus to measure serum-mediated inhibition of viral entry into HeLa cells overexpressing ACE2 and TMPRSS2 (Figure 2A). Serum neutralization ID_50_ was measured through neutralizing activities of a range of sera concentrations to determine the half-maximal inhibition of infectivity (ID_50_) (Figure 3A-F). Consistent with previous findings(*31*), mice immunized with RBD-NP adjuvanted with CpG/Alum resulted in highly variable neutralizing responses, with two mice having serum neutralizing activity below the limit of detection. In contrast to CpG/Alum, RBD-NP vaccines adjuvanted with SNP and TLRa-SNPs induced significantly higher neutralizing ID_50_ (*p* values in Table S10). Notably, TLRa-SNPs elicited higher ID_50_ than the “high titer” classification according to FDA’s recommendation (ID_50_ ∼ 10^2.4^) (*35*). Moreover, mice sera neutralization activities were compared with human patients’ convalescent plasma by testing the pseudoviruses’ infectivity at the lowest sera dilution (1:100 dilution; Figure 3G). Variable results were observed for SNP and CpG/Alum controls, where not all mice receiving vaccines with these two adjuvants reached 0% infectivity. Mice immunized with vaccine adjuvanted with CpG/Alum resulted in significantly higher infectivity than those immunized with vaccines adjuvanted with TLRa-SNPs, with an average comparable to mice that received non-adjuvanted RBD-NP vaccine (*p* values in Table S11). On the contrary, we measured 0% infectivity from all mice receiving vaccines adjuvanted with TLRa-SNPs, indicating neutralization of all lentivirus and prevention of their entry into and infection of HeLa cells. Notably, the sera were even more neutralizing than the human patients’ convalescent plasma, for which the average infectivity was 15%. Overall, TLRa-SNPs as vaccine adjuvants generated superior humoral responses in potency, durability, breadth, and neutralization compared to both SNP and CpG/Alum controls.

**Figure 3:**
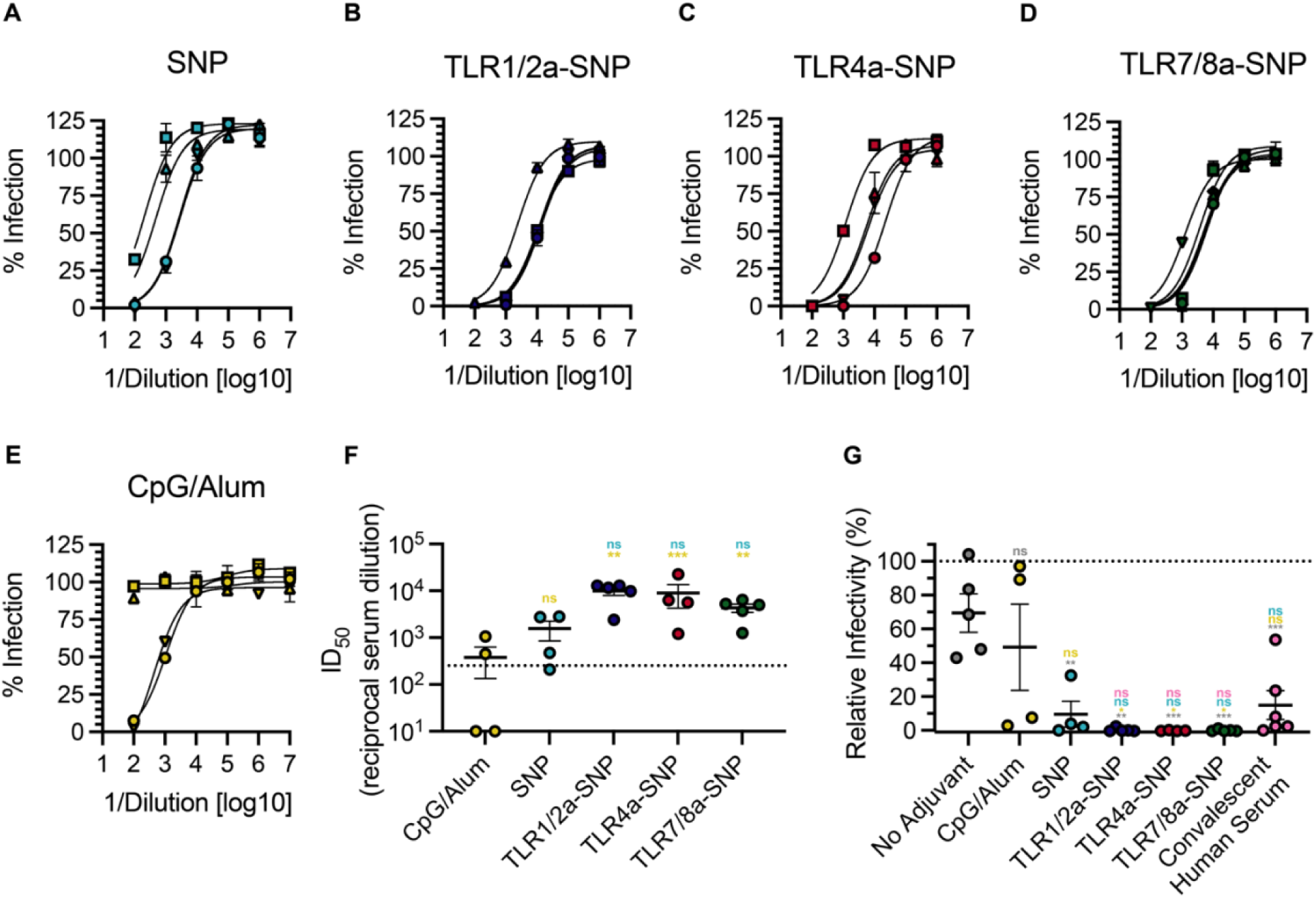
Neutralizing antibodies in mice post RBD-NP vaccines adjuvanted with TLRa-SNPs. **(A-E)** Percent infectivity for all vaccine treatments at a range of Week 5 serum dilutions as determined by a SARS-CoV-2 spike-pseudotyped viral neutralization assay. **(F)** Comparison of ID_50_ values determined from neutralization curves. Dotted line denotes the threshold for which the FDA considers as “high titer” (*35*). **(G)** Relative percent infectivity of all vaccine formulations compared to convalescent human serum at 1:100 dilution. Data (n = 4-5) are shown as mean +/- SEM. *p* values were determined using the general linear model followed by Tukey’s HSD multiple comparisons procedure on the logged neutralization ID_50_ values or relative infectivity comparisons. Complete *p* values for comparisons are shown in Table S10-S11. \**p* < 0.05, \*\**p* < 0.01, \*\*\**p* < 0.001, and \*\*\*\**p* < 0.0001.

### TLRa-SNPs elicit tunable Th-skewed responses

After assessing the potency of TLRa-SNPs in inducing durable and broad antibody responses, we further evaluated whether the nature of TLRa adjuvant affected the immune response of the TLRa-SNPs, since TLRas have been shown to generate unique immune signaling and T-helper responses (*36, 37*). We hypothesized that different TLRa-SNPs could elicit unique levels of IgG1 and IgG2c titers, as these isotypes are respectively strong indicators of Th2- and Th1-skewed responses. IgG isotypes were therefore measured across the different vaccine treatments. Indeed, while all TLRa-SNPs led to improved total IgG antibody responses compared to SNP and CpG/Alum controls, they generated different levels of IgG isotypes. While TLR4a-SNP and TLR7/8a-SNP generated similar IgG1 responses compared to SNP and CpG/Alum, TLR1/2a-SNP induced significantly higher IgG1 titers compared to all other groups, with average titers an order of magnitude higher compared to those generated by SNPs (Figure 4A, *p* values in Table S12). On the other hand, all TLRa-SNPs elicited higher IgG2c titers, with at least a 5-fold increase in average titers compared to the controls (Figure 4B, *p* values in Table S13). The differences in IgG1 and IgG2c titers suggested that different TLRa-SNPs resulted in different “flavors” of antibody responses, which we measured by calculating the ratio of IgG2c to IgG1 titers (Figure 4C). The ratio suggested TLR1/2a-SNP elicited a Th2-skewed response whereas TLR4a-SNP and TLR7/8a-SNP induced a Th1-skewed response. These results are drastically different from the balanced Th1/Th2 responses generated by SNP and CpG/Alum controls. While Th1-skewed responses may lead to better disease outcomes for SARS-CoV-2 infections (*38–41*), the tailored Th-responses induced from the different TLRa-SNPs could position them as a potent and modular adjuvant platform for inducing immunity against other challenging viruses by enabling the generation of bespoke adjuvant responses with distinct immune signatures.

**Figure 4:**
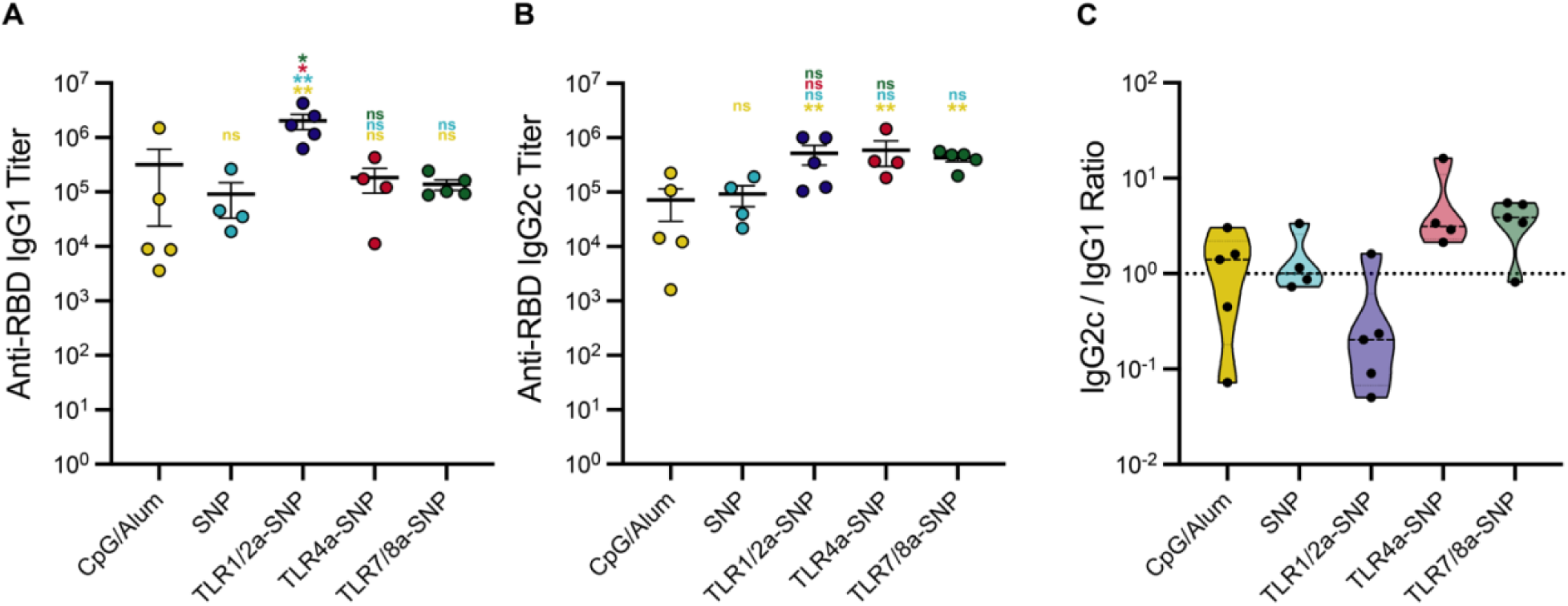
Antibody subtype response to RBD-NP vaccines adjuvanted with TLRa-SNPs. **(A)** Anti-RBD IgG1 and **(B)** IgG2c titers from sera collected on Week 5, 2 weeks after boost. **(C)** Ratio of Anti-RBD IgG2c to IgG1 titers. Lower values (below 1) suggest a Th2 response or humoral response, and higher values (above 1) imply a Th1 response or cellular response. Data (n = 4-5) are shown as mean +/- SEM. *p* values listed were using the general linear model followed by Tukey’s HSD multiple comparison procedure on the logged titer values. Complete p values for comparisons are shown in Table S12-S13. \**p* < 0.05, \*\**p* < 0.01, \*\*\**p* < 0.001, and \*\*\*\**p* < 0.0001.

### Different TLRa-SNPs induced unique acute cytokine induction in the draining lymph node and robust germinal center responses

We hypothesized that the differences in Th-skewed responses from different TLRa-SNP formulations were due to different TLRas leading to differences in immune signaling. To measure the acute immune response post-vaccination, we assessed dLN cytokine profiles 1-day (24 hours) post-immunization through a Luminex assay (Figure 5A). Instead of reporting the results in pg/mL, which can suffer from plate/batch/lot inconsistencies, nonspecific binding, differences between standards and real biological samples, and other experimental artifacts, we analyzed the raw fluorescence signal by correcting for nonspecific binding as a covariate in the regression analysis to better infer the true biological patterns in Luminex data (*42–45*). Sufficient levels of cytokines in the dLN are necessary to mount a successful vaccine response, and different cytokines induced can result in different Th responses. Comparing to mice adjuvanted with SNP, those that received TLRa-SNPs (TLR1/2a-SNP, TLR4a-SNP, or TLR7/8a-SNP) all induced significantly higher median fluorescence intensities (MFIs) of IFN-γ, IP10 (CXCL10), IL3, and IL15 (Figure 5B, *p* values in Table S14). These are critical proinflammatory cytokines for initiating strong immune responses and overall improved Th response (*46*). Elevated levels of IL6, IL25, and IL31 observed with TLR1/2a-SNP adjuvanted vaccine could explain the observed Th-2 skewed response. On the other hand, elevated levels of cytokines such as IFN-γ, IP10, IL18, IL27, and GMCSF observed with TLR4a-SNP could be the cause of the observed Th1-skewed response. Notably, of the 48 cytokines measured in the Luminex assay, only 11 did not show significant differences between SNP and at least one of the TLRa-SNPs (Figure S8). Thus, to better understand the overall cytokine induction profile of each adjuvant treatment, we performed a dimensional reduction analysis by creating a penalized supervised star plot (PSS) (Figure 5C). We were successful in generating distinct and separate cluster for each adjuvant. Not only did TLRa-SNPs induce higher effector cytokine production in the dLNs, but different formulations of TLRa-SNPs also led to unique acute cytokine profiles post-vaccination.

**Figure 5:**
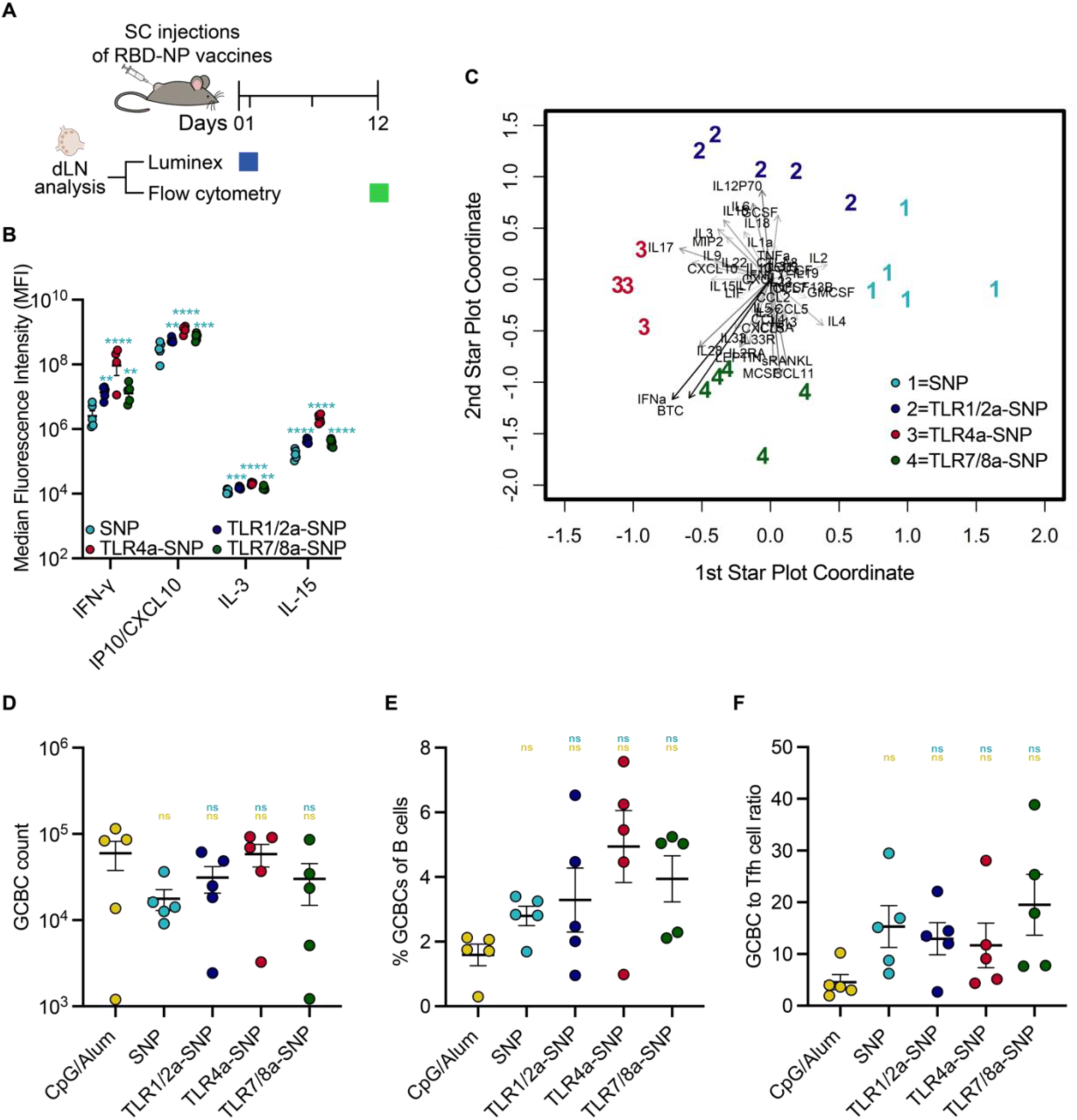
Draining lymph nodes (dLNs) analysis post-immunization with RBD-NP vaccines adjuvanted with SNP or TLRa-SNPs. **(A)** Timeline of immunization and dLN analysis. Luminex analysis was performed 1 day (24 hours) after vaccination, and flow cytometry was performed to measure germinal center B cells (GCBCs) and T-follicular helper cells (Tfhs) population 12 days after vaccination. **(B)** All three TLRa-SNPs induced significantly higher beneficial proinflammatory cytokine levels of IFN-γ, IP10, IL-3, and IL-15 in the dLNs compared to SNP, reported in median fluorescence intensity (MFI). **(C)** Dimensional reduction analysis of the full murine Luminex 48-plexed cytokine assay (Figure S8) by performing a penalized supervised star (PSS) plot of the RBD-NP vaccines. The result shows a clear separation of vaccines containing different adjuvants, with each number signifying an individual sample from the treatment groups. Vectors are the projection coefficients for each individual cytokine. **(D)** Total GCBC count and **(E)** frequency of GCBC from all B cells from RBD-NP vaccines. **(F)** GCBC to Tfh cell ratio. Data (n = 4-5) are shown as mean +/- SEM. *p* values for the Luminex assay were from a generalized maximum entropy estimation regression adjusted to control the false discovery rate and *p* values for flow cytometry were determined using the general linear model followed by Tukey’s HSD comparison procedure on the logged values. Complete *p* values for comparisons are shown in Table S14-S16 & S18.

We also assessed germinal center (GC) activity in the dLN 12 days after immunizing with RBD-NP adjuvanted with TLRa-SNPs using flow cytometry (Figure 5A, sample gating scheme in Figure S9). While mice treated with CpG/Alum adjuvanted vaccines had the highest count of germinal center B cells (GCBCs), they also had the lowest percent of GCBCs of total B cells, indicating CpG/Alum’s inability to convert B cells to GCBCs (Figure 5D-E, *p* values in Table S15-S16). On the contrary, mice immunized with vaccines adjuvanted with TLRa-SNPs resulted in higher percentages of GCBCs as well as higher total GCBC counts compared to mice receiving vaccines adjuvanted with SNP, suggesting robust GC activities. Moreover, compared to TLRa-SNP groups, mice receiving vaccines adjuvanted with SNP resulted in the lowest count of T-follicular helper cells (Tfhs) (Figure S10, *p* values in Table S17). We then assessed the ratio of GCBCs to Tfhs since this ratio has been found to correlate with the quality of T cell help (*47–49*). Interestingly, SNP and TLRa-SNP groups all resulted in a higher GCBC:Tfh ratio compared to the CpG/Alum group (Figure 5F, *p* values in Table S18). The overall improvements in Tfh count and the higher ratio of GCBC:Tfh suggest that TLRa-SNPs can improve the quality of T cell help compared to SNP and CpG/Alum adjuvants. Since robust GC activity is important for generating better neutralizing and higher affinity antibodies, these data are consistent with the vaccination results that demonstrated increased magnitude and higher neutralizing antibodies with the TLRa-SNP groups. These experiments suggest TLRa-SNPs are superior in generating unique early proinflammatory responses and robust GC activity compared to SNP and CpG/Alum, resulting in an overall improved potency, durability, and breadth of humoral responses.

### Saponin nanoparticle incorporating both TLR4a and TLR7/8a generated synergistic immune stimulation

Prior research has shown that simultaneously triggering TRIF-coupled TLR and endosomal TLR can synergistically activate the immune system (*5, 50, 51*). As such, activating both TRIF-coupled TLR4 and endosomal TLR7/8 has been found to increase antigen-specific, neutralizing antibodies by enhancing the persistence of GCs and plasma-cell responses compared to activating single TLR ligands (*50*). We, therefore, created a new formulation of TLRa-SNP, TLR4a-TLR7/8a-SNP, in which we incorporated equimolar amounts of TLR4a MPLA and cholesteryl TLR7/8a imidazoquinoline derivative in SNPs (Figure 6A). The total molar ratio of TLRa to saponin was kept identical to the previously described TLRa-SNP formulations (Table S1). The average hydrodynamic diameter of TLR4a-TLR7/8a-SNP, 47.9 nm, is consistent to the diameters measured from SNPs and other TLRa-SNPs (Figure 1B, Figure S11A-B). Similar to the other TLRa-SNPs evaluated, TLR4a-TLR7/8a-SNP significantly enhanced AF647-NP accumulation in the dLNs in mice compared to CpG/Alum (Figure S11C-E, *p* values in Table S4). To confirm whether co-incorporating TLR4a and TLR7/8a in the same SNP is synergistic, we immunized C57BL/6 mice (n=5) with vaccines formulated with 1.5 μg of RBD-NP adjuvanted with either 10 μg of TLR4a-TLR7/8a-SNP or a 1:1 mixture of TLR4a-SNP and TLR7/8a-SNP (5 μg each, referred as TLR4a-SNP + TLR7/8a-SNP). Mice were vaccinated on Week 0 and boosted on Week 3 matching the previous schedule. Binding titers were measured to compare with those from SNP, TLR4a-SNP, and TLR7/8a-SNP formulations (Figure 6B). We observed no significant difference of anti-RBD total IgG titers over time in mice adjuvanted with TLR4a-SNP + TLR7/8a-SNP compared to mice adjuvanted with either TLR4a-SNP or TLR7/8a-SNP (Figure 6C, *p* values in Table S19). In contrast, mice adjuvanted with TLR4a-TLR7/8a-SNP resulted in overall higher titers than all other groups, with a significant increase observed after the boost. The co-incorporation of both TLR4a and TLR7/8a on the same SNP is therefore pivotal for their synergistic effect. Consistent with other TLRa-SNP formulations, mice receiving vaccines adjuvanted with either TLR4a-SNP + TLR7/8a SNPs or TLR4a-TLR7/8a-SNP maintained robust binding titers against wild-type spike and different SARS-CoV-2 VOCs (Figure S11F-G). Finally, both the TLR4a-SNP + TLR7/8a-SNP and TLR4a-TLR7/8a-SNP formulations maintained similar IgG1 and IgG2c titers compared to mice adjuvanted with either TLR4a-SNP or TLR7/8a-SNP, resulting in overall equally Th1-skewed responses (Figure 6D-F, *p* values in Table S20-S21).

**Figure 6:**
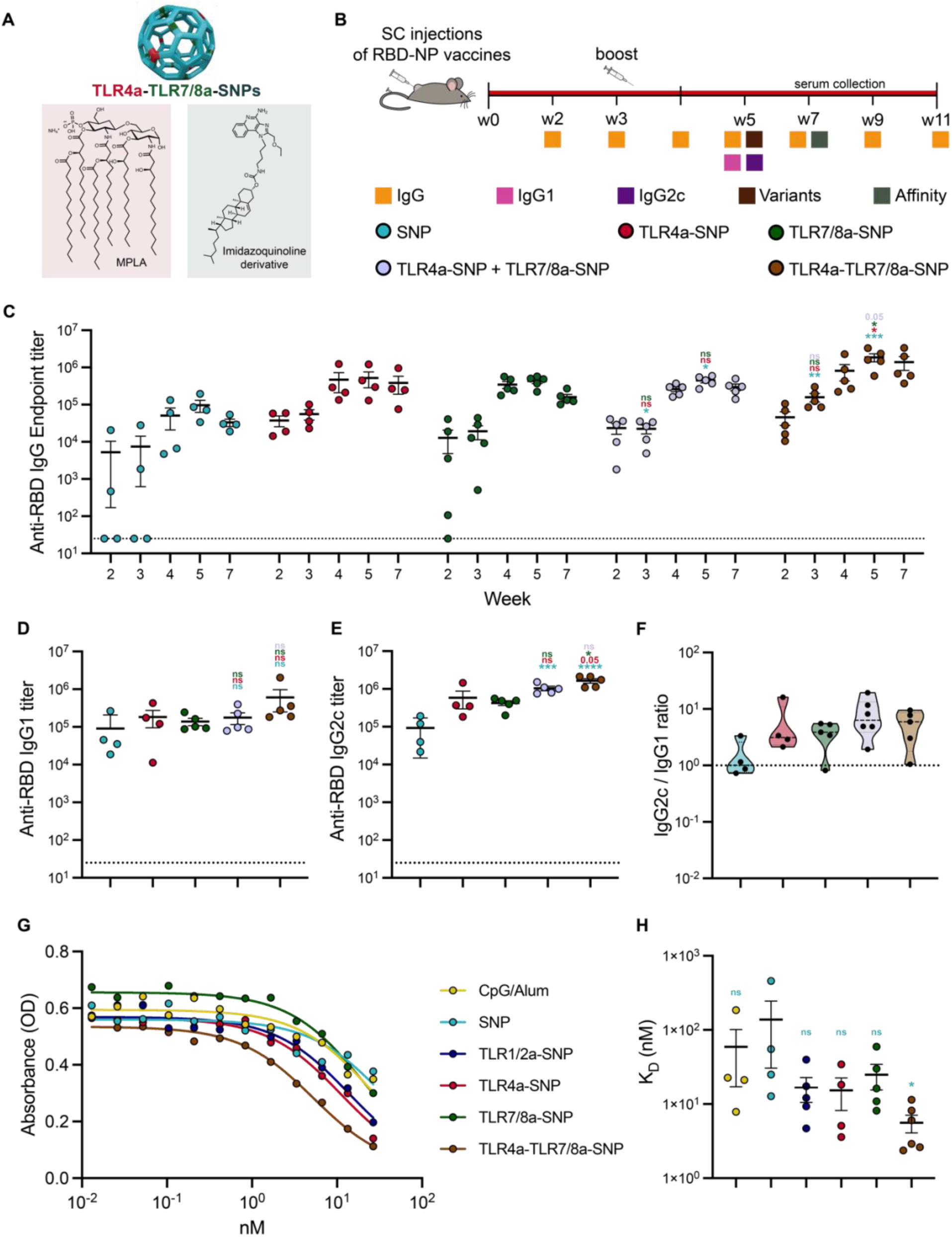
TLRa-SNP incorporating both TLR4a and TLR7/8a generated synergistic effect. **(A)** Schematic representation of TLR4a-TLR7/8a-SNP for which SNPs co-incorporate TLR4a MPLA and TLR7/8a imidazoquinoline derivative. **(B)** Timeline of RBD-NP vaccines immunization and blood collection to determine IgG titers. Mice were immunized on Week 0 and boosted on Week 3 with RBD-NP vaccines adjuvanted with a 1:1 mixture of TLR4a-SNP and TLR7/8a-SNP or TLR4a-TLR7/8a-SNP. IgG1, IgG2c, and variants titers were measured on Week 5. **(C)** Anti-RBD IgG binding endpoint titers of RBD-NP vaccines. **(D**) IgG1 titers and **(E)** IgG2c titers. **(F)** Ratio of Anti-RBD IgG2c to IgG1 titers. Lower values (below 1) suggest a Th2 response or humoral response, and higher values (above 1) imply a Th1 response or cellular response. **(G)** Week 7 competitive binding curves for different RBD-NP vaccines compared to an mAb reference competing with the same mAb. **(H)** Calculated K_D_ values from fitted binding curves. Data (n = 4-5) are shown as mean +/- SEM. *p* values were determined using the general linear model followed by Tukey’s HSD comparison procedure on the logged titer values for IgG titer comparisons. Complete *p* values for comparisons are shown in Table S19-S22. \**p* < 0.05, \*\**p* < 0.01, \*\*\**p* < 0.001, and \*\*\*\**p* < 0.0001.

To determine whether RBD-NP vaccines adjuvanted with TLRa-SNPs resulted in improved antibody quality, we investigated the post-boost serum antibody affinities toward RBD using a competitive binding ELISA (Figure 6G-H). The polyclonal populations of antibodies produced by mice that received vaccines adjuvanted with SNP and CpG/Alum were found to have the highest K_D_ with values of 134 and 60 nM, respectively. All antibodies from mice that received vaccines adjuvanted with TLRa-SNPs exhibit K_D_ values almost an order of magnitude lower than the control vaccine with SNP, with the TLR4a-TLR7/8a-SNP group eliciting a significantly lower K_D_ of 5.5 nM (*p* values in Table S22). These results are consistent with the findings that TLRa-SNPs enhance GC responses as well as improve antibody neutralization.

### TLRa-SNPs as gp120 vaccine adjuvants induce robust humoral responses

Finally, to confirm that TLRa-SNPs can elicit robust immune responses against other antigens, we vaccinated C57BL/6 mice (n=5) with HIV gp120 antigen adjuvanted with TLR1/2a-SNP, TLR4a-SNP, TLR7/8a-SNP, and TLR4a-TLR7/8a-SNP. Titer responses were compared to the two following controls, gp120 vaccines adjuvanted with SNP or Alum. Mice were immunized on Week 0 and boosted on Week 4 and sera were collected from Week 0 to Week 6 to assess total anti-gp120 IgG and subtype titers (Figure 7A). Before boosting, most mice that received Alum or SNP adjuvanted vaccines did not have detectable titers, but all mice that received vaccines adjuvanted with TLRa-SNPs seroconverted and elicited significantly higher titers (Figure 7B, *p* values in Table S23). Two weeks post-boost (Week 6), vaccines containing TLRa-SNP adjuvants generated significantly higher antibody titers compared to those with Alum, with average endpoint titers around 2000-fold higher. Notably, mice adjuvanted with TLR4a-TLR7/8a-SNP elicited the strongest response. Moreover, TLRa-SNPs also led to over 3-fold higher titers than SNP. Similarly, we observed significantly higher overall antibody production from mice adjuvanted with TLRa-SNPs compared to the controls by determining the AUCs of titers over the 6-week period (Figure 7C, *p* values in Table S24).

**Figure 7:**
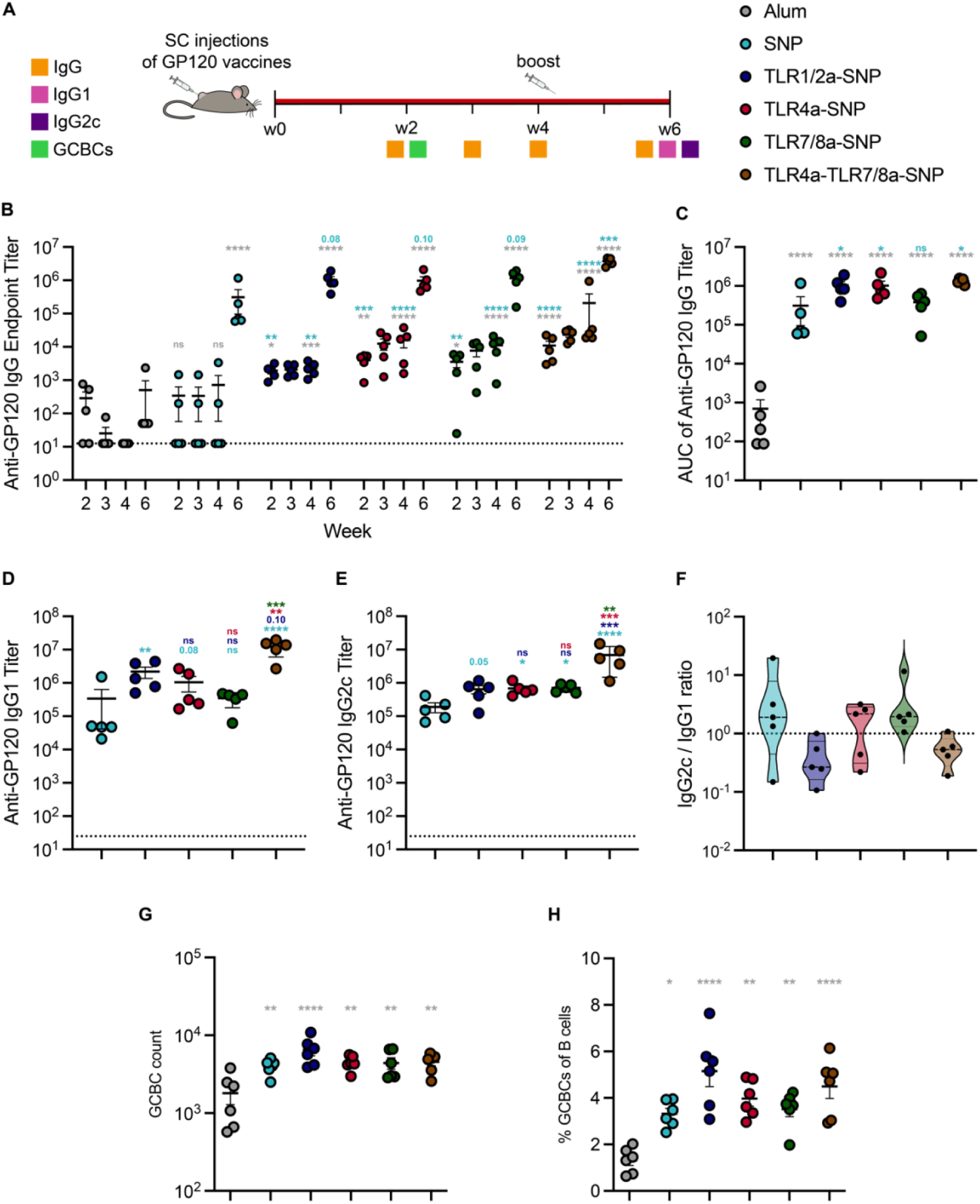
In vivo humoral response to HIV gp120 vaccines adjuvanted with TLRa-SNPs. **(A)** Timeline of immunization and blood collection to measure IgG titers. Mice were immunized on Week 0 and boosted on Week 4 with gp120 vaccines adjuvanted with Alum, SNP, or TLRa-SNPs. IgG1 and IgG2c titers were determined on Week 6. **(B)** Anti-gp120 IgG binding endpoint titers of gp120 vaccines adjuvanted with Alum, SNP, or TLRa-SNPs. **(C)** Area under the curves (AUCs) of anti-gp120 IgG endpoint antibody titers from Week 0 to Week 6 of different gp120 vaccines. **(D)** Anti-gp120 IgG1 and **(E)** IgG2c titers from sera collected on Week 6, 2 weeks after boost. **(F)** Ratio of Anti-gp120 IgG2c to IgG1 titers. Lower values (below 1) suggest a Th2 response or humoral response, and higher values (above 1) imply a Th1 response or cellular response. **(G)** Total GCBC count from gp120 vaccines and **(H)** Frequency of GCBC from all B cells. Data (n = 5-6) are shown as mean +/- SEM. *p* values were determined using the general linear model followed by Tukey’s HSD comparison procedure. Complete *p* values for comparisons are shown in Table S23-S28. \**p* < 0.05, \*\**p* < 0.01, \*\*\**p* < 0.001, and \*\*\*\**p* < 0.0001.

To confirm that the unique Th-skewed responses observed from different TLRa-SNPs in RBD-NP vaccines were due to influences of TLRas in immune signaling, we measured Week 6 IgG1 and IgG2c titers from mice immunized with gp120 vaccines. Consistent with our previous observations, mice adjuvanted with TLR1/2a-SNP, TLR4a-SNP, and TLR7/8a-SNP all induced robust IgG2c responses while only TLR1/2a-SNP generated significantly higher IgG1 titers compared to SNP (Figure 7D-E, *p* values in Table S25-26). On the other hand, TLR4a-TLR7/8a-SNP elicited significantly higher IgG1 and IgG2c titers compared to SNP. This resulted in a balanced Th1/Th2 response from mice adjuvanted with TLR4a-TLR7/8a-SNP, a more Th2-skewed response from mice adjuvanted with TLR1/2a-SNP, and a more Th1-skewed response from mice adjuvanted with TLR4a-SNP and TLR7/8a-SNP (Figure 7F). Additionally, we assessed GC activity in the dLN 14 days after immunizing with gp120 adjuvanted with TLRa-SNPs (Figure 7A). Mice immunized with vaccines adjuvanted with TLRa-SNPs resulted in overall higher count and percent of GCBCs compared to mice adjuvanted with SNP or Alum (Figure 7G-H, *p* values in Table S27-28).

Finally, we immunized male C57BL/6 mice with gp120 vaccines adjuvanted with either SNP or TLRa-SNPs to confirm TLRa-SNPs have similar potent vaccine adjuvanting effects in both sexes (Figure S12A). We observed similar antibody titers between the male and female mice after being vaccinated with gp120 vaccines adjuvanted with TLRa-SNPs (Figure S12B-C). Consistent with previous findings in female mice, mice adjuvanted with TLR1/2a-SNP, TLR4a-SNP, TLR7/8a-SNP, or TLR4a-TLR7/8a-SNP all elicited significantly higher titers both pre- and post-boost compared to the SNP control (Figure S12B-C, *p* values in Table S29). All together, these results confirmed TLRa-SNPs’ ability to generate potent and robust antibody responses with distinctive Th-skewed responses regardless of the antigen or gender of the animal.

## Discussion

Over the past few decades, modern vaccinology has moved towards the use of subunit protein-based vaccines over more traditional vaccine formats on account of their increased affordability, ease of manufacturing, and high safety profile (*2*). Unfortunately, the poor immunogenicity of many protein antigens reinforces the need for developing more potent and modular adjuvants. In this study, we designed an adjuvant platform combining three approaches in adjuvant design and selection: (i) molecular TLRa adjuvants resembling pathogen-associated molecular patterns found on bacteria, viruses, and other foreign invaders our immune system is trained to recognize; (ii) saponin adjuvants from natural sources that improve lymphatic flow; and (iii) particulate design displaying both adjuvants together with improved trafficking kinetics and cellular uptake. Previous studies have reported the design of combined TLR4a MPLA and saponins into nanoparticle constructs, such as AS01 and SMNP. Nonetheless, a tunable platform approach must consider incorporating other clinically relevant TLRas. In this regard, we demonstrated that lipid TLRas, such as MPLA and Pam3CSK4, could be readily incorporated with clinically relevant stoichiometric ratios due to the hydrophobic nature of SNPs. Non-lipid TLRas, such as TLR7/8a Resiquimod, could be incorporated by conjugating a cholesterol motif that hydrophobically interacts with SNPs. The facile assembly and modularity of the SNP platform also allow the design and investigation of SNPs containing multiple TLRas in a same particle such as TLR4a and TLR7/8a (TLR4a-TLR7/8a-SNP). In this regard, consistent with previous reports suggesting TLR4a and TLR7/8a synergistically activate the innate immune system (*5, 50*), we observed improved antibody responses from mice receiving vaccines adjuvanted with TLR4a-TLR7/8a-SNP. While prior research supported synergistic responses by activating both TRIF-coupled TLR4 and endosomal TLR7/8, examining TLRa-SNPs combining TLR1/2a and TLR4a or TLR1/2a with TLR7/8a may be of interest for future studies.

A recent report by Irvine and coworkers provided detailed mechanistic analyses of the improved humoral response induced by ISCOMATRIX and SMNP, similar to SNP and TLR4a-SNP in the present study (*7*). Mainly, both adjuvants enhance lymph flow in a mast cell-dependent manner, which promotes antigen delivery into the dLNs. Since this mechanism was determined to be saponin-dependent, we assumed all TLRa-SNPs improved vaccine immune responses in a similar manner. Further, since mast cells were shown to be essential in saponins’ ability to improve lymph flow and are effector cells of Th-2 response, we hypothesized that TLR1/2a-SNP conferred superior humoral responses due to its ability to generate a Th-2 skewed response. Future investigation could determine other synergistic effects of different TLRas with SNPs. Moreover, no significant elevation in proinflammatory cytokines or chemokines in the dLNs was observed when mice received soluble MPLA mixed with ISCOMATRIX compared to mice receiving ISCOMATRIX alone, suggesting it is necessary to deliver both adjuvants in the same particle for their synergistic effects. In addition, previous reports have also suggested that delivering soluble TLRas can lead to potentially toxic levels of systemic inflammation due to their rapid clearance into the bloodstream (*7, 52, 53*). Hence, delivering soluble TLRas without carriers such as Alum or various nanoparticle constructs poses safety risks that may hinder clinical translation (*54*). As such, soluble TLRas mixed with SNP matching TLRa-SNP formulations were not tested as controls in this study. Furthermore, Irvine and coworkers reported no signs of toxicity of their SMNP design. The present study includes the use of clinically safe TLRas dosed in mice in lower masses than previously reported (*7, 31, 55, 56*). We therefore assumed the TLRa-SNPs were non-toxic as well, an assumption supported by the absence of any toxicity signs in mice such as weight loss or injection site reactogenicity.

Numerous preclinical and clinical studies have utilized the RBD-NP antigen RBD-16GS-I53-50 developed by King and coworkers (*27–30*). In this study, we used the same RBD-NP antigen in a dose- and schedule-matched study design to allow for comparison of the potency of TLRa-SNPs with previously screened adjuvants (*27–30*). From those reports, AS03 and CpG/Alum have been reported to induce comparable humoral responses, including robust neutralizing responses, and both outperformed other clinically relevant adjuvants such as AS37, AddaVax, and Essai O/W 1849101 (*30*). The use of CpG/Alum as a control in the present study therefore highlights the superiority of all TLRa-SNPs (i.e., TLR1/2a-SNP, TLR4a-SNP, TLR7/8a-SNP, and TLR4a-TLR7/8a-SNP) in generating potent, durable, broad, and neutralizing antibody responses (*30*). This observation is even more striking considering that the dosage of CpG/Alum was doubled compared to that used in literature (*30*). Additionally, mice receiving vaccines adjuvanted with TLRa-SNPs demonstrated complete neutralization in pseudovirion neutralization assays, in contrast to convalescent human plasma which resulted in a quantifiable level of infectivity. This observation is particularly striking considering all human patients had previously received up-to-date COVID-19 vaccines, including an original prime and boost regimen plus an additional booster of either Moderna or Pfizer COVID-19 mRNA vaccines prior to contracting COVID-19 only 8-12 weeks before sample collection. Previous reports have demonstrated that patients with such hybrid immunity have superior neutralizing antibodies than patients who had been infected but not previously vaccinated (*57–61*). These data show great promise for further investigation of the TLRa-SNPs adjuvant technology in larger animal models. This would be especially of interest in assessing neutralizing antibody responses post-vaccination with HIV vaccines adjuvanted with TLRa-SNPs since there are generally no detectable neutralizing antibodies towards HIV in mice.

Additionally, in an NHP study, treatment groups receiving AS03-adjuvanted RBD-NP antigen vaccines had extended antibody titers and more durable Omicron variant protection when compared to values previously reported for commercial Pfizer-BioNTech and Moderna mRNA vaccines (*28*). Nonetheless, antibody titers in NHPs waned to pre-boost magnitudes by 6 months post-boost, necessitating additional boosters to maintain robust protection (*28*). We showed that mice receiving RBD-NP vaccines adjuvanted with TLRa-SNPs maintained antibody endpoint titers for an entire fpage following that were of similar magnitude to animals receiving vaccines adjuvanted with SNP or CpG/Alum at Week 11 following vaccination. These results are promising for future studies on larger animal models such as NHPs. These astonishing results were further emphasized by the rapid seroconversion of all mice after a single immunization (i.e., the prime immunization), thereby demonstrating the potential of single-immunization vaccines adjuvanted with these TLRa-SNPs. The rapid and complete seroconversion was further verified with HIV gp120 protein antigen, suggesting the superiority and potential broad application of TLRa-SNPs compared to SNP and clinically relevant control adjuvants. Notably, we did not observe any substantial differences between the two clinical controls, CpG/Alum (mimicking Dynanvax’s CpG1018) and SNP (mimicking Novavax’s Matrix M). Although the adjuvant potencies remained similar for these two control adjuvants, the differences in their immune-stimulating mechanism might create synergistic effects when combined, illustrated by the potent, durable, broad, and neutralizing humoral responses induced by TLRa-SNPs, which take advantage of multiple immune-stimulating mechanisms. Overall, TLRa-SNPs adjuvants can rapidly lead to protective levels of neutralizing antibodies and prolong the durability of neutralizing responses, therefore strongly decreasing the need for costly booster shots, which is crucial for vaccines capable of rapidly and broadly protecting worldwide populations during a pandemic.

The unique acute cytokine induction profiles and Th-skewed responses observed in this study for each TLRa-SNP formulation are consistent with prior literature (*3, 5, 62*). TLR1/2a-SNP, TLR4a-SNP, and TLR7/8a-SNP generated robust IgG2c titers, thereby suggesting a strong Th1 response. Indeed, all three TLRas can induce Th1 response by activating the NF-κB pathway. Elevated levels of IL-12 and IL1β, hallmark cytokines of NF-κB activation, in the Luminex assay further verify the incorporation of these three TLRas and their impact on immune signaling (*3*). Additionally, only TLR1/2a-SNP elicited higher IgG1 titer, indicating an overall Th2-skewed response. The activation of TLR1 and TLR2 triggers the ERK1 and ERK2 pathways, which could result in higher Th2 responses (*3*). Similarly, the initiation of endosomal TLRs such as TLR7 and TLR8 activates the IRF-pathway, which in turn induces strong type I interferon response such as the secretion of IFN-α and IFN-β. The PSS plot clearly indicates a stronger expression of IFN-α (demonstrated by the vector) in the TLR7/8a-SNP cluster (cluster 4, Figure 5C). While the Luminex panel did not include IFN-β, we nonetheless observed other strong interferon responses, such as high IFN-γ, IP10, and IL28 production from mice adjuvanted with TLR7/8a-SNP. The overall balanced but slightly Th-1 skewed response is consistent with prior findings (*3, 53, 62*). Likewise, the Th1-skewed response observed with TLR4a-SNP is consistent with the previously reported SMNP, for which only IgG2a titers, but not IgG1, were significantly higher than the SNP control (*7*). Overall, the Luminex assay helped to further verify the success of TLRas incorporation to the SNP as well as the unique Th responses induced from the library of TLRa-containing saponin-based nanoparticles. In this regard, literature reports have shown that while Th2-skewed responses resulted in more favorable clinical outcomes for some infectious diseases such as Rabies virus (*63, 64*), Th1-skewed responses were favorable for diseases such as COVID-19 (*38–41*). Thus, having a library of TLRa-SNPs with tailored Th-skewed responses may have profound clinical implications for which a specific adjuvant formulation could be selected during the early stage of vaccine development depending on the disease the vaccine is designed to prevent.

Additional characterization of the TLRa-SNPs, including evaluation of how these nanoparticles could influence antigen stability and conformation when combined, could be of interest to further extend the broad use of this nanoparticle platform. Moreover, protein-based vaccines have been previously reported to not induce strong cellular immunity. RBD-NP, the primary antigen used in this study, was found to not generate a significant CD8+ T cell response (*29*). We observed comparable results when assessing antigen-specific IFN-γ+CD8+ T cells by ELISpot following activation of splenocytes two weeks post-boost with pan-RBD peptides (Figure S13, *p* values in Table S30). While most clinical vaccines focus on inducing humoral responses (*65*), further insights on TLRa-SNPs’ ability to generate cellular responses could make them promising candidates as T cell or cancer vaccines. In addition, other than generating different Th-skewed responses, different TLRas have also been found to induce differences in other immune cell populations, such as different levels of long-lived plasma cells or memory cells. Studying the activation and induction of innate cells and effector cells by different TLRa-SNPs could be of interest for developing a comprehensive understanding of the immune-stimulating nature of these adjuvant formulations.

Following our efforts to develop broadly applicable vaccine technologies, the newly designed toll-like receptor agonist containing saponin-nanoparticles (TLRa-SNPs) showed great promise in improving the potency, durability, breadth, and neutralization of vaccines with the potential of preventing infection by complex immune evasive viruses such as SARS-CoV-2 and HIV. Notably, the different adjuvant formulations induced unique acute cytokine and immune-signaling profiles, leading to different Th-responses. This robust and tunable adjuvant library reinforces the global effort of the Coalition for Epidemic Preparedness Innovations and the World Health Organization to develop and manufacture vaccines within 100 days in response to “Disease X,” an infectious agent currently unknown to cause human disease.

## Materials and Methods

### Materials

Cholesterol (ovine wool, > 98%) and 1,2-dipalmitoyl-*sn*-glycero-3-phosphocholine (DPPC) were purchased from Avanti Polar Lipids. Quil-A adjuvant, Monophosphoryl Lipid A (from *S. minnesota R595*, MPLA-SM VacciGrade) and Pam3CSK4 (VacciGrade) were purchased from Invivogen. Cholesteryl imidazoquinoline derivative was synthetized in a one-step reaction from 4-amino-2-(ethoxy methyl)-1H-imidazo[4, 5-C]quinoline-1-butylamine (Career Henan Chemical Co., Henan Province, China; see supporting information). SARS-CoV-2 proteins and antibodies were purchased from Sino Biological including SARS-CoV-2 RBD protein (40592-V08H), SARS-CoV-2 spike protein (40589-V08H4), Alpha B.1.1.7 spike (40591-V08H10), Beta B.1.351 spike (40591-V08H12), Delta B.1.617.2 spike (40591-V08H23), Omicron B.1.1.529 spike (Sino Biological 40591-V08H41), SARS CoV-2 spike RBD monoclonal antibody (40150-D001), and SARS CoV-2 spike RBD monoclonal antibody with HRP (40150-D001-H). HIV antigen gp120 was purchased from Immune Technology Corporation (IT-001-022p). CpG1826 (Vac-1826) and Alum (Alhydrogel 2%, vac-alu) were purchased from Invivogen. Maxisorp plates were purchased from Thermofisher. Goat anti-mouse IgG Fc secondary antibody (A16084) HRP (Horseradish peroxidase) was purchased from Invitrogen. Goat anti-mouse IgG1 and IgG2c Fc secondary antibodies with HRP (ab97250, ab97255) and 3,3’,5,5’-Tetramethylbenzidine (TMB) high sensitivity ELISA Substrate were purchased from Abcam. Single color mouse IFN-γ ELISPOT kit was purchased from Cellular Technology Limited. Antigen-specific cell stimulating peptides (Pan-SARS-CoV-2 S-RBD peptides) were purchased from JPT Peptide Technologies (PM-WCPV-S-RBD-1).

### Formulation of SNPs and TLRa-SNPs

The formulation of lipid-based nanoparticles has been previously reported (*6–11*). Briefly, cholesterol and DPPC (20 mg/mL) were each dissolved in a solution of 20% (w/v) of MEGA-10 detergent in Milli-Q water. Quil-A saponin (20 mg/mL) was dissolved in pure Milli-Q water. All solutions were heated at 50 °C until complete dissolution before use. When required, Cholesteryl imidazoquinoline, MPLA, and Pam3CSK4 (2 mg/mL) were each dissolved in a solution of 20% (w/v) of MEGA-10 detergent in Milli-Q water and heated at 70 °C under sonication until dissolution. All formulations were obtained after quickly mixing the appropriate volume of each compound at 60 °C without stirring in this order: Cholesterol/DPPC/Adjuvant/Quil-A saponin followed by its dilution with PBS 1X to reach a final concentration of 1 mg/mL in cholesterol (Table S1). Solutions were left for equilibration overnight at room temperature and dialyzed against PBS 1X (MWCO 10 kDa) for 5 days. Solutions were then filtered using 0.2 μm Acrodisc® Syringe Filters and concentrated using Centricon spin filters (MWCO 50 kDa, 3000 RCF, 40 mins) in stock solutions of 5 mg/ml in Quil-A Saponin.

### Dynamic Light Scattering and Zeta Potential

The hydrodynamic diameter and surface charge of the NPs were respectively measured on a DynaPro II plate reader (Wyatt Technology) and a Zetasizer Nano Zs (Malvern Instruments). Three independent measurements were performed for each sample.

### CryoEM and cryoET sample preparation and data collection

CryoEM and cryoET samples were prepared using Vitrobot Mark IV. First, 1.2/1.3 Quantifoil grids (Ted Pella inc., 658-200-CU-100) were glow discharged using PELCO easiGlow™ for 30 s. For cryoET, 6 nm gold fiducial was mixed with the samples in a 1:10 volume ratio. Then, 2 mg/mL sample was applied to the glow discharged gird (Electron Microscopy Sciences, 25486), followed by removing the extra sample amount using filter paper and quickly plunging into liquid ethane. CryoEM and cryoET data were acquired by Titan Krios G3 electron microscope equipped with a K3 direct detector and energy filter with an accelerated voltage of 300 kV using SerialEM (*66*). CryoEM micrographs were collected on a pixel size of 1.13A/pixel with and -2 μm defocus. CryoET data was collected from -60 °C to +60 °C with a 2 °C increment. The total electron dose was 80 e^-^/A^2^ with -3 μm defocus, and the pixel size was 1.13A. Reconstruction was done by IMOD (*67*) and visualized by chimera (*68*).

### In vitro RAW-Blue reporter assay

RAW-Blue (NF-κB-SEAP) reporter cell line (Invivogen, raw-sp) was used to confirm the proper incorporation of TLRas with the SNPs and their potencies compared to soluble TLRas. RAW-blue cells were cultured at 37 °C with 5% CO_2_ in DMEM supplemented with L-glutamine (2 mM), D-glucose (4.5 g/L), 10% HI-FBS, and penicillin (100 U/mL)/streptomycin (100 µg). Every other passage, zeocin (100 µg/mL) was added to the culture medium. Serial dilutions of solvent controls, soluble SNP components (cholesterol, DPPC, saponin), soluble TLRas (TLR1/2 Pam3CSK4, TLR4 MPLAs, and TLR7/8 cholesteryl imidazoquinoline derivative), SNP, and TLRa-SNPs were added to a 96-well tissue culture treated plate to achieve final concentrations between 40 and 0.01 µg/mL of TLRas (or equivalent concentration of individual component based on TLRa-SNPs added for the negative controls). We assumed 100% TLRa incorporation during TLRa-SNP formulations to determine the TLRa concentrations. About 100,000 cells were added to each well in 180 µL of medium and were incubated for 24 h at 37 °C in a 5% CO_2_ incubator. Manufacturer instructions were followed for SEAP quantification, and absorbance levels were detected at 655 nm after 2 h incubation with QUANTI-Blue Solution (Invivogen). Nonlinear regression fits were estimated using the “Log(agonist) vs. response – EC50” function in GraphPad Prism 8.4 software.

### In vivo lymph node accumulation of NP antigen with TLRa-SNPs

C57BL/6 mice were injected subcutaneously at the tail base with 100 μL of PBS 1X buffer containing 10 μg AF647-NPs and 10 μg of SNP, 10 μg of TLRa-SNPs, or 20 μg of CpG and 100 μg of Alum. Mice were euthanized 48h post-injection with CO_2_ and their inguinal draining lymph nodes were imaged using an In Vivo Imagining System (IVIS Lago). Imaging procedures and data analysis methods were identical to those thoroughly described in previously published work (*69–71*). AF647-NPs were imaged using 30 second exposure time, excitation wavelength of 660 nm, and emission wavelength of 710 nm, medium binning, and a F/stop of 4. Total radiant efficiency was quantified using the Living Image software version 4.2 with the region of interest defined as an oval of consistent size around the entire lymph node.

### Preparation of RBD-NPs

The nanoparticle immunogen (RBD-NP) components and the nanoparticle formulation methodology have been previously described (*27*). RBD-NP was kept in the following buffer conditions: 50 mM Tris pH 8, 150 mM NaCl, 100 mM l-arginine and 5% sucrose.

### Vaccine Formulation

The model vaccines included an antigen of either 1.5 μg dose of RBD-NP (University of Washington) or 10 μg dose of gp120 (SIV/mac239, Immune Technology Corporation). Depending on the formulations, vaccines also contained an adjuvant of CpG1826/Alum (20 μg + 100 μg, respectfully, Invivogen), SNP, or one of the four TLRa-SNPs with an equivalent dose of 10 μg of Quil-A saponins (TLR1/2a-SNP, TLR4a-SNP, TLR7/8a-SNP, or TLR4a-TLR7/8a-SNP). An additional vaccine was formulated with an equivalent dose of 5 μg saponins of TLR4a-SNP and 5 μg saponins of TLR7/8a-SNP to test their synergistic effects. CpG/Alum group adjuvanted with RBD-NP was prepared similarly as described previously by Pulendran and coworkers as the clinical control,(*29, 30*) although both the CpG and Alum dosages were doubled in the present study. Vaccines were prepared in PBS 1X to a volume of 100 μl per dose and loaded into syringes with a 26-gauge needle for subcutaneous injection. Mice were boosted on Week 3 for RBD-NP vaccines or Week 4 for gp120 vaccines.

### Mice and Vaccination

Six-to-seven week old female (except for one experiment where male mice were vaccinated with gp120 vaccines) C57BL/6 (B6) mice were purchased from Charles River and housed in the animal facility at Stanford University. Several days before vaccine administration, mice were shaved on their right flank. On the day of injection, 100 µL of soluble vaccines were subcutaneously injected on the right side of their flank under brief isoflurane anesthesia. Mouse blood was collected from the tail vein following the schematic shown in Figure 2 and Figure 7. The inguinal dLNs were collected for GC analysis as well as Luminex after euthanasia and spleens were collected for splenocyte ELISpot analysis after euthanasia.

### Mouse serum antibody concentration and affinity with ELISAs

Serum Anti-RBD or Anti-gp120 IgG antibody endpoint titers were measured using an endpoint ELISA. Maxisorp plates were coated with SARS-CoV-2 RBD protein, spike protein, a spike protein variant (B.1.1.7, B.1.351, B.1.617.2, or B.1.529), or gp120 at 2 µg/mL in PBS 1X overnight at 4 °C and subsequently blocked with PBS 1X containing 1 wt% BSA for 1 h at 25 °C. Plates were washed 5x in between each step with PBS 1X containing 0.05% of Tween-20. Serum samples were diluted in diluent buffer (PBS 1X containing 1% BSA) starting at 1:100 followed by a 4-fold serial dilution and incubated in the coated plates for 2 h at 25 °C. Then, goat-anti-mouse IgG Fc-HRP (1:10,000), IgG1 Fc-HRP (1:10,000), or IgG2c Fc-HRP (1:10,000) was incubated for 1h at 25 °C. Plates were developed with TMB substrate for 6 mins and the reaction was stopped with 1Ν HCl aqueous solution. Plates were analyzed using a Synergy H1 microplate reader (BioTek Instruments) at 450 nm in absorbance. Total IgG, subtypes, and variants were imported into GraphPad Prism 8.4.1 to determine serum titers by fitting the curves (absorbance vs dilution) with a three-parameter non-linear logistic regression (baseline constrained to 0.054, the negative control average). The dilution titer value at which the endpoint threshold (0.2 and 0.1 for RBD-NP and gp120 vaccines, respectively) was crossed for each curve was reported as endpoint titer. Samples failing to meet endpoint threshold at a 1:100 dilution were set to below the limit quantitation for the assay (which we plotted as 1:25 dilution).

For affinity measurements, competitive ELISAs were performed in a similar manner to previously published procedures (*72*). In these assays, 2X dilutions of the serum (starting at 1:50) were mixed with a constant 0.07 nM concentration of an HRP-conjugated anti-RBD monoclonal antibody (Sino Biological) and incubated for 2h at room temperature. Plates were developed with TMB substrate for 6 mins and the reaction was stopped with 1N HCl aqueous solution. Plates were analyzed using a Synergy H1 microplate reader (BioTek Instruments) at 450 nm in absorbance. Data were fitted with a one-site competitive binding model using GraphPad Prism 10. The control mAb reference antibody was assumed to have a K_D_ of 1 nM based on common affinities found in the industry. This assumption only affects the absolute K_D_ values reported and not the relative differences between treatment groups.

### SARS-CoV-2 Spike-pseudotyped Viral Neutralization Assay

Neutralization assays were conducted as described previously (*73*). Briefly, SARS-CoV-2 spike-pseudotyped lentivirus was produced in HEK239T cells. Six million cells were seeded one day prior to transfection. A five plasmid system (*74*) and BioT (Bioland Scientific) were used for viral production per manufacturer’s protocol with the full-length wild-type spike sequence from the Wuhan-Hu-1 strain of SARS-CoV-2. Virus-containing culture supernatants were harvested ≈72 hours after transfection by centrifugation and filtered through a 0.45 µm syringe filter. Stocks were stored at −80 °C prior to use. For the neutralization assay, ACE2/HeLa cells were plated 1–2 days prior to infection (*75*). Mouse and human sera were heat inactivated at 56°C for 30 min prior to use. Sera and viruses were diluted in cell culture medium and supplemented with a polybrene at a final concentration of 5µg/mL. Serum/virus dilutions were incubated at 37°C for 1 h. After incubation, media was removed from cells and replaced with serum/virus dilutions and incubated at 37 °C for 2 days. Cells were then lysed using BriteLite (Perkin Elmer) luciferase readout reagent, and luminescence was measured with a BioTek plate reader. Each plate was normalized by wells with cells or virus only and curves were fit with a three-parameter non-linear logistic regression inhibitor curve to obtain IC_50_ values. Serum dilution curves display mean infectivity ± SEM for each individual mouse at each serum dilution. Normalized values were fit with a three-parameter non-linear logistic regression inhibitor curve in GraphPad Prism 8.4.1 to obtain IC_50_ values. Fits were constrained to have a value of 0% at the bottom of the fit.

### Luminex Cytokine Assay

Mice were immunized with RBD-NP and different TLRa-SNPs as described in the immunization section. 24 hours after injection, mice were euthanized and the injection inguinal dLNs were excised and processed for analysis (*76*). Briefly, the dLNs were weighted and 10 mL of lysis buffer (Cell Extraction Buffer FNN0011 with 1x Halt protease inhibitor, Themo Fisher) was added per 1 g of tissue. Tissues were then grinded using a Dounce Tissue Grinder and centrifuged at 10,000 RCF for 5 min using protein lo-bind Eppendorf tubes to pellet tissue debris. Supernatants were collected into new protein lo-bind Eppendorf tubes. Meanwhile, a Pierce BCA assay (Thermo Fisher) was set up to analyze the total protein content of the samples. When possible, samples were diluted and normalized to 2 mg/mL of total sample protein content per sample using additional lysis buffer. Tissues/homogenates were kept on ice throughout processing and normalized samples were stored at -80 °C. Injection dLN homogenates were provided to the HIMC for analysis by a 48-plex murine Luminex cytokine panel.

### Dimensional Reduction Analysis via Penalized Supervised Star Plot

Cytokine MFI data were detrended for cage and nonspecific binding effects using ordinary least squares (*77*). Detrended data were used to construct a penalized supervised star plot (*78*). Analyses were conducted in SAS® v.9.4 (SAS® Institute, Cary, North Carolina, USA) and R (www.r-project.org). Star plots were constructed with R packages maptools (*79*), matrixcalc (*80*), plotrix (*81*), JPEN (*82*), and VCA (*83*).

### Immunophenotyping in Lymph Nodes by Flow Cytometry

According to previously reported methodology (*72*), mice were euthanized using CO_2_ 12 (RBD-NP vaccines) or 14 (gp120 vaccines) days post immunization. Inguinal LNs were collected and dissociated into single cell suspensions. Cells were stained for viability using Ghost Dye Violet 510 (Tonbo Biosciences, Cat: 13-0870-T100) for 5 min on ice and washed with FACS buffer (PBS 1X with 2% FBS, 1mM EDTA). Fc receptor was blocked using anti-CD16/CD38 (clone: 2.4G2, BD Biosciences, cat: 553142) for 5 min on ice and then stained with fluorochrome conjugated antibodies: CD19 (PerCP-Cy5.5, clone: 1D3, BioLegend, cat: 152406) CD95 (PE-Cy7, clone: Jo2, BD Biosciences, cat: 557653), CD38 (BUV395, clone: 90, BD Biosciences, Cat: 740245), CXCR4 (BV421, clone: L276F12, BioLegend, cat: 146511), CD86 (BV785, clone: GL1, BioLegend, cat: 105043), GL7 (AF488, clone: GL7, BioLegend, cat: 144613), CD3 (AF700, clone: 17A2, BioLegend, cat: 100216), CD4 (BV650, clone: GK1.5, BioLegend, cat: 100469), CXCR5 (BV711, clone: L138D7, BioLegend, cat: 145529) and PD1 (PE-Dazzle^TM^594, clone: 29F.1A12, BioLegend, cat: 135228) for 30 min on ice (Table S2). Cells were washed, fixed with 4% PFA on ice, washed again and analyzed on an LSRII flow cytometer (BD Biosciences). Data were analyzed with FlowJ 10 (FlowJo, LLC, representative gating scheme in Figure S9).

### ELISpot

The number of antigen-specific IFN-γ producing splenocytes were determined to correlate with antigen-specific IFN-γ producing CD8+ T cells using a Mouse IFN-γ Single-Color ELISPOT kit (CTL ImmunoSpot). Spleen cells were harvested 2 weeks post-boost (D35) following the immunization outlined in Figure S13A. 800,000 splenocytes per sample were pipetted into the well and were stimulated with 1 μg/mL Pan-SARS-CoV-2-RBD peptides (JPT) for 24 hours at 37°C. Spots were then developed following manufacturer’s instruction (Immunospot S6 Ultra M2).

### Animal Protocol

Mice were cared for following the Institutional Animal Care and Use guidelines. All animal studies were performed in accordance with the National Institutes of Health guidelines and the approval of Stanford Administrative Panel on Laboratory Animal Care (Protocol APLAC-32109).

### Collection of Serum from Human Patients

Convalescent COVID-19 blood was collected from donors 8–12 weeks after the onset of symptoms, all donors had previously been immunized with COVID vaccines authorized by the FDA (total of three immunizations, excluding the omicron bivalent booster). Blood was collected in microtubes with serum gel for clotting (Starstedt), centrifuged for 5 min at 10,000 g and then serum was stored at −80 °C until used. Blood collection was performed by finger-prick in accordance with National Institutes of Health guidelines, the approval of the Stanford Human Subjects Research and IRB Compliance Office (IRB-58511) and the consent of the individuals.

### Statistical Analysis

For *in vivo* experiments, animals were caged blocked. Mead’s Resource Equation was used to calculate the sample size above which additional subjects (n) will not have significant impact on power (*33, 34*). For most experiments, a sample size of *n* = 5 per group was used. A few mice died due to reasons external to the treatment effects as evaluated by the veterinarian staff. Groups resulting with *n* = 4 still had enough power for statistical analyses using Mead’s Resource Equation. No animals were considered outliers.

All results except for Luminex are expressed as mean ± standard error of mean (SEM). Unless otherwise specified, comparison between multiple groups and *p* values were conducted with the general linear model (GLM) followed by Tukey HSD procedure test in JMP and we accounted and adjusted for unequal variances, cage blocking, and multiple comparison. Statistical significance was considered as *p* < 0.05. Selected *p* values are shown in the text and the figures and remaining *p* values are in the supporting information.

For the Luminex assay, statistical analyses were provided by the Statistical Consultation Service of the Human Immune Monitoring Center at the Institute for Immunity, Transplantation and Infection, Stanford University School of Medicine. The logarithm of median fluorescence intensity (MFI) was regressed on treatment group, cage, and logarithm of nonspecific binding MFI using generalized maximum entropy estimation regression (*84*). Within each treatment group, *p* values were adjusted to control the false discovery rate (*85, 86*) at 5% across all cytokines. Analyses and plotting were done in SAS® (SAS® Institute, Cary, North Carolina, USA).

## Acknowledgments

This work was financially supported by the Center for Human Systems Immunology with the Bill & Melinda Gates Foundation (OPP1113682; OPP1211043; INV027411; INV-010680) and the National Institute of Allergy and Infectious Disease (P01AI167966). BSO is grateful for an Eastman Kodak Fellowship. MVFI is thankful for the support by the National Institutes of Health under award numbers T32GM007365, F30AI152943. XZ is supported by a Stanford ChEM-H COVID-19 Drug and Vaccine Prototyping seed grant. OMS, JY, and CJ are thankful for a National Science Foundation Graduate Research Fellowship. OMS is also thankful for Hancock Fellowship of the Stanford Graduate Fellowship in Science and Engineering. ELM is grateful for the NIH Biotechnology Training Program (T32 GM008412). The authors would like to thank former and current members of the Appel lab for their technical expertise and scientific discussion, especially Dr. Celine S. Liong, Dr. Catherine M. Kasse and Anahita Nejatfard for their help in animal work and advice. We acknowledge Dr. Wah Chiu and Dr. Peter Kim for their assistance and advice. We extend our gratitude to Dr. Tyson H. Holmes and Dr. Yael Rosenberg-Hasson at the Stanford Human Immune Monitoring Center as well as Duo Xu for his invaluable advice in implementing the spike-pseudotyped lentiviral neutralization assay (to MVFI). We also would like to thank the human patients for donating their blood, the staff of the BioE/ChemE Animal Facility who cared for the mice and the Stanford Core Facilities that were central to the completion of this work. Specifically, we thank the Vincent Coates Foundation Mass Spectrometry Laboratory (Stanford University Mass Spectrometry - RRID:SCR_017801), and the Department of Chemistry Nuclear Magnetic Resonance facility (Stanford University, NIH High End Instrumentation grant 1 S10 OD028697-01). Cryo-Electron Microscopy was performed at the Stanford-SLAC facility (NIH shared instrumentation grant S10OD021600). FACS sorting was performed on an instrument in the Shared FACS Facility obtained using NIH S10 Shared Instrument Grant (S10RR027431-01).

## Funding

Bill & Melinda Gates Foundation (OPP1113682; OPP1211043; INV027411; INV-010680)

National Institute of Allergy and Infectious Disease (P01AI167966)

BSO: Eastman Kodak Fellowship

MVFI: National Institutes of Health under award numbers (T32GM007365; F30AI152943)

OMS, JY, CKL: National Science Foundation Graduate Research Fellowship

OMS: Hancock Fellowship of the Stanford Graduate Fellowship in Science and Engineering

XZ: Stanford ChEM-H COVID-19 Drug and Vaccine Prototyping Seed Grant

ELM: NIH Biotechnology Training Program (T32 GM008412)

NIH High End Instrumentation grant 1 (S10 OD028697-01)

NIH shared instrumentation grant (S10OD021600)

NIH S10 Shared Instrument Grant (S10RR027431-01)

## Author contributions

Conceptualization: BSO, JB, OMS, JY, BP, EAA.

Methodology: BSO, JB, JZA, OMS, JY, ASV, EAA

Investigation: BSO, JB, MVFI, JZA, XZ, OMS, JY, JHK, CKJ, ELM, LC, BP

Visualization: BSO, JB Supervision: BSO, JB, NPK, EAA

Resources: BSO, JB, MVFI, XZ, JY, ASV, LC, BP, NPK

Data curation: BSO

Validation: BSO, JB, JZA, XZ, OMS, NPK, EAA

Formal analysis: BSO, JB, MVFI, JZA, CKJ

Project administration: BSO, JB, ASV, NPK, EAA

Funding acquisition: BP, NPK, EAA

Writing—original draft: BSO, JB

Writing—review & editing: BSO, JB, XZ, OMS, JHK, NPK, EAA

All authors have given approval to the final version of the manuscript. ‡These authors contributed equally.

## Data and materials availability

All data needed to evaluate the conclusions in the paper are present in the paper and/or the Supplementary Materials.

## Table of contents for supplementary materials

Supplementary Methods

FiguresS1 to S13

Tables S1 to S30

Supplementary Data

