## Supplemental Information for "Saponin Nanoparticle Adjuvants Incorporating Toll-Like Receptor Agonists Drive Distinct Immune Signatures and Potent Vaccine Responses"

Ben S. Ou *et al.*

**This PDF file includes:**

Supplementary Methods
Figures S1 to S13
Tables S1 to S30
Supplementary Data

### Supplemental Methods

#### Materials

4-amino-2-(ethoxy methyl)-1H-imidazo[4, 5-C]quinoline-1-butylamine was purchased from Career Henan Chemical Co. (Henan Province, China) and used as received. All other chemicals and solvents (Sigma-Millipore, VWR International) were used as received, unless specified otherwise. Reactions requiring anhydrous conditions were conducted with dry solvents under an inert atmosphere (nitrogen). Analytical thin layer chromatography (TLC) was carried out on aluminum-baked silica gel matrix plates (Sigma Aldrich) visualized using UV light (254 nm) and/or revealed with 1% aqueous KMnO4 followed by heating. Colum chromatography was performed on silica gel (0.063-0.200 mm).

#### NMR Spectroscopy

Compounds were characterized by $^{1}H$, $^{13}C$, COSY, HSQC experiments on a Brucker neo500 ($^{1}H$: 500 MHz, $^{13}C$: 125 MHz) spectrometers. Chemical shifts (δ) are given in parts per million (ppm) and coupling constants J in Hertz (Hz); peak multiplicity is reported as follow: s = singlet, bs = broad singlet, d = doublet, dd = doublet of doublets, t = triplet, q = quartet, m = multiplet. High Resolution Mass Spectrometry was performed on a Orbitrap Exploris 240 BioPharma Mass Spectrometer (Stanford University Mass Spectrometry) with heated electrospray ionization (HESI).

#### Synthesis and characterization of Cholesteryl imidazoquinoline derivative

100 mg of 4-amino-2-(ethoxy methyl)-1H-imidazo[4, 5-C]quinoline-1-butylamine (100 mg, 0.319 mmol) was dissolved in 8 mL of anhydrous $CH_2Cl_2$/DMF (85:25). In parallel, cholesteryl chloroformate (145 mg, 0.322 mmol, 1.01 eq.) was dissolved in 3 mL of anhydrous $CH_2Cl_2$ and triethylamine (53 µL, 0.383 mmol, 1.2 eq.) was then added. The solution was added dropwise at 0 °C into the first one and the reaction was stirred at room temperature for 16 hours. Solvents were removed under reduced pressure and the resulting solid was purified by column chromatography on silica gel eluting with a step gradient of MeOH/$Et_3N$ (99:1) from 0 % to 6 % v/v in $CH_2Cl_2$ (Rf: 0.5, 9/1 $CH_2Cl_2$/MeOH). The title compound was obtained as a white solid foam (90 mg, 39 %).

$^{1}$**H NMR** (500 MHz, $CD_2Cl_2$) $\delta$ (ppm): 0.69 (s, 3H), 0.86 (d, *J*= 2.0 Hz, 3H), 0.87 (d, *J*= 2.2 Hz, 3H), 0.90-1.62 (m, 28H), 1.65-1.75 (m, 2H), 1.78-1.89 (m, 3H), 1.91-2.08 (m, 6H), 2.19-2.34 (m, 2H), 3.16-3.27 (m, 2H), 3.60 (q, *J*= 7.2 Hz, 2H), 4.37-4.47 (m, 1H), 4.52-4.60 (m, 2H), 4.78 (s, 1H), 4.82-4.90 (m, 2H), 5.34-5.39 (m, 1H), 5.61 (bs, 1H), 7.31-7.36 (m, 1H), 7.50 (m, 1H), 7.75 (dd, *J*= 1.4, 8.3 Hz, 1H), 7.95 (d, *J*= 8.3 Hz, 1H).

$^{13}$**C NMR** (125 MHz, $CD_2Cl_2$) $\delta$ (ppm): 12.0, 15.3, 18.9, 19.5, 21.4, 22.7, 22.9, 24.2, 24.6, 27.6, 27.8 28.4, 28.5, 28.8, 32.2, 32.3, 36.2, 36.5, 36.9, 37.4, 38.9, 39.9, 39.9, 40.1, 40.5, 42.7, 46.3, 50.5, 56.5, 57.1, 65.5, 66.6, 74.6, 115.8, 120.4, 122.7, 127.0, 127.3, 127.6, 134.3, 140.4, 145.2, 149.5, 151.8, 156.4

#### Supplemental Figures

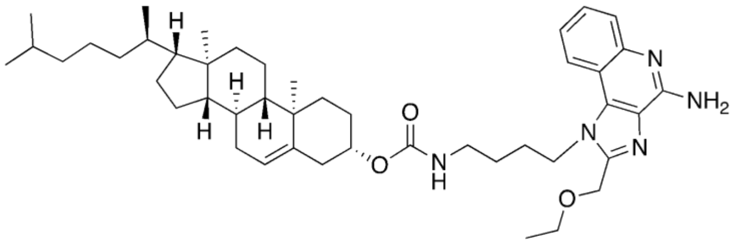

Chemical Formula: $C_{45}H_{67}N_5O_3$

**Figure S1. Chemical Structure of Cholesteryl imidazoquinoline derivative**

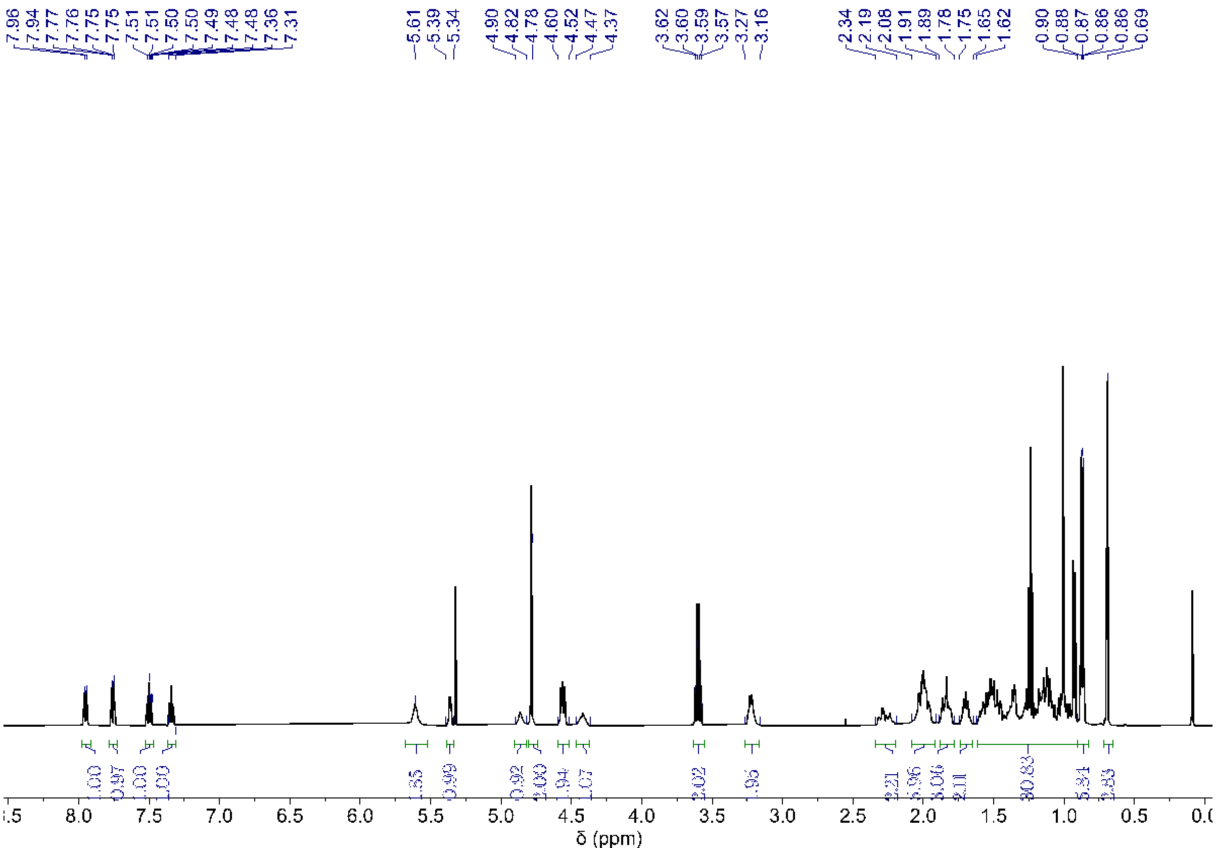

**Figure S2. $^1$H NMR spectrum of Cholesteryl imidazoquinoline derivative (500 MHz, $CD_2Cl_2$)**

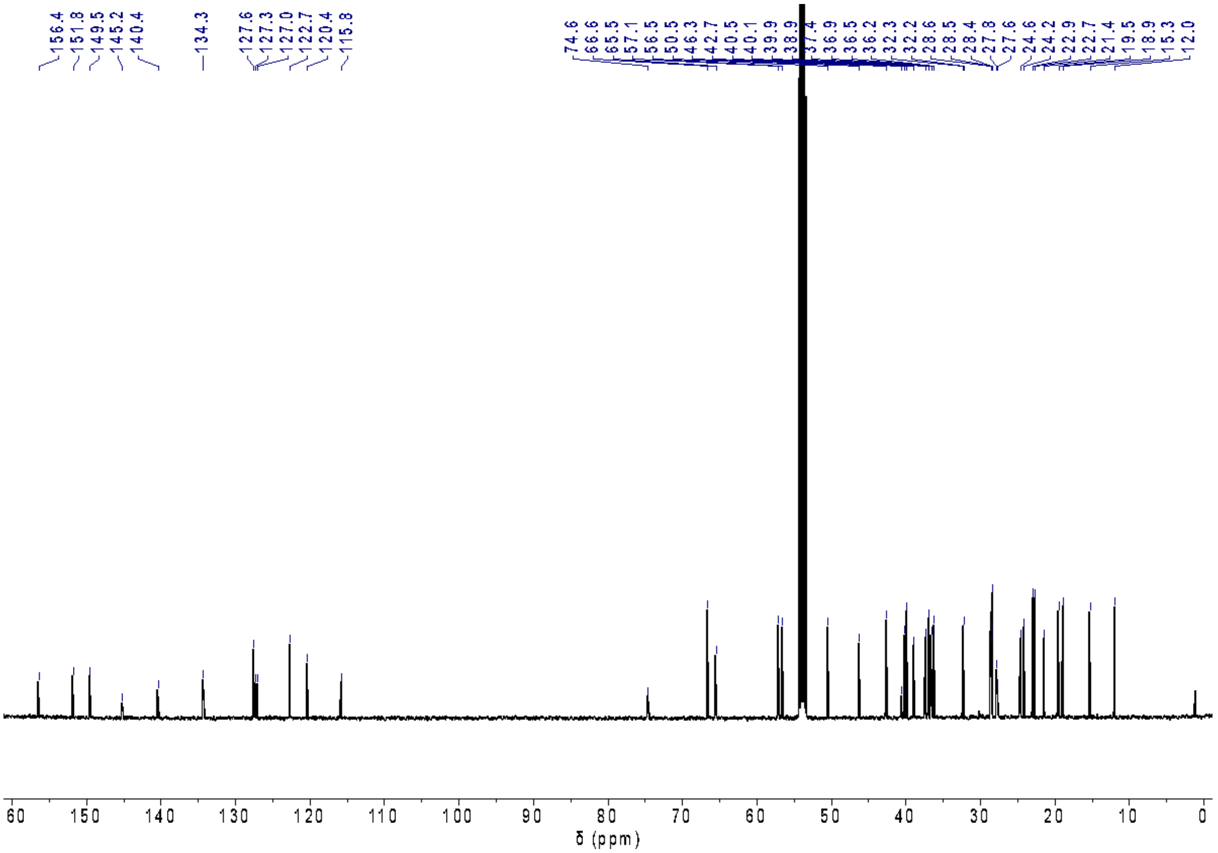


**Figure S3. $^{13}$C NMR spectrum of Cholesteryl imidazoquinoline derivative (125 MHz, $CD_2Cl_2$)**

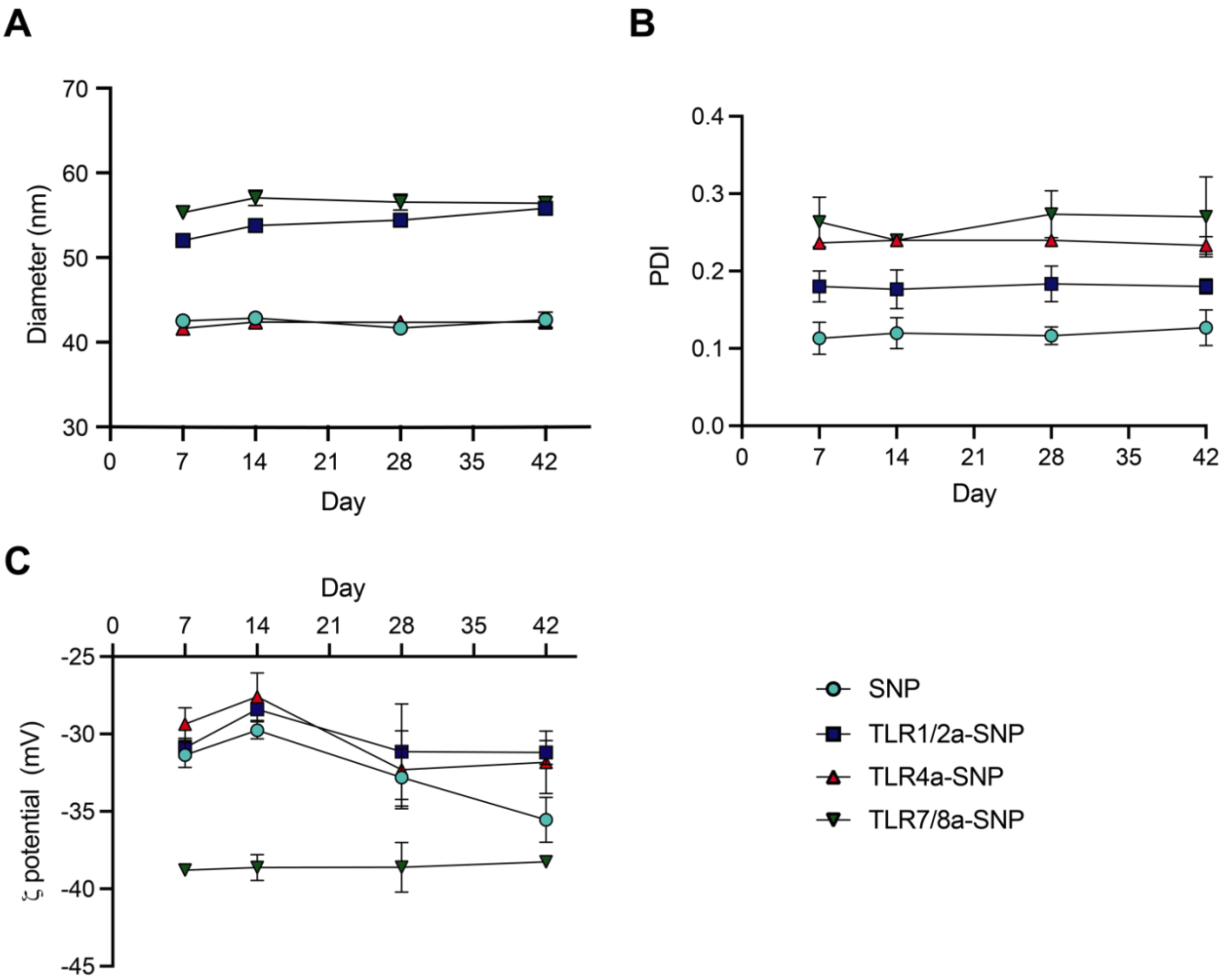


**Figure S4. Colloidal stability of SNP and TLRa-SNPs over the course of 42 days. (A)** Hydrodynamic diameter, **(B)** PdI, and **(C)** surface charge of SNP and TLRa-SNPs. Formulations were stored at 4°C.

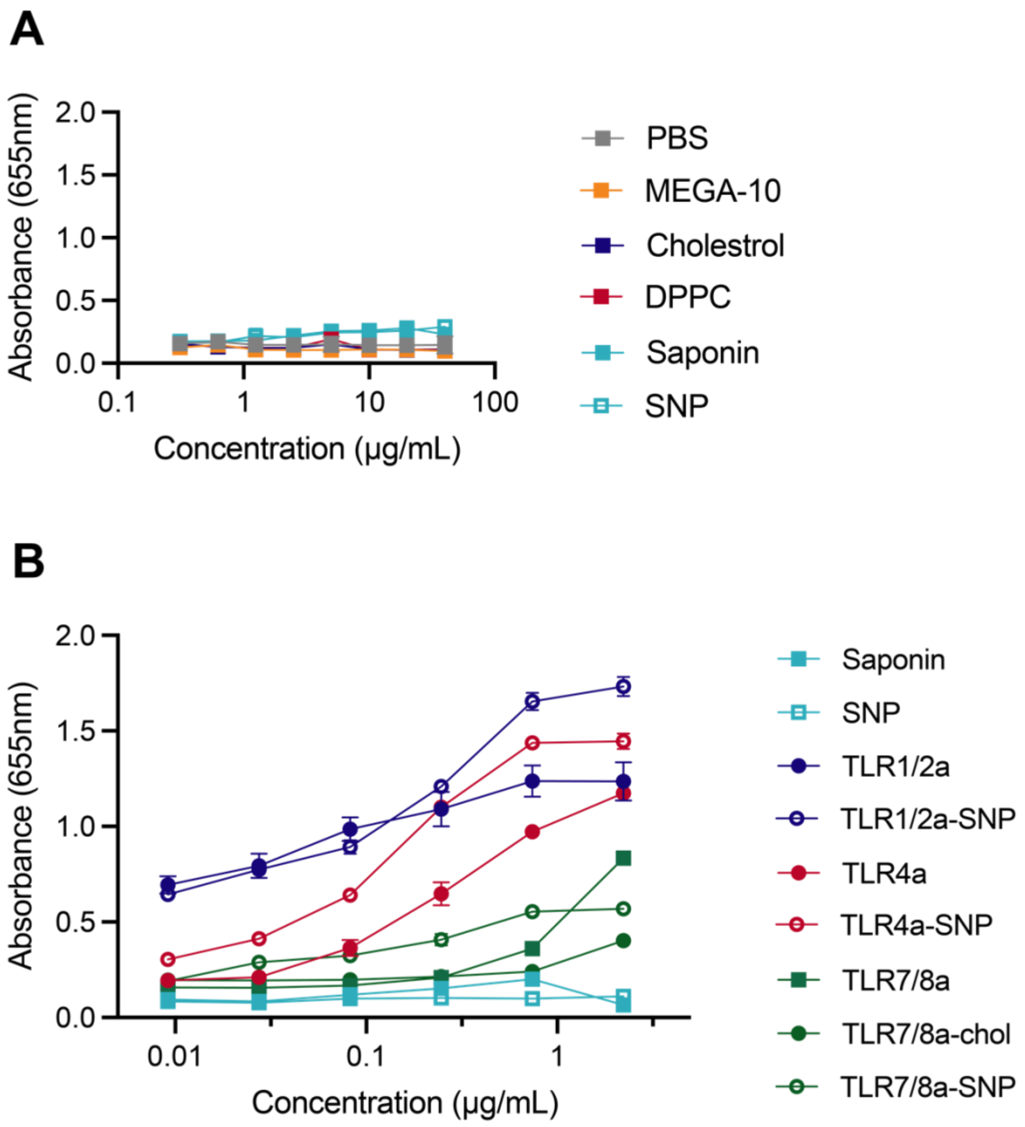


**Figure S5. In vitro RAW-Blue activation curves of (A)** relevant chemicals and buffers and **(B)** soluble TLRas, SNP, and TLRa-SNPs.

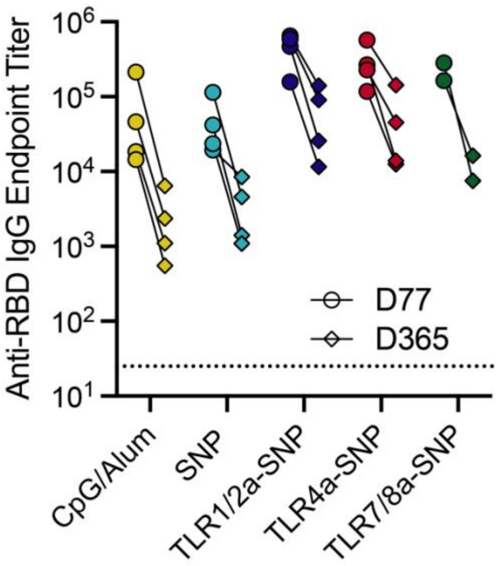


**Figure S6. Anti-RBD IgG endpoint titers drop of different RBD-NP vaccines with different adjuvants from D77 to D365.**

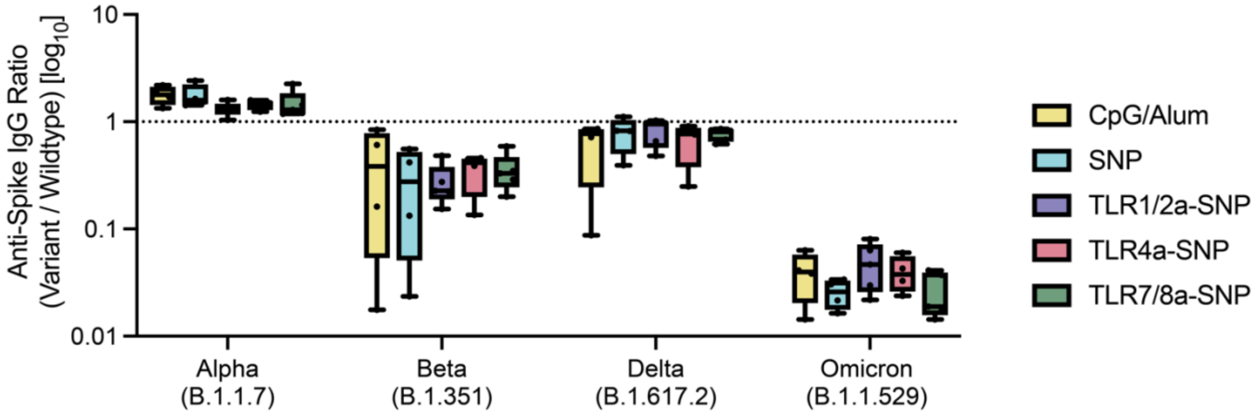


**Figure S7. Anti-spike IgG endpoint titer drop against SARS-CoV-2 variants of concern and WT spike titer of RBD-NP vaccines with different adjuvants.** Boxes shown are interquartile range.

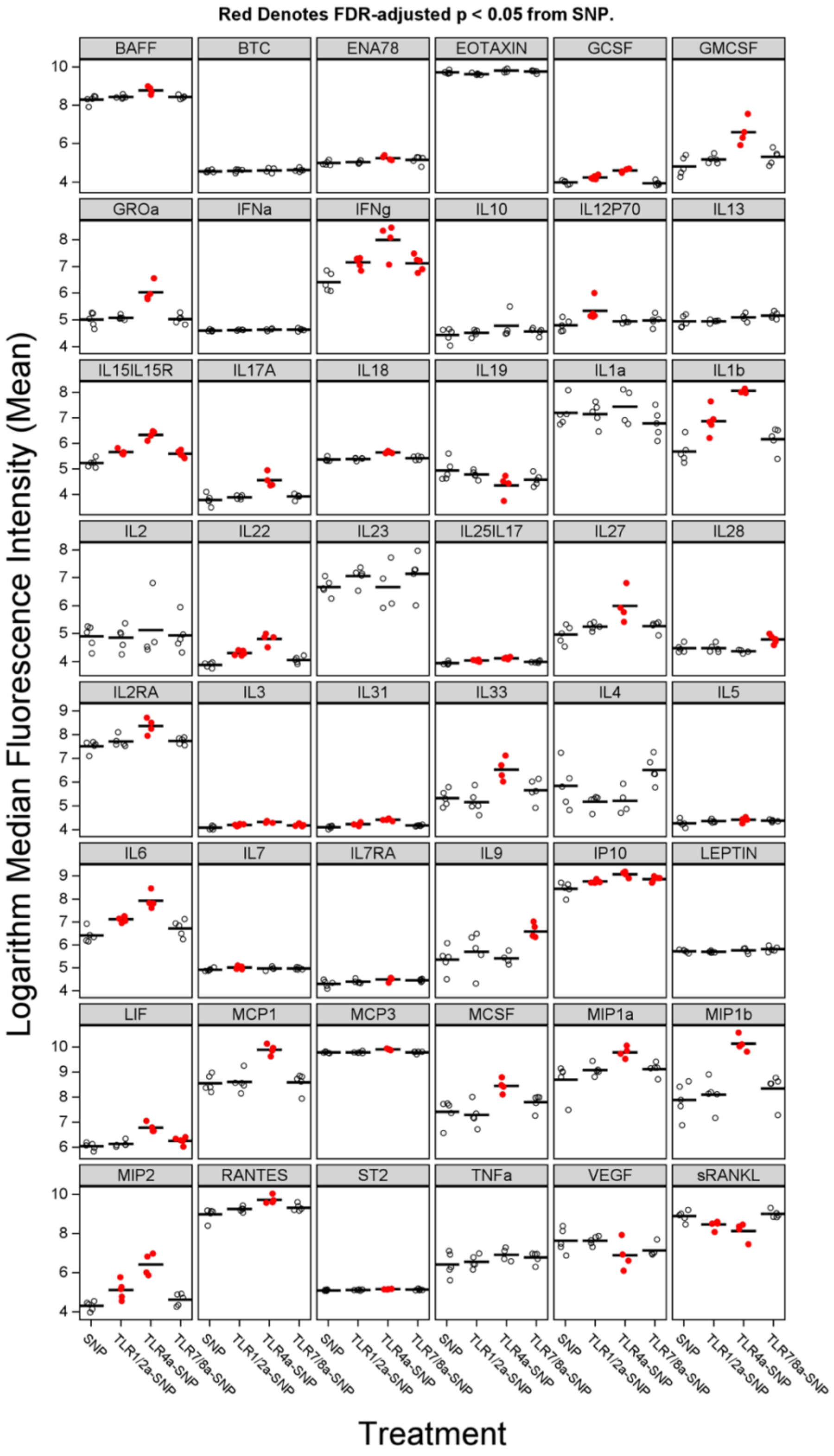


**Figure S8. Full artifact corrected Luminex dataset for serum cytokine levels 24 hours after vaccine immunization in logarithm Median Fluorescence Intensity (MFI).** Plotted data are detrended for covariates of cage and nonspecific bindings and bars shown are mean.

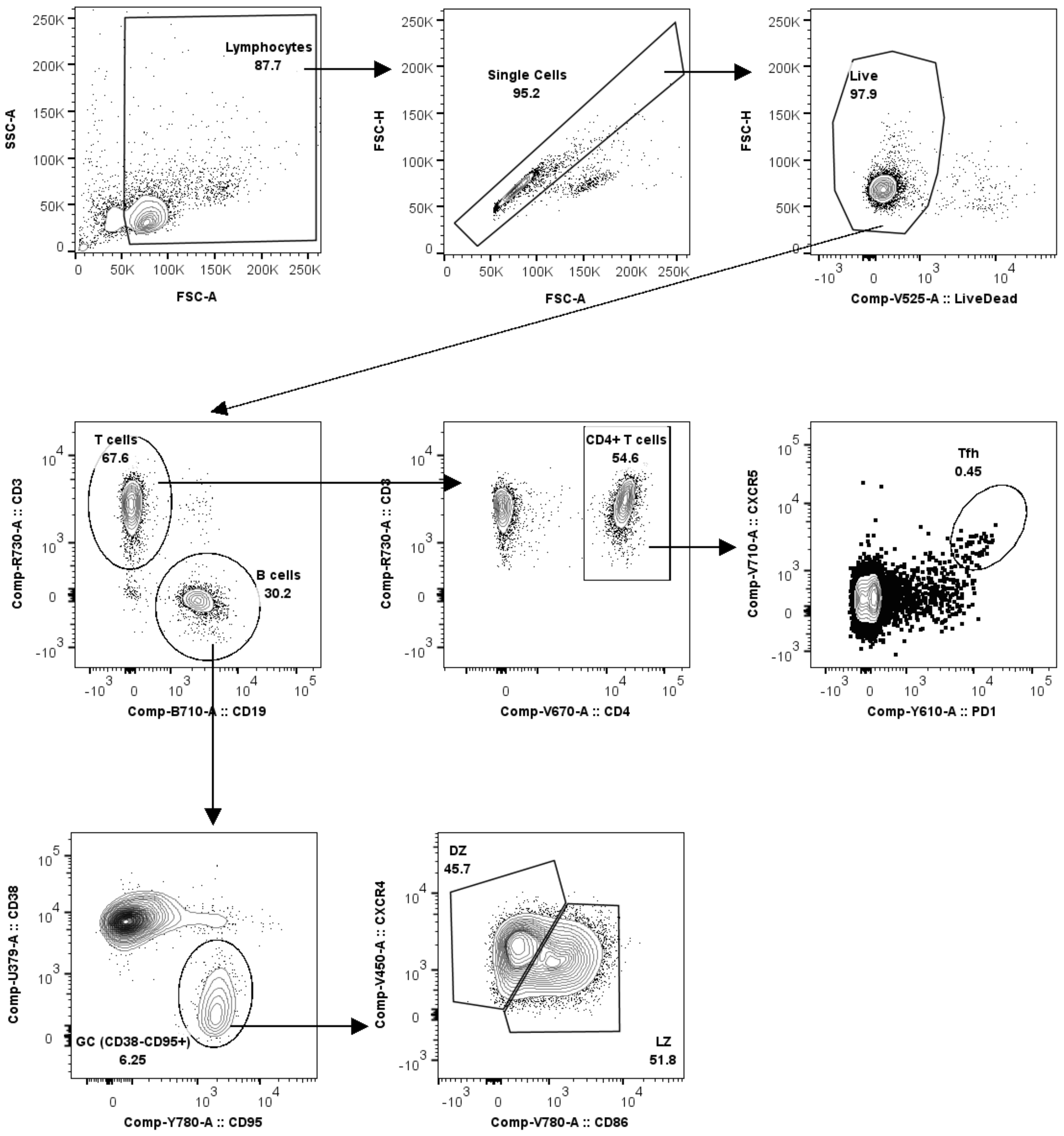


**Figure S9. Representative gating strategy for GC analysis. GCBCs were defined as CD38-CD95+.**

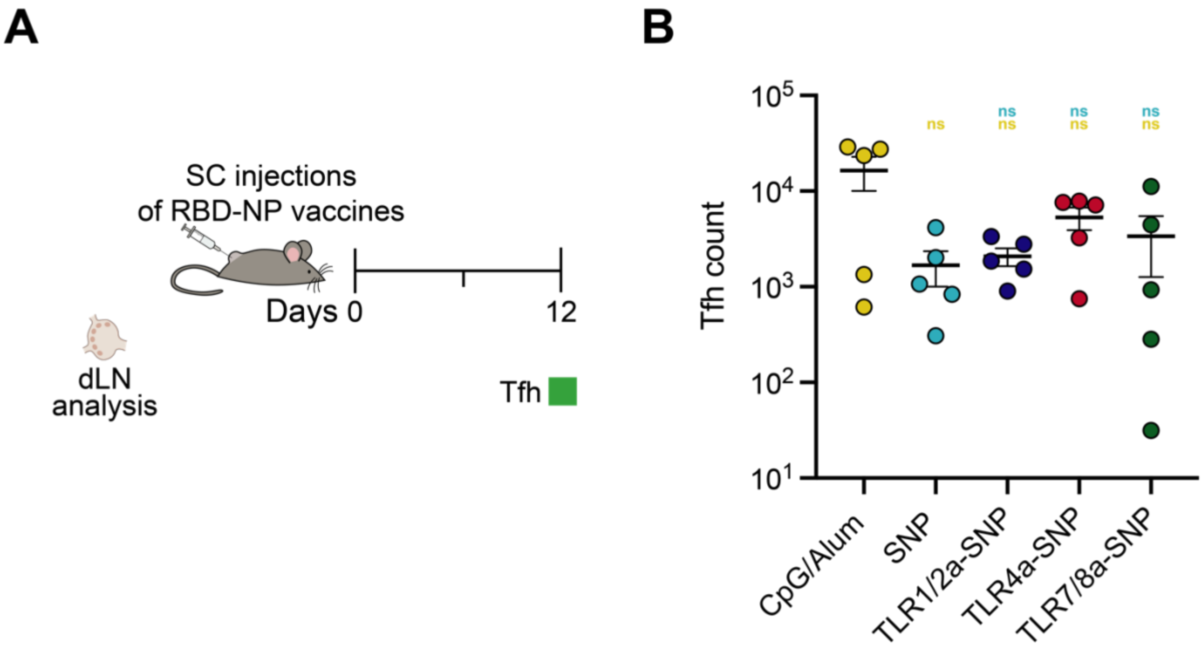


**Figure S10. Draining lymph nodes (dLNs) analysis post-immunization with RBD-NP vaccines adjuvanted with SNP or TLRa-SNPs. (A)** T-follicular helper cells (Tfh) were measured 12 days after immunization. **(B)** Total Tfh count from RBD-NP vaccines. Data (n = 5) are shown as mean +/- SEM. *p* values were determined using the general linear model followed by Tukey's HSD comparison procedure on the logged values. Complete *p* values for comparisons are shown in Table S17.

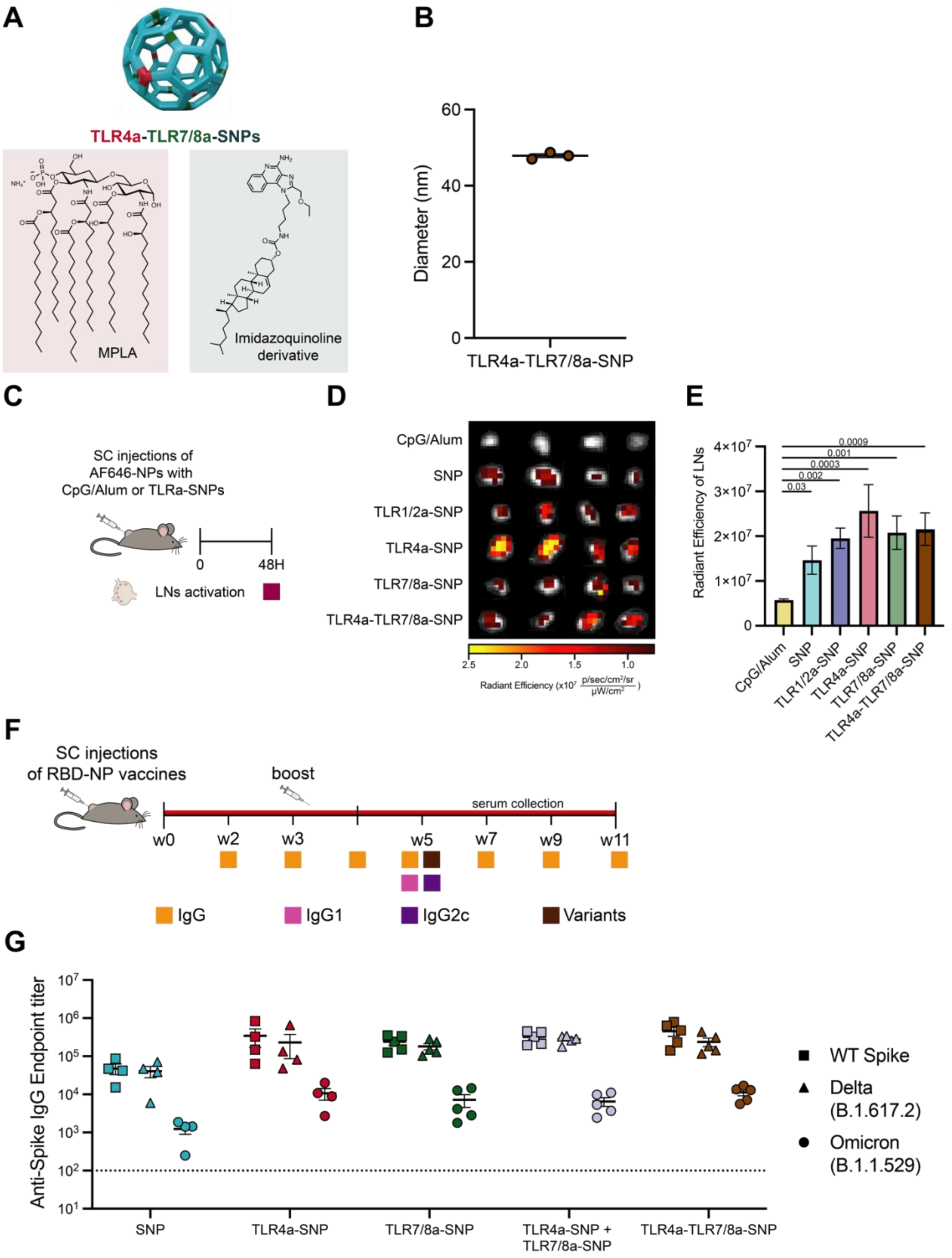

A
TLR4a-TLR7/8a-SNPs
MPLA
Imidazoquinoline derivative
B
Diameter (nm)
TLR4a-TLR7/8a-SNP
C
SC injections of AF646-NPs with CpG/Alum or TLRa-SNPs
0
48H
LNs activation
D
CpG/Alum
SNP
TLR1/2a-SNP
TLR4a-SNP
TLR7/8a-SNP
TLR4a-TLR7/8a-SNP
Radiant Efficiency
E
Radiant Efficiency of LNs
0.0009
0.001
0.0003
0.002
0.03
F
SC injections of RBD-NP vaccines
boost
serum collection
w0
w2
w3
w5
w7
w9
w11
IgG
IgG1
IgG2c
Variants
G
Anti-Spike IgG Endpoint titer
SNP
TLR4a-SNP
TLR7/8a-SNP
TLR4a-SNP + TLR7/8a-SNP
TLR4a-TLR7/8a-SNP
WT Spike
Delta (B.1.617.2)
Omicron (B.1.1.529)

**Figure S11. TLR4a-TLR7/8a-SNP generated consistent binding titers against variants of concern. (A)** Schematic representation of TLR4a-TLR7/8a-SNP where saponin nanoparticles co-incorporate TLR4a MPLA and TLR7/8a imidazoquinoline derivative. **(B)** Hydrodynamic diameter of TLR4a-TLR7/8a-SNP. **(C)** Schematic of *in vivo* evaluation of antigen (AF647-NP) accumulation with different TLRa-SNPs and CpG/Alum control. **(D)** Fluorescence IVIS imagining of the inguinal draining lymph nodes 48h after subcutaneous injection at the tail base. **(E)** Quantification of the lymph node accumulations at 48h. **(F)** Timeline of immunization and blood collection to determine IgG titers. Mice were immunized on Week 0 and boosted on Week 3 with RBD-NP vaccines adjuvanted with either a mixture of TLR4a-SNP and TLR7/8a-SNP or TLR4a-TLR7/8a-SNP. Variants titers were determined on Week 5 and compared with other groups. **(G)** Anti-spike IgG binding endpoint titers. Titers were determined for wildtype WT spike as well as Delta (B.1.617.2), and Omicron (B.1.1.529) variants of the spike protein. Data were shown as mean +/- SEM. *p* values were determined with one-way ANOVA with Tukey test of the logged radiant efficiency values. Complete *p* values for comparisons are shown in Table S4.

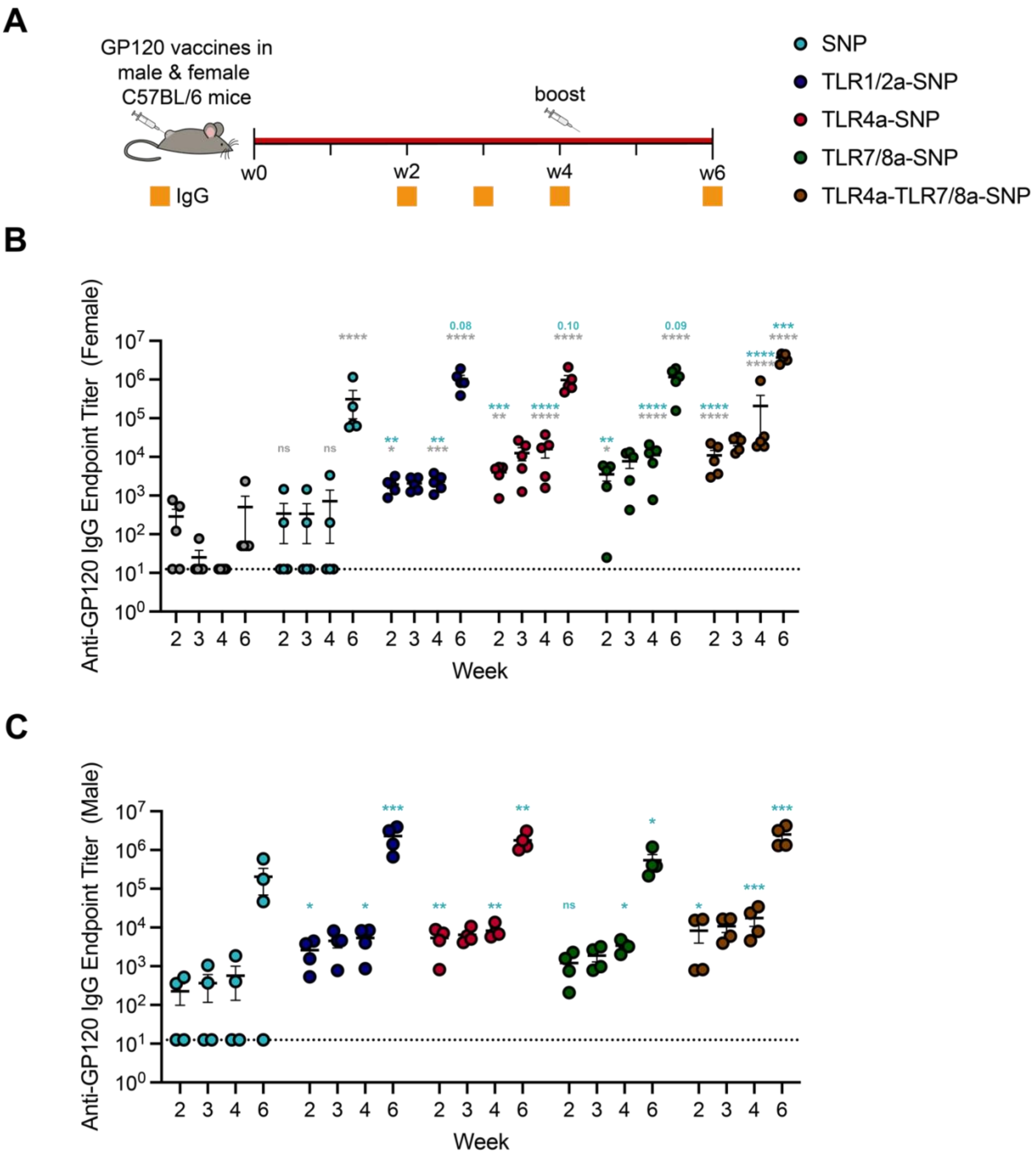


**Figure S12**. **In vivo humoral response to HIV gp120 vaccines adjuvanted with TLRa-SNPs in male and female mice. (A)** Timeline of immunization and blood collection to measure IgG titers. C57BL/6 mice of both sexes were immunized on Week 0 and boosted on Week 4 with gp120 vaccines adjuvanted with SNP or TLRa-SNPs. Anti-gp120 IgG binding endpoint titers in **(B)** female mice and **(C)** male mice. Data (n = 4-5) are shown as mean +/- SEM. Complete *p* values for comparisons are shown in Table S23 and S29. *p < 0.05, **p < 0.01, ***p < 0.001, and ****p < 0.0001.

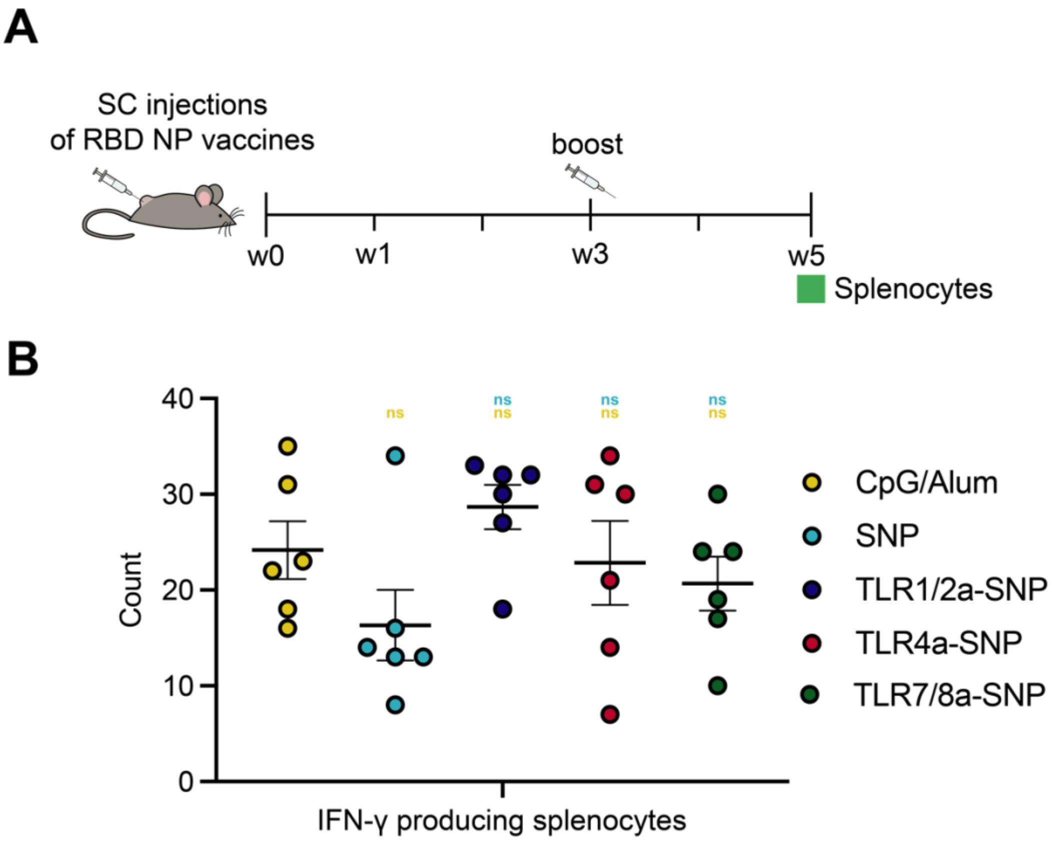


**Figure S13. Antigen-specific IFN-γ producing splenocytes analysis with ELISpot. (A)** Timeline of the experimental setup and spleen collection on Week 5 to determine antigen-specific CD8+ T cell population. **(B)** The number of IFN-γ producing CD8+ T cells upon antigen stimulation of 800,000 splenocytes/well of the vaccine groups. Data (n = 6) are shown as mean +/- SEM. Complete *p* values for comparisons are shown in Table S30.

#### Supplemental Tables

**Table S1.** Molar ratio of the formulation of SNP and TLRa-SNPs

| | **TLRa Adjuvant** | **Quil-A saponin** | **DPPC** | **Cholesterol** |
|---|---|---|---|---|
| **SNP** | - | 10 | 10 | 5 |
| **TLR1/2a-SNP** | 1 | 10 | 2.5 | 10 |
| **TLR4a-SNP** | 1 | 10 | 2.5 | 10 |
| **TLR7/8a-SNP** | 1 | 10 | 2.5 | 9 |
| **TLR4a-TLR7/8a-SNP** | 0.5 (TLR4a)<br>0.5 (TLR7/8a) | 10 | 2.5 | 9.5 |

**Table S2.** Flow cytometry antibody information

| Antibody (all anti-mouse) | Manufacturer | Clone |
|---|---|---|
| CD19-PerCP-Cy5.5 | BioLegend | 1D3 |
| CD95-PE-Cy7 | BD Biosciences | Jo2 |
| CD38-BUV395 | BD Biosciences | 90 |
| CXCR4-BV421 | BioLegend | L276F12 |
| CD86-BV785 | BioLegend | GL1 |
| GL7-A488 | BioLegend | GL7 |
| CD3-AF700 | BioLegend | 17A2 |
| CD4-BV785 | BioLegend | GK1.5 |
| CXCR5-BV711 | BioLegend | L138D7 |
| PD1-PE-Daazle™594 | BioLegend | 29F.1A12 |

**Table S3**. *p* values from a one-way ANOVA followed by Tukey multiple comparison test for the absorbance values at the highest TLRa concentration between different soluble TLRa and TLRa-SNPs (referring to Figure 1e-g, Figure S5).

| Equivalent TLRa concentration at 2.22µg/mL Absorbance value | | Adjusted *p* values |
|---|---|---|
| SNP | vs. TLR1/2a | **<0.0001** |
| SNP | vs. TLR1/2a-SNP | **<0.0001** |
| TLR1/2a | vs. TLR1/2a-SNP | **<0.0001** |
| SNP | vs. TLR4a | **<0.0001** |
| SNP | vs. TLR4a-SNP | **<0.0001** |
| TLR4a | vs. TLR4a-SNP | **0.0001** |
| SNP | vs. TLR7/8a-chol | **<0.0001** |
| SNP | vs. TLR7/8a-SNP | **<0.0001** |
| TLR7/8a-chol | vs. TLR7/8a-SNP | **0.0112** |

**Table S4.** *p* values from a one-way ANOVA followed by Tukey multiple comparison test for radiant efficiency of AF647-NPs accumulation in the lymph nodes compared between different TLRa-SNPs (referring to Figure 1j, Figure S11e).

| 48h lymph node accumulation<br>Radiant Efficiency [$\log_{10}$] | | Adjusted *p* values |
|---|---|---|
| CpG/Alum | vs. SNP | **0.0271** |
| CpG/Alum | vs. TLR1/2a-SNP | **0.0015** |
| CpG/Alum | vs. TLR4a-SNP | **0.0003** |
| CpG/Alum | vs. TLR7/8a-SNP | **0.0011** |
| CpG/Alum | vs. TLR4a-TLR7/8a-SNP | **0.0009** |
| SNP | vs. TLR1/2a-SNP | 0.7438 |
| SNP | vs. TLR4a-SNP | 0.2850 |
| SNP | vs. TLR7/8a-SNP | 0.6503 |
| SNP | vs. TLR4a-TLR7/8a-SNP | 0.5920 |
| TLR1/2a-SNP | vs. TLR4a-SNP | 0.9578 |
| TLR1/2a-SNP | vs. TLR7/8a-SNP | 0.6503 |
| TLR1/2a-SNP | vs. TLR4a-TLR7/8a-SNP | 0.9998 |
| TLR4a-SNP | vs. TLR7/8a-SNP | 0.9832 |
| TLR4a-SNP | vs. TLR4a-TLR7/8a-SNP | 0.9917 |
| TLR7/8a-SNP | vs. TLR4a-TLR7/8a-SNP | >0.9999 |

**Table S5**. *p* values from a general linear model (GLM) followed by Tukey's HSD multiple comparisons procedure for specific anti-RBD endpoint IgG titers time points compared between different RBD-NP vaccines (referring to Figure 2b).

| Week 2<br>IgG titers [$log_{10}$] | | Adjusted *p* values |
|---|---|---|
| CpG/Alum | vs. SNP | 0.8787 |
| CpG/Alum | vs. TLR1/2a-SNP | **0.0015** |
| CpG/Alum | vs. TLR4a-SNP | **<0.0001** |
| CpG/Alum | vs. TLR7/8a-SNP | **0.0430** |
| SNP | vs. TLR1/2a-SNP | 0.0565 |
| SNP | vs. TLR4a-SNP | **0.0001** |
| SNP | vs. TLR7/8a-SNP | 0.4265 |
| TLR1/2a-SNP | vs. TLR4a-SNP | 0.2234 |
| TLR1/2a-SNP | vs. TLR7/8a-SNP | 0.8250 |
| TLR4a-SNP | vs. TLR7/8a-SNP | **0.0193** |

| Week 3<br>IgG titers [$log_{10}$] | | Adjusted *p* values |
|---|---|---|
| CpG/Alum | vs. SNP | 0.9875 |
| CpG/Alum | vs. TLR1/2a-SNP | **0.0013** |
| CpG/Alum | vs. TLR4a-SNP | **<0.0001** |
| CpG/Alum | vs. TLR7/8a-SNP | **0.0014** |
| SNP | vs. TLR1/2a-SNP | **0.0148** |
| SNP | vs. TLR4a-SNP | **0.0001** |
| SNP | vs. TLR7/8a-SNP | **0.0156** |
| TLR1/2a-SNP | vs. TLR4a-SNP | 0.4774 |
| TLR1/2a-SNP | vs. TLR7/8a-SNP | >0.9999 |
| TLR4a-SNP | vs. TLR7/8a-SNP | 0.4658 |

| Week 4<br>IgG titers [$log_{10}$] | | Adjusted *p* values |
|---|---|---|
| CpG/Alum | vs. SNP | 0.9898 |
| CpG/Alum | vs. TLR1/2a-SNP | **0.0010** |
| CpG/Alum | vs. TLR4a-SNP | **0.0153** |
| CpG/Alum | vs. TLR7/8a-SNP | **0.0088** |
| SNP | vs. TLR1/2a-SNP | **0.0108** |
| SNP | vs. TLR4a-SNP | 0.0810 |
| SNP | vs. TLR7/8a-SNP | 0.0590 |
| TLR1/2a-SNP | vs. TLR4a-SNP | 0.9770 |
| TLR1/2a-SNP | vs. TLR7/8a-SNP | 0.9692 |
| TLR4a-SNP | vs. TLR7/8a-SNP | >0.9999 |

| Week 5 IgG titers [$log_{10}$] | | Adjusted *p* values |
|---|---|---|
| CpG/Alum | vs. SNP | 0.8997 |
| CpG/Alum | vs. TLR1/2a-SNP | **<0.0001** |
| CpG/Alum | vs. TLR4a-SNP | **0.0456** |
| CpG/Alum | vs. TLR7/8a-SNP | **0.0356** |
| SNP | vs. TLR1/2a-SNP | **<0.0001** |
| SNP | vs. TLR4a-SNP | **0.0032** |
| SNP | vs. TLR7/8a-SNP | **0.0259** |
| TLR1/2a-SNP | vs. TLR4a-SNP | **0.0180** |
| TLR1/2a-SNP | vs. TLR7/8a-SNP | **0.0007** |
| TLR4a-SNP | vs. TLR7/8a-SNP | 0.9570 |

| Week 7 IgG titers [$log_{10}$] | | Adjusted *p* values |
|---|---|---|
| CpG/Alum | vs. SNP | 0.9677 |
| CpG/Alum | vs. TLR1/2a-SNP | **<0.0001** |
| CpG/Alum | vs. TLR4a-SNP | **0.0011** |
| CpG/Alum | vs. TLR7/8a-SNP | **0.0040** |
| SNP | vs. TLR1/2a-SNP | **0.0006** |
| SNP | vs. TLR4a-SNP | **0.0046** |
| SNP | vs. TLR7/8a-SNP | **0.0241** |
| TLR1/2a-SNP | vs. TLR4a-SNP | 0.8517 |
| TLR1/2a-SNP | vs. TLR7/8a-SNP | 0.2325 |
| TLR4a-SNP | vs. TLR7/8a-SNP | 0.8301 |

| Week 9 IgG titers [$log_{10}$] | | Adjusted *p* values |
|---|---|---|
| CpG/Alum | vs. SNP | 0.9987 |
| CpG/Alum | vs. TLR1/2a-SNP | **0.0005** |
| CpG/Alum | vs. TLR4a-SNP | **0.0087** |
| CpG/Alum | vs. TLR7/8a-SNP | **0.0157** |
| SNP | vs. TLR1/2a-SNP | **0.0087** |
| SNP | vs. TLR4a-SNP | **0.0181** |
| SNP | vs. TLR7/8a-SNP | **0.0430** |
| TLR1/2a-SNP | vs. TLR4a-SNP | 0.7886 |
| TLR1/2a-SNP | vs. TLR7/8a-SNP | 0.3792 |
| TLR4a-SNP | vs. TLR7/8a-SNP | 0.9736 |

| Week 11 IgG titers [$log_{10}$] | | Adjusted *p* values |
|---|---|---|
| CpG/Alum | vs. SNP | 0.9997 |
| CpG/Alum | vs. TLR1/2a-SNP | **0.0003** |
| CpG/Alum | vs. TLR4a-SNP | **0.0065** |
| CpG/Alum | vs. TLR7/8a-SNP | **0.0117** |
| SNP | vs. TLR1/2a-SNP | **0.0010** |
| SNP | vs. TLR4a-SNP | **0.0118** |
| SNP | vs. TLR7/8a-SNP | **0.0282** |
| TLR1/2a-SNP | vs. TLR4a-SNP | 0.7485 |
| TLR1/2a-SNP | vs. TLR7/8a-SNP | 0.3369 |
| TLR4a-SNP | vs. TLR7/8a-SNP | 0.9725 |

**Table S6.** *p* values from a general linear model (GLM) followed by Tukey's HSD multiple comparisons procedure for area under the curves (AUCs) of anti-RBD endpoint IgG titers compared between different RBD-NP vaccines (referring to Figure 2c).

| AUCs of titers from Week 0 to Week 11 [$log_{10}$] | | Adjusted *p* values |
|---|---|---|
| CpG/Alum | vs. SNP | 0.9898 |
| CpG/Alum | vs. TLR1/2a-SNP | **0.0018** |
| CpG/Alum | vs. TLR4a-SNP | **0.0267** |
| CpG/Alum | vs. TLR7/8a-SNP | **0.0197** |
| SNP | vs. TLR1/2a-SNP | **0.0082** |
| SNP | vs. TLR4a-SNP | 0.0742 |
| SNP | vs. TLR7/8a-SNP | 0.0747 |
| TLR1/2a-SNP | vs. TLR4a-SNP | 0.8071 |
| TLR1/2a-SNP | vs. TLR7/8a-SNP | 0.6984 |
| TLR4a-SNP | vs. TLR7/8a-SNP | >0.9999 |

**Table S7.** *p* values from a general linear model (GLM) followed by Tukey's HSD multiple comparisons procedure for Week 5 half-maximal binding dilution of anti-RBD IgG titers compared between different RBD-NP vaccines (referring to Figure 2d).

| Week 5<br>IgG $EC_{50}$ titers [$log_{10}$] | | Adjusted *p* values |
|---|---|---|
| CpG/Alum | vs. SNP | 0.8488 |
| CpG/Alum | vs. TLR1/2a-SNP | **<0.0001** |
| CpG/Alum | vs. TLR4a-SNP | 0.0945 |
| CpG/Alum | vs. TLR7/8a-SNP | **0.0447** |
| SNP | vs. TLR1/2a-SNP | **<0.0001** |
| SNP | vs. TLR4a-SNP | **0.0054** |
| SNP | vs. TLR7/8a-SNP | **0.0327** |
| TLR1/2a-SNP | vs. TLR4a-SNP | **0.0106** |
| TLR1/2a-SNP | vs. TLR7/8a-SNP | **0.0003** |
| TLR4a-SNP | vs. TLR7/8a-SNP | 0.9718 |

**Table S8.** *p* values from a general linear model (GLM) followed by Tukey's HSD multiple comparisons procedure for D365 anti-RBD endpoint IgG titers compared between different RBD-NP vaccines (referring to Figure 2e).

| D365 IgG titers [$log_{10}$] | | Adjusted *p* values |
|---|---|---|
| CpG/Alum | vs. SNP | >0.9999 |
| CpG/Alum | vs. TLR1/2a-SNP | **0.0108** |
| CpG/Alum | vs. TLR4a-SNP | **0.0198** |
| CpG/Alum | vs. TLR7/8a-SNP | - |
| SNP | vs. TLR1/2a-SNP | **0.0231** |
| SNP | vs. TLR4a-SNP | **0.0469** |
| SNP | vs. TLR7/8a-SNP | - |
| TLR1/2a-SNP | vs. TLR4a-SNP | 0.9940 |
| TLR1/2a-SNP | vs. TLR7/8a-SNP | - |
| TLR4a-SNP | vs. TLR7/8a-SNP | - |

*TLR7/8a-SNP vaccinated mice were excluded from the comparison due to an insufficient sample size that remained.

**Table S9.** *p* values from a general linear model (GLM) followed by Tukey HSD comparisons procedure for Week 5 anti-spike variant endpoint IgG titers compared between different CpG/Alum to other vaccine groups (referring to Figure 2f).

| WT Spike IgG titers [$log_{10}$] | | Adjusted *p* values |
|---|---|---|
| CpG/Alum | vs. SNP | 0.9467 |
| CpG/Alum | vs. TLR1/2a-SNP | **0.0145** |
| CpG/Alum | vs. TLR4a-SNP | 0.0716 |
| CpG/Alum | vs. TLR7/8a-SNP | 0.0531 |

| Alpha (B.1.1.7) IgG titers [$log_{10}$] | | Adjusted *p* values |
|---|---|---|
| CpG/Alum | vs. SNP | 0.9241 |
| CpG/Alum | vs. TLR1/2a-SNP | **0.0275** |
| CpG/Alum | vs. TLR4a-SNP | 0.0957 |
| CpG/Alum | vs. TLR7/8a-SNP | 0.0730 |

| Beta (B.1.351) IgG titers [$log_{10}$] | | Adjusted *p* values |
|---|---|---|
| CpG/Alum | vs. SNP | 0.9953 |
| CpG/Alum | vs. TLR1/2a-SNP | **0.0086** |
| CpG/Alum | vs. TLR4a-SNP | **0.0251** |
| CpG/Alum | vs. TLR7/8a-SNP | **0.0153** |

| Delta (B.1.617.2) IgG titers [$log_{10}$] | | Adjusted *p* values |
|---|---|---|
| CpG/Alum | vs. SNP | 0.7060 |
| CpG/Alum | vs. TLR1/2a-SNP | **0.0027** |
| CpG/Alum | vs. TLR4a-SNP | **0.0391** |
| CpG/Alum | vs. TLR7/8a-SNP | 0.0138 |

| Omicron (B.1.1.529) IgG titers [$log_{10}$] | | Adjusted *p* values |
|---|---|---|
| CpG/Alum | vs. SNP | 0.9981 |
| CpG/Alum | vs. TLR1/2a-SNP | **0.0068** |
| CpG/Alum | vs. TLR4a-SNP | 0.0567 |
| CpG/Alum | vs. TLR7/8a-SNP | 0.1367 |

**Table S10.** *p* values from a general linear model (GLM) followed by Tukey's HSD multiple comparisons procedure for Week 5 specific $ID_{50}$ neutralization titers compared between different RBD-NP vaccines (referring to Figure 3f).

| Week 5<br>Neutralization $EC_{50}$ titers [$log_{10}$] | | Adjusted *p* values |
|---|---|---|
| CpG/Alum | vs. SNP | 0.1329 |
| CpG/Alum | vs. TLR1/2a-SNP | **0.0033** |
| CpG/Alum | vs. TLR4a-SNP | **0.0008** |
| CpG/Alum | vs. TLR7/8a-SNP | **0.0048** |
| SNP | vs. TLR1/2a-SNP | 0.3735 |
| SNP | vs. TLR4a-SNP | 0.1545 |
| SNP | vs. TLR7/8a-SNP | 0.5527 |
| TLR1/2a-SNP | vs. TLR4a-SNP | 0.9892 |
| TLR1/2a-SNP | vs. TLR7/8a-SNP | 0.9927 |
| TLR4a-SNP | vs. TLR7/8a-SNP | 0.8749 |

**Table S11.** *p* values from a general linear model (GLM) followed by Tukey's HSD multiple comparisons procedure for relative percent infectivity at 1:100 dilution compared between different RBD-NP vaccines (referring to Figure 3g).

| Week 5<br>Relativity percent infectivity at 1:100 dilution | | Adjusted *p* values |
|---|---|---|
| No adjuvant | vs. Human Convalescent Serum (HCS) | **0.0078** |
| No adjuvant | vs CpG/Alum | 0.8295 |
| No adjuvant | vs. SNP | **0.0084** |
| No adjuvant | vs. TLR1/2a-SNP | **0.0009** |
| No adjuvant | vs. TLR4a-SNP | **0.0017** |
| No adjuvant | vs. TLR7/8a-SNP | **0.0008** |
| HCS | vs CpG/Alum | 0.2596 |
| HCS | vs. SNP | 0.9998 |
| HCS | vs. TLR1/2a-SNP | 0.9309 |
| HCS | vs. TLR4a-SNP | 0.9397 |
| HCS | vs. TLR7/8a-SNP | 0.9239 |
| CpG/Alum | vs. SNP | 0.2066 |
| CpG/Alum | vs. TLR1/2a-SNP | **0.0460** |
| CpG/Alum | vs. TLR4a-SNP | 0.0617 |
| CpG/Alum | vs. TLR7/8a-SNP | **0.0439** |
| SNP | vs. TLR1/2a-SNP | 0.9959 |
| SNP | vs. TLR4a-SNP | 0.9959 |
| SNP | vs. TLR7/8a-SNP | 0.9950 |
| TLR1/2a-SNP | vs. TLR4a-SNP | >0.9999 |
| TLR1/2a-SNP | vs. TLR7/8a-SNP | >0.9999 |
| TLR4a-SNP | vs. TLR7/8a-SNP | >0.9999 |

**Table S12.** *p* values from a general linear model (GLM) followed by Tukey's HSD multiple comparisons procedure for Week 5 anti-RBD endpoint IgG1 titers between different RBD-NP vaccines (referring to Figure 4a).

| Week 5 IgG1 titers [$\log_{10}$] | | Adjusted *p* values |
|---|---|---|
| CpG/Alum | vs. SNP | 0.9938 |
| CpG/Alum | vs. TLR1/2a-SNP | **0.0011** |
| CpG/Alum | vs. TLR4a-SNP | 0.7067 |
| CpG/Alum | vs. TLR7/8a-SNP | 0.3993 |
| SNP | vs. TLR1/2a-SNP | **0.0058** |
| SNP | vs. TLR4a-SNP | 0.9427 |
| SNP | vs. TLR7/8a-SNP | 0.7647 |
| TLR1/2a-SNP | vs. TLR4a-SNP | **0.0242** |
| TLR1/2a-SNP | vs. TLR7/8a-SNP | **0.0324** |
| TLR4a-SNP | vs. TLR7/8a-SNP | 0.9935 |

**Table S13.** *p* values from a general linear model (GLM) followed by Tukey's HSD multiple comparisons procedure for Week 5 anti-RBD endpoint IgG2c titers between different RBD-NP vaccines (referring to Figure 4b).

| Week 5 IgG2c titers [$log_{10}$] | | Adjusted *p* values |
|---|---|---|
| CpG/Alum | vs. SNP | 0.6517 |
| CpG/Alum | vs. TLR1/2a-SNP | **0.0064** |
| CpG/Alum | vs. TLR4a-SNP | **0.0081** |
| CpG/Alum | vs. TLR7/8a-SNP | **0.0039** |
| SNP | vs. TLR1/2a-SNP | 0.1389 |
| SNP | vs. TLR4a-SNP | 0.1180 |
| SNP | vs. TLR7/8a-SNP | 0.0908 |
| TLR1/2a-SNP | vs. TLR4a-SNP | 0.9997 |
| TLR1/2a-SNP | vs. TLR7/8a-SNP | 0.9987 |
| TLR4a-SNP | vs. TLR7/8a-SNP | >0.9999 |

**Table S14.** *p* values from a generalized maximum entropy estimation regression adjusted to control the false discovery rate for murine Luminex 48-plexed cytokine assay between SNP and TLRa-SNPs (referring to Figure 5b, Figure S8).

| BAFF<br>MFIs [$log_{10}$] | Adjusted<br>*p* values |
|---|---|
| SNP vs. TLR1/2a-SNP | 0.3429 |
| SNP vs. TLR4a-SNP | **<0.0001** |
| SNP vs. TLR7/8a-SNP | 0.2748 |

| ENA78<br>MFIs [$log_{10}$] | Adjusted<br>*p* values |
|---|---|
| SNP vs. TLR1/2a-SNP | 0.4066 |
| SNP vs. TLR4a-SNP | **0.0023** |
| SNP vs. TLR7/8a-SNP | 0.0730 |

| GCSF<br>MFIs [$log_{10}$] | Adjusted<br>*p* values |
|---|---|
| SNP vs. TLR1/2a-SNP | **<0.0001** |
| SNP vs. TLR4a-SNP | **<0.0001** |
| SNP vs. TLR7/8a-SNP | 0.5290 |

| GROa<br>MFIs [$log_{10}$] | Adjusted<br>*p* values |
|---|---|
| SNP vs. TLR1/2a-SNP | 0.5754 |
| SNP vs. TLR4a-SNP | **<0.0001** |
| SNP vs. TLR7/8a-SNP | 0.8858 |

| IFNγ<br>MFIs [$log_{10}$] | Adjusted<br>*p* values |
|---|---|
| SNP vs. TLR1/2a-SNP | **0.0015** |
| SNP vs. TLR4a-SNP | **<0.0001** |
| SNP vs. TLR7/8a-SNP | **0.0051** |

| IL12P70<br>MFIs [$log_{10}$] | Adjusted<br>*p* values |
|---|---|
| SNP vs. TLR1/2a-SNP | **0.0013** |
| SNP vs. TLR4a-SNP | 0.3350 |
| SNP vs. TLR7/8a-SNP | 0.2180 |

| IL15IL15R<br>MFIs [$log_{10}$] | Adjusted<br>*p* values |
|---|---|
| SNP vs. TLR1/2a-SNP | **<0.0001** |
| SNP vs. TLR4a-SNP | **<0.0001** |
| SNP vs. TLR7/8a-SNP | **<0.0001** |

| IL18<br>MFIs [$log_{10}$] | Adjusted<br>*p* values |
|---|---|
| SNP vs. TLR1/2a-SNP | 0.4066 |
| SNP vs. TLR4a-SNP | **<0.0001** |
| SNP vs. TLR7/8a-SNP | 0.1301 |

| BTC<br>MFIs [$log_{10}$] | Adjusted<br>*p* values |
|---|---|
| SNP vs. TLR1/2a-SNP | 0.5570 |
| SNP vs. TLR4a-SNP | 0.3432 |
| SNP vs. TLR7/8a-SNP | 0.2180 |

| EOTAXIN<br>MFIs [$log_{10}$] | Adjusted<br>*p* values |
|---|---|
| SNP vs. TLR1/2a-SNP | 0.0780 |
| SNP vs. TLR4a-SNP | 0.0845 |
| SNP vs. TLR7/8a-SNP | 0.2641 |

| GMCSF<br>MFIs [$log_{10}$] | Adjusted<br>*p* values |
|---|---|
| SNP vs. TLR1/2a-SNP | 0.2731 |
| SNP vs. TLR4a-SNP | **<0.0001** |
| SNP vs. TLR7/8a-SNP | 0.1301 |

| IFNα<br>MFIs [$log_{10}$] | Adjusted<br>*p* values |
|---|---|
| SNP vs. TLR1/2a-SNP | 0.3214 |
| SNP vs. TLR4a-SNP | 0.1188 |
| SNP vs. TLR7/8a-SNP | 0.1657 |

| IL10<br>MFIs [$log_{10}$] | Adjusted<br>*p* values |
|---|---|
| SNP vs. TLR1/2a-SNP | 0.6468 |
| SNP vs. TLR4a-SNP | 0.0762 |
| SNP vs. TLR7/8a-SNP | 0.3873 |

| IL13<br>MFIs [$log_{10}$] | Adjusted<br>*p* values |
|---|---|
| SNP vs. TLR1/2a-SNP | 0.8324 |
| SNP vs. TLR4a-SNP | 0.1143 |
| SNP vs. TLR7/8a-SNP | 0.0513 |

| IL17A<br>MFIs [$log_{10}$] | Adjusted<br>*p* values |
|---|---|
| SNP vs. TLR1/2a-SNP | 0.3918 |
| SNP vs. TLR4a-SNP | **<0.0001** |
| SNP vs. TLR7/8a-SNP | 0.2409 |

| IL19<br>MFIs [$log_{10}$] | Adjusted<br>*p* values |
|---|---|
| SNP vs. TLR1/2a-SNP | 0.4082 |
| SNP vs. TLR4a-SNP | **0.0087** |
| SNP vs. TLR7/8a-SNP | 0.1301 |

| IL1a MFIs [$log_{10}$] | Adjusted *p* values |
|---|---|
| SNP vs. TLR1/2a-SNP | 0.8535 |
| SNP vs. TLR4a-SNP | 0.4447 |
| SNP vs. TLR7/8a-SNP | 0.3313 |

| IL1b MFIs [$log_{10}$] | Adjusted *p* values |
|---|---|
| SNP vs. TLR1/2a-SNP | **<0.0001** |
| SNP vs. TLR4a-SNP | **<0.0001** |
| SNP vs. TLR7/8a-SNP | 0.0882 |

| IL2 MFIs [$log_{10}$] | Adjusted *p* values |
|---|---|
| SNP vs. TLR1/2a-SNP | 0.8646 |
| SNP vs. TLR4a-SNP | 0.7350 |
| SNP vs. TLR7/8a-SNP | 0.9397 |

| IL22 MFIs [$log_{10}$] | Adjusted *p* values |
|---|---|
| SNP vs. TLR1/2a-SNP | **<0.0001** |
| SNP vs. TLR4a-SNP | **<0.0001** |
| SNP vs. TLR7/8a-SNP | 0.0513 |

| IL23 MFIs [$log_{10}$] | Adjusted *p* values |
|---|---|
| SNP vs. TLR1/2a-SNP | 0.2731 |
| SNP vs. TLR4a-SNP | 1.0000 |
| SNP vs. TLR7/8a-SNP | 0.1673 |

| IL25IL17 MFIs [$log_{10}$] | Adjusted *p* values |
|---|---|
| SNP vs. TLR1/2a-SNP | **<0.0001** |
| SNP vs. TLR4a-SNP | **<0.0001** |
| SNP vs. TLR7/8a-SNP | 0.0882 |

| IL27 MFIs [$log_{10}$] | Adjusted *p* values |
|---|---|
| SNP vs. TLR1/2a-SNP | 0.2169 |
| SNP vs. TLR4a-SNP | **<0.0001** |
| SNP vs. TLR7/8a-SNP | 0.1673 |

| IL28 MFIs [$log_{10}$] | Adjusted *p* values |
|---|---|
| SNP vs. TLR1/2a-SNP | 1.0000 |
| SNP vs. TLR4a-SNP | 0.2796 |
| SNP vs. TLR7/8a-SNP | **0.0005** |

| IL2RA MFIs [$log_{10}$] | Adjusted *p* values |
|---|---|
| SNP vs. TLR1/2a-SNP | 0.3253 |
| SNP vs. TLR4a-SNP | **<0.0001** |
| SNP vs. TLR7/8a-SNP | 0.1698 |

| IL3 MFIs [$log_{10}$] | Adjusted *p* values |
|---|---|
| SNP vs. TLR1/2a-SNP | **0.0003** |
| SNP vs. TLR4a-SNP | **<0.0001** |
| SNP vs. TLR7/8a-SNP | **0.0041** |

| IL31 MFIs [$log_{10}$] | Adjusted *p* values |
|---|---|
| SNP vs. TLR1/2a-SNP | **0.0002** |
| SNP vs. TLR4a-SNP | **<0.0001** |
| SNP vs. TLR7/8a-SNP | 0.0730 |

| IL33 MFIs [$log_{10}$] | Adjusted *p* values |
|---|---|
| SNP vs. TLR1/2a-SNP | 0.5142 |
| SNP vs. TLR4a-SNP | **<0.0001** |
| SNP vs. TLR7/8a-SNP | 0.2194 |

| IL4 MFIs [$log_{10}$] | Adjusted *p* values |
|---|---|
| SNP vs. TLR1/2a-SNP | 0.1816 |
| SNP vs. TLR4a-SNP | 0.1502 |
| SNP vs. TLR7/8a-SNP | 0.1301 |

| IL5 MFIs [$log_{10}$] | Adjusted *p* values |
|---|---|
| SNP vs. TLR1/2a-SNP | 0.1816 |
| SNP vs. TLR4a-SNP | **0.0243** |
| SNP vs. TLR7/8a-SNP | 0.1301 |

| IL6 MFIs [$log_{10}$] | Adjusted *p* values |
|---|---|
| SNP vs. TLR1/2a-SNP | **0.0001** |
| SNP vs. TLR4a-SNP | **<0.0001** |
| SNP vs. TLR7/8a-SNP | 0.1301 |

| IL7 MFIs [$log_{10}$] | Adjusted *p* values |
|---|---|
| SNP vs. TLR1/2a-SNP | **0.0370** |
| SNP vs. TLR4a-SNP | 0.2701 |
| SNP vs. TLR7/8a-SNP | 0.2180 |

| IL7RA MFIs [$log_{10}$] | Adjusted *p* values |
|---|---|
| SNP vs. TLR1/2a-SNP | 0.2595 |
| SNP vs. TLR4a-SNP | **0.0074** |
| SNP vs. TLR7/8a-SNP | 0.0513 |

| IL9 MFIs [$log_{10}$] | Adjusted *p* values |
|---|---|
| SNP vs. TLR1/2a-SNP | 0.3022 |
| SNP vs. TLR4a-SNP | 0.9333 |
| SNP vs. TLR7/8a-SNP | **0.0005** |

| IP10 MFIs [$log_{10}$] | Adjusted *p* values |
|---|---|
| SNP vs. TLR1/2a-SNP | **0.0045** |
| SNP vs. TLR4a-SNP | **<0.0001** |
| SNP vs. TLR7/8a-SNP | **0.0005** |

| LIF MFIs [$log_{10}$] | Adjusted *p* values |
|---|---|
| SNP vs. TLR1/2a-SNP | 0.3672 |
| SNP vs. TLR4a-SNP | **<0.0001** |
| SNP vs. TLR7/8a-SNP | **0.0486** |

| MCP3 MFIs [$log_{10}$] | Adjusted *p* values |
|---|---|
| SNP vs. TLR1/2a-SNP | 0.8484 |
| SNP vs. TLR4a-SNP | **<0.0001** |
| SNP vs. TLR7/8a-SNP | 0.4645 |

| MIP1a MFIs [$log_{10}$] | Adjusted *p* values |
|---|---|
| SNP vs. TLR1/2a-SNP | 0.2967 |
| SNP vs. TLR4a-SNP | **<0.0001** |
| SNP vs. TLR7/8a-SNP | 0.1657 |

| MIP2 MFIs [$log_{10}$] | Adjusted *p* values |
|---|---|
| SNP vs. TLR1/2a-SNP | **0.0018** |
| SNP vs. TLR4a-SNP | **<0.0001** |
| SNP vs. TLR7/8a-SNP | 0.1961 |

| ST2 MFIs [$log_{10}$] | Adjusted *p* values |
|---|---|
| SNP vs. TLR1/2a-SNP | 0.3946 |
| SNP vs. TLR4a-SNP | **0.0027** |
| SNP vs. TLR7/8a-SNP | 0.1657 |

| VEGF MFIs [$log_{10}$] | Adjusted *p* values |
|---|---|
| SNP vs. TLR1/2a-SNP | 0.9420 |
| SNP vs. TLR4a-SNP | **0.0272** |
| SNP vs. TLR7/8a-SNP | 0.1454 |

| LEPTIN MFIs [$log_{10}$] | Adjusted *p* values |
|---|---|
| SNP vs. TLR1/2a-SNP | 0.4370 |
| SNP vs. TLR4a-SNP | 0.5943 |
| SNP vs. TLR7/8a-SNP | 0.1698 |

| MCP1 MFIs [$log_{10}$] | Adjusted *p* values |
|---|---|
| SNP vs. TLR1/2a-SNP | 0.8739 |
| SNP vs. TLR4a-SNP | **<0.0001** |
| SNP vs. TLR7/8a-SNP | 0.6804 |

| MCSF MFIs [$log_{10}$] | Adjusted *p* values |
|---|---|
| SNP vs. TLR1/2a-SNP | 0.3946 |
| SNP vs. TLR4a-SNP | **0.0001** |
| SNP vs. TLR7/8a-SNP | 0.1698 |

| MIP1b MFIs [$log_{10}$] | Adjusted *p* values |
|---|---|
| SNP vs. TLR1/2a-SNP | 0.5605 |
| SNP vs. TLR4a-SNP | **<0.0001** |
| SNP vs. TLR7/8a-SNP | 0.1961 |

| RANTES MFIs [$log_{10}$] | Adjusted *p* values |
|---|---|
| SNP vs. TLR1/2a-SNP | 0.1563 |
| SNP vs. TLR4a-SNP | **<0.0001** |
| SNP vs. TLR7/8a-SNP | 0.0513 |

| TNFa MFIs [$log_{10}$] | Adjusted *p* values |
|---|---|
| SNP vs. TLR1/2a-SNP | 0.5936 |
| SNP vs. TLR4a-SNP | 0.0820 |
| SNP vs. TLR7/8a-SNP | 0.1961 |

| sRANKL MFIs [$log_{10}$] | Adjusted *p* values |
|---|---|
| SNP vs. TLR1/2a-SNP | **0.0162** |
| SNP vs. TLR4a-SNP | **0.0001** |
| SNP vs. TLR7/8a-SNP | 0.5327 |

**Table S15.** *p* values from a general linear model (GLM) followed by Tukey's HSD multiple comparisons procedure for GCBC count compared between different RBD-NP vaccines (referring to Figure 5d).

| GCBC count [$log_{10}$] | | Adjusted *p* values |
|---|---|---|
| CpG/Alum | vs. SNP | 0.9767 |
| CpG/Alum | vs. TLR1/2a-SNP | 0.9979 |
| CpG/Alum | vs. TLR4a-SNP | 0.9962 |
| CpG/Alum | vs. TLR7/8a-SNP | 0.9406 |
| SNP | vs. TLR1/2a-SNP | 0.9987 |
| SNP | vs. TLR4a-SNP | 0.8788 |
| SNP | vs. TLR7/8a-SNP | 0.9998 |
| TLR1/2a-SNP | vs. TLR4a-SNP | 0.9614 |
| TLR1/2a-SNP | vs. TLR7/8a-SNP | 0.9911 |
| TLR4a-SNP | vs. TLR7/8a-SNP | 0.7978 |

**Table S16.** *p* values from a general linear model (GLM) followed by Tukey's HSD multiple comparisons procedure for % GCBC of B cells compared between different RBD-NP vaccines (referring to Figure 5e).

| % GCBC of B cells | | Adjusted *p* values |
|---|---|---|
| CpG/Alum | vs. SNP | 0.4379 |
| CpG/Alum | vs. TLR1/2a-SNP | 0.4563 |
| CpG/Alum | vs. TLR4a-SNP | 0.0901 |
| CpG/Alum | vs. TLR7/8a-SNP | 0.1486 |
| SNP | vs. TLR1/2a-SNP | >0.9999 |
| SNP | vs. TLR4a-SNP | 0.8678 |
| SNP | vs. TLR7/8a-SNP | 0.9552 |
| TLR1/2a-SNP | vs. TLR4a-SNP | 0.8536 |
| TLR1/2a-SNP | vs. TLR7/8a-SNP | 0.9474 |
| TLR4a-SNP | vs. TLR7/8a-SNP | 0.9987 |

**Table S17.** *p* values from a general linear model (GLM) followed by Tukey's HSD multiple comparisons procedure for Tfh count compared between different RBD-NP vaccines (referring to Figure S10).

| Tfh count [$\log_{10}$] | | Adjusted *p* values |
|---|---|---|
| CpG/Alum | vs. SNP | 0.9767 |
| CpG/Alum | vs. TLR1/2a-SNP | 0.9979 |
| CpG/Alum | vs. TLR4a-SNP | 0.9962 |
| CpG/Alum | vs. TLR7/8a-SNP | 0.9406 |
| SNP | vs. TLR1/2a-SNP | 0.9987 |
| SNP | vs. TLR4a-SNP | 0.8788 |
| SNP | vs. TLR7/8a-SNP | 0.9998 |
| TLR1/2a-SNP | vs. TLR4a-SNP | 0.9614 |
| TLR1/2a-SNP | vs. TLR7/8a-SNP | 0.9911 |
| TLR4a-SNP | vs. TLR7/8a-SNP | 0.7978 |

**Table S18.** *p* values from a general linear model (GLM) followed by Tukey's HSD multiple comparisons procedure for GCBC to Tfh ratio compared between different RBD-NP vaccines (referring to Figure 5f).

| Tfh count [$log_{10}$] | | Adjusted *p* values |
|---|---|---|
| CpG/Alum | vs. SNP | 0.3562 |
| CpG/Alum | vs. TLR1/2a-SNP | 0.5906 |
| CpG/Alum | vs. TLR4a-SNP | 0.7240 |
| CpG/Alum | vs. TLR7/8a-SNP | 0.1037 |
| SNP | vs. TLR1/2a-SNP | 0.9934 |
| SNP | vs. TLR4a-SNP | 0.9670 |
| SNP | vs. TLR7/8a-SNP | 0.9452 |
| TLR1/2a-SNP | vs. TLR4a-SNP | 0.9994 |
| TLR1/2a-SNP | vs. TLR7/8a-SNP | 0.7786 |
| TLR4a-SNP | vs. TLR7/8a-SNP | 0.6502 |

**Table S19.** *p* values from a general linear model (GLM) followed by Tukey's HSD multiple comparisons procedure for specific anti-RBD endpoint IgG titers time points compared between different RBD-NP vaccines (referring to Figure 6c).

| Week 2<br>IgG titers [$log_{10}$] | | Adjusted *p* values |
|---|---|---|
| SNP | vs TLR4a + TLR7/8a-SNP | **0.0257** |
| SNP | vs. TLR4a-TLR7/8a-SNP | **0.0083** |
| TLR4a-SNP | vs TLR4a-SNP + TLR7/8a-SNP | 0.9986 |
| TLR4a-SNP | vs. TLR4a-TLR7/8a-SNP | 0.9972 |
| TLR7/8a-SNP | vs TLR4a-SNP + TLR7/8a-SNP | 0.3206 |
| TLR7/8a-SNP | vs. TLR4a-TLR7/8a-SNP | 0.1154 |
| TLR4a-SNP + TLR7/8a-SNP | vs. TLR4a-TLR7/8a-SNP | 0.9630 |

| Week 3<br>IgG titers [$log_{10}$] | | Adjusted *p* values |
|---|---|---|
| SNP | vs TLR4a + TLR7/8a-SNP | **0.0434** |
| SNP | vs. TLR4a-TLR7/8a-SNP | **0.0016** |
| TLR4a-SNP | vs TLR4a-SNP + TLR7/8a-SNP | 0.8930 |
| TLR4a-SNP | vs. TLR4a-TLR7/8a-SNP | 0.8962 |
| TLR7/8a-SNP | vs TLR4a-SNP + TLR7/8a-SNP | 0.9750 |
| TLR7/8a-SNP | vs. TLR4a-TLR7/8a-SNP | 0.1424 |
| TLR4a-SNP + TLR7/8a-SNP | vs. TLR4a-TLR7/8a-SNP | 0.3467 |

| Week 4<br>IgG titers [$log_{10}$] | | Adjusted *p* values |
|---|---|---|
| SNP | vs TLR4a + TLR7/8a-SNP | **0.0236** |
| SNP | vs. TLR4a-TLR7/8a-SNP | **0.0029** |
| TLR4a-SNP | vs TLR4a-SNP + TLR7/8a-SNP | 0.9990 |
| TLR4a-SNP | vs. TLR4a-TLR7/8a-SNP | 0.9003 |
| TLR7/8a-SNP | vs TLR4a-SNP + TLR7/8a-SNP | 0.9964 |
| TLR7/8a-SNP | vs. TLR4a-TLR7/8a-SNP | 0.9031 |
| TLR4a-SNP + TLR7/8a-SNP | vs. TLR4a-TLR7/8a-SNP | 0.7405 |

| Week 5<br>IgG titers [$log_{10}$] | | Adjusted *p* values |
|---|---|---|
| SNP | vs TLR4a + TLR7/8a-SNP | **0.0175** |
| SNP | vs. TLR4a-TLR7/8a-SNP | **0.0001** |
| TLR4a-SNP | vs TLR4a-SNP + TLR7/8a-SNP | 0.9975 |
| TLR4a-SNP | vs. TLR4a-TLR7/8a-SNP | **0.0445** |
| TLR7/8a-SNP | vs TLR4a-SNP + TLR7/8a-SNP | >0.9999 |
| TLR7/8a-SNP | vs. TLR4a-TLR7/8a-SNP | **0.0431** |
| TLR4a-SNP + TLR7/8a-SNP | vs. TLR4a-TLR7/8a-SNP | 0.0524 |

| Week 7<br>IgG titers [$log_{10}$] | | Adjusted *p* values |
|---|---|---|
| SNP | vs TLR4a + TLR7/8a-SNP | **0.0058** |
| SNP | vs. TLR4a-TLR7/8a-SNP | **<0.0001** |
| TLR4a-SNP | vs TLR4a-SNP + TLR7/8a-SNP | 0.9997 |
| TLR4a-SNP | vs. TLR4a-TLR7/8a-SNP | 0.1119 |
| TLR7/8a-SNP | vs TLR4a-SNP + TLR7/8a-SNP | 0.7234 |
| TLR7/8a-SNP | vs. TLR4a-TLR7/8a-SNP | **0.0110** |
| TLR4a-SNP + TLR7/8a-SNP | vs. TLR4a-TLR7/8a-SNP | 0.1082 |

**Table S20.** *p* values from a general linear model (GLM) followed by Tukey's HSD multiple comparisons procedure for Week 5 anti-RBD endpoint IgG1 titers between different RBD-NP vaccines (referring to Figure 6d).

| Week 5<br>IgG1 titers [$log_{10}$] | | Adjusted *p* values |
|---|---|---|
| SNP | vs TLR4a + TLR7/8a-SNP | 0.6642 |
| SNP | vs. TLR4a-TLR7/8a-SNP | 0.1107 |
| TLR4a-SNP | vs TLR4a-SNP + TLR7/8a-SNP | 0.9879 |
| TLR4a-SNP | vs. TLR4a-TLR7/8a-SNP | 0.4149 |
| TLR7/8a-SNP | vs TLR4a-SNP + TLR7/8a-SNP | 0.9998 |
| TLR7/8a-SNP | vs. TLR4a-TLR7/8a-SNP | 0.5287 |
| TLR4a-SNP + TLR7/8a-SNP | vs. TLR4a-TLR7/8a-SNP | 0.6274 |

**Table S21.** *p* values from a general linear model (GLM) followed by Tukey's HSD multiple comparisons procedure for Week 5 anti-RBD endpoint IgG2c titers between different RBD-NP vaccines (referring to Figure 6e).

| Week 5<br>IgG2c titers [$log_{10}$] | | Adjusted *p* values |
|---|---|---|
| SNP | vs TLR4a + TLR7/8a-SNP | **0.0002** |
| SNP | vs. TLR4a-TLR7/8a-SNP | **<0.0001** |
| TLR4a-SNP | vs TLR4a-SNP + TLR7/8a-SNP | 0.2990 |
| TLR4a-SNP | vs. TLR4a-TLR7/8a-SNP | 0.0543 |
| TLR7/8a-SNP | vs TLR4a-SNP + TLR7/8a-SNP | 0.2083 |
| TLR7/8a-SNP | vs. TLR4a-TLR7/8a-SNP | **0.0299** |
| TLR4a-SNP + TLR7/8a-SNP | vs. TLR4a-TLR7/8a-SNP | 0.7956 |

**Table S22.** *p* values from a general linear model (GLM) followed by Tukey's HSD multiple comparisons procedure for Week 7 anti-RBD antibody affinity ($K_D$) between different RBD-NP vaccines (referring to Figure 6h).

| $K_D$ (nM) [$log_{10}$] | | Adjusted *p* values |
|---|---|---|
| CpG/Alum | vs. SNP | 0.9763 |
| CpG/Alum | vs. TLR1/2a-SNP | 0.8973 |
| CpG/Alum | vs. TLR4a-SNP | 0.7824 |
| CpG/Alum | vs. TLR7/8a-SNP | 0.9961 |
| CpG/Alum | vs. TLR4a-TLR7/8a-SNP | 0.1226 |
| SNP | vs. TLR1/2a-SNP | 0.4019 |
| SNP | vs. TLR4a-SNP | 0.2961 |
| SNP | vs. TLR7/8a-SNP | 0.7420 |
| SNP | vs. TLR4a-TLR7/8a-SNP | **0.0169** |
| TLR1/2a-SNP | vs. TLR4a-SNP | 0.9999 |
| TLR1/2a-SNP | vs. TLR7/8a-SNP | 0.9960 |
| TLR1/2a-SNP | vs. TLR4a-TLR7/8a-SNP | 0.6436 |
| TLR4a-SNP | vs. TLR7/8a-SNP | 0.9688 |
| TLR4a-SNP | vs. TLR4a-TLR7/8a-SNP | 0.8798 |
| TLR7/8a-SNP | vs. TLR4a-TLR7/8a-SNP | 0.2873 |

**Table S23.** *p* values from a general linear model (GLM) followed by Tukey's HSD multiple comparisons procedure for specific anti-gp120 endpoint IgG titers time points compared between different gp120 vaccines (referring to Figure 7b and S12b).

| Week 2 IgG titers [$log_{10}$] | | Adjusted *p* values |
|---|---|---|
| Alum | vs. SNP | 0.9904 |
| Alum | vs. TLR1/2a-SNP | **0.0123** |
| Alum | vs. TLR4a-SNP | **0.0100** |
| Alum | vs. TLR7/8a-SNP | **0.0248** |
| Alum | vs. TLR4a-TLR7/8a-SNP | **<0.0001** |
| SNP | vs. TLR1/2a-SNP | **0.0017** |
| SNP | vs. TLR4a-SNP | **0.0001** |
| SNP | vs. TLR7/8a-SNP | **0.0037** |
| SNP | vs. TLR4a-TLR7/8a-SNP | **<0.0001** |
| TLR1/2a-SNP | vs. TLR4a-SNP | 0.9766 |
| TLR1/2a-SNP | vs. TLR7/8a-SNP | 0.9999 |
| TLR1/2a-SNP | vs. TLR4a-TLR7/8a-SNP | 0.4900 |
| TLR4a-SNP | vs. TLR7/8a-SNP | 0.9254 |
| TLR4a-SNP | vs. TLR4a-TLR7/8a-SNP | 0.9062 |
| TLR7/8a-SNP | vs. TLR4a-TLR7/8a-SNP | 0.3477 |

| Week 3 IgG titers [$log_{10}$] | | Adjusted *p* values |
|---|---|---|
| Alum | vs. SNP | 0.7683 |
| Alum | vs. TLR1/2a-SNP | **<0.0001** |
| Alum | vs. TLR4a-SNP | **<0.0001** |
| Alum | vs. TLR7/8a-SNP | **<0.0001** |
| Alum | vs. TLR4a-TLR7/8a-SNP | **<0.0001** |
| SNP | vs. TLR1/2a-SNP | **0.0011** |
| SNP | vs. TLR4a-SNP | **<0.0001** |
| SNP | vs. TLR7/8a-SNP | **<0.0001** |
| SNP | vs. TLR4a-TLR7/8a-SNP | **<0.0001** |
| TLR1/2a-SNP | vs. TLR4a-SNP | 0.5602 |
| TLR1/2a-SNP | vs. TLR7/8a-SNP | 0.9924 |
| TLR1/2a-SNP | vs. TLR4a-TLR7/8a-SNP | 0.0630 |
| TLR4a-SNP | vs. TLR7/8a-SNP | 0.9813 |
| TLR4a-SNP | vs. TLR4a-TLR7/8a-SNP | 0.8559 |
| TLR7/8a-SNP | vs. TLR4a-TLR7/8a-SNP | 0.4336 |

| Week 4 IgG titers [$log_{10}$] | | Adjusted *p* values |
|---|---|---|
| Alum | vs. SNP | 0.3801 |
| Alum | vs. TLR1/2a-SNP | **<0.0001** |
| Alum | vs. TLR4a-SNP | **<0.0001** |
| Alum | vs. TLR7/8a-SNP | **<0.0001** |
| Alum | vs. TLR4a-TLR7/8a-SNP | **<0.0001** |
| SNP | vs. TLR1/2a-SNP | **0.0016** |
| SNP | vs. TLR4a-SNP | **<0.0001** |
| SNP | vs. TLR7/8a-SNP | **<0.0001** |
| SNP | vs. TLR4a-TLR7/8a-SNP | **<0.0001** |
| TLR1/2a-SNP | vs. TLR4a-SNP | 0.5296 |
| TLR1/2a-SNP | vs. TLR7/8a-SNP | 0.6981 |
| TLR1/2a-SNP | vs. TLR4a-TLR7/8a-SNP | **0.0054** |
| TLR4a-SNP | vs. TLR7/8a-SNP | 0.9998 |
| TLR4a-SNP | vs. TLR4a-TLR7/8a-SNP | 0.3791 |
| TLR7/8a-SNP | vs. TLR4a-TLR7/8a-SNP | 0.2426 |

| Week 6 IgG titers [$log_{10}$] | | Adjusted *p* values |
|---|---|---|
| Alum | vs. SNP | **<0.0001** |
| Alum | vs. TLR1/2a-SNP | **<0.0001** |
| Alum | vs. TLR4a-SNP | **<0.0001** |
| Alum | vs. TLR7/8a-SNP | **<0.0001** |
| Alum | vs. TLR4a-TLR7/8a-SNP | **<0.0001** |
| SNP | vs. TLR1/2a-SNP | 0.0803 |
| SNP | vs. TLR4a-SNP | 0.0995 |
| SNP | vs. TLR7/8a-SNP | 0.0901 |
| SNP | vs. TLR4a-TLR7/8a-SNP | **0.0007** |
| TLR1/2a-SNP | vs. TLR4a-SNP | >0.9999 |
| TLR1/2a-SNP | vs. TLR7/8a-SNP | >0.9999 |
| TLR1/2a-SNP | vs. TLR4a-TLR7/8a-SNP | 0.2915 |
| TLR4a-SNP | vs. TLR7/8a-SNP | >0.9999 |
| TLR4a-SNP | vs. TLR4a-TLR7/8a-SNP | 0.2451 |
| TLR7/8a-SNP | vs. TLR4a-TLR7/8a-SNP | 0.2661 |

**Table S24.** *p* values from a general linear model (GLM) followed by Tukey's HSD multiple comparisons procedure for area under the curves (AUCs) of anti-gp120 endpoint IgG titers compared between different gp120 vaccines (referring to Figure 7c).

| AUCs of titers from Week 0 to Week 11 [$\log_{10}$] | | Adjusted *p* values |
|---|---|---|
| Alum | vs. SNP | **<0.0001** |
| Alum | vs. TLR1/2a-SNP | **<0.0001** |
| Alum | vs. TLR4a-SNP | **<0.0001** |
| Alum | vs. TLR7/8a-SNP | **<0.0001** |
| Alum | vs. TLR4a-TLR7/8a-SNP | **<0.0001** |
| SNP | vs. TLR1/2a-SNP | **0.0416** |
| SNP | vs. TLR4a-SNP | **0.0489** |
| SNP | vs. TLR7/8a-SNP | 0.6451 |
| SNP | vs. TLR4a-TLR7/8a-SNP | **0.0125** |
| TLR1/2a-SNP | vs. TLR4a-SNP | >0.9999 |
| TLR1/2a-SNP | vs. TLR7/8a-SNP | 0.4151 |
| TLR1/2a-SNP | vs. TLR4a-TLR7/8a-SNP | 0.9893 |
| TLR4a-SNP | vs. TLR7/8a-SNP | 0.4559 |
| TLR4a-SNP | vs. TLR4a-TLR7/8a-SNP | 0.9825 |
| TLR7/8a-SNP | vs. TLR4a-TLR7/8a-SNP | 0.1554 |

**Table S25.** *p* values from a general linear model (GLM) followed by Tukey's HSD multiple comparisons procedure for Week 5 anti-gp120 endpoint IgG1 titers between different gp120 vaccines (referring to Figure 7d).

| Week 6<br>IgG1 titers [$log_{10}$] | | Adjusted *p* values |
|---|---|---|
| SNP | vs. TLR1/2a-SNP | **0.0055** |
| SNP | vs. TLR4a-SNP | 0.0874 |
| SNP | vs. TLR7/8a-SNP | 0.4234 |
| SNP | vs. TLR4a-TLR7/8a-SNP | **<0.0001** |
| TLR1/2a-SNP | vs. TLR4a-SNP | 0.6264 |
| TLR1/2a-SNP | vs. TLR7/8a-SNP | 0.1622 |
| TLR1/2a-SNP | vs. TLR4a-TLR7/8a-SNP | 0.0997 |
| TLR4a-SNP | vs. TLR7/8a-SNP | 0.8534 |
| TLR4a-SNP | vs. TLR4a-TLR7/8a-SNP | **0.0063** |
| TLR7/8a-SNP | vs. TLR4a-TLR7/8a-SNP | **0.0009** |

**Table S26.** *p* values from a general linear model (GLM) followed by Tukey's HSD multiple comparisons procedure for Week 5 anti-gp120 endpoint IgG2c titers between different gp120 vaccines (referring to Figure 7e).

| Week 6<br>IgG2c titers [$log_{10}$] | | Adjusted *p* values |
|---|---|---|
| SNP | vs. TLR1/2a-SNP | 0.0574 |
| SNP | vs. TLR4a-SNP | **0.0194** |
| SNP | vs. TLR7/8a-SNP | **0.0133** |
| SNP | vs. TLR4a-TLR7/8a-SNP | **<0.0001** |
| TLR1/2a-SNP | vs. TLR4a-SNP | 0.9791 |
| TLR1/2a-SNP | vs. TLR7/8a-SNP | 0.9414 |
| TLR1/2a-SNP | vs. TLR4a-TLR7/8a-SNP | **0.0003** |
| TLR4a-SNP | vs. TLR7/8a-SNP | 0.9997 |
| TLR4a-SNP | vs. TLR4a-TLR7/8a-SNP | **0.0008** |
| TLR7/8a-SNP | vs. TLR4a-TLR7/8a-SNP | **0.0012** |

**Table S27.** *p* values from a general linear model (GLM) followed by Tukey's HSD multiple comparisons procedure for GCBC count compared between different gp120 vaccines (referring to Figure 7g).

| GCBC count [$log_{10}$] | | Adjusted *p* values |
|---|---|---|
| Alum | vs. SNP | **0.0039** |
| Alum | vs. TLR1/2a-SNP | **<0.0001** |
| Alum | vs. TLR4a-SNP | **0.0012** |
| Alum | vs. TLR7/8a-SNP | **0.0023** |
| Alum | vs. TLR4a-TLR7/8a-SNP | **0.0012** |
| SNP | vs. TLR1/2a-SNP | 0.4930 |
| SNP | vs. TLR4a-SNP | 0.9980 |
| SNP | vs. TLR7/8a-SNP | 0.9999 |
| SNP | vs. TLR4a-TLR7/8a-SNP | 0.9974 |
| TLR1/2a-SNP | vs. TLR4a-SNP | 0.7534 |
| TLR1/2a-SNP | vs. TLR7/8a-SNP | 0.6115 |
| TLR1/2a-SNP | vs. TLR4a-TLR7/8a-SNP | 0.7669 |
| TLR4a-SNP | vs. TLR7/8a-SNP | 0.9999 |
| TLR4a-SNP | vs. TLR4a-TLR7/8a-SNP | 0.9999 |
| TLR7/8a-SNP | vs. TLR4a-TLR7/8a-SNP | 0.9998 |

**Table S28.** *p* values from a general linear model (GLM) followed by Tukey's HSD multiple comparisons procedure for % GCBC of B cells compared between different RBD-NP vaccines (referring to Figure 7h).

| % GCBC of B cells | | Adjusted *p* values |
|---|---|---|
| Alum | vs. SNP | **0.0202** |
| Alum | vs. TLR1/2a-SNP | **<0.0001** |
| Alum | vs. TLR4a-SNP | **0.0011** |
| Alum | vs. TLR7/8a-SNP | **0.0084** |
| Alum | vs. TLR4a-TLR7/8a-SNP | **<0.0001** |
| SNP | vs. TLR1/2a-SNP | **0.0394** |
| SNP | vs. TLR4a-SNP | 0.8693 |
| SNP | vs. TLR7/8a-SNP | 0.9992 |
| SNP | vs. TLR4a-TLR7/8a-SNP | 0.3367 |
| TLR1/2a-SNP | vs. TLR4a-SNP | 0.3524 |
| TLR1/2a-SNP | vs. TLR7/8a-SNP | 0.0857 |
| TLR1/2a-SNP | vs. TLR4a-TLR7/8a-SNP | 0.8693 |
| TLR4a-SNP | vs. TLR7/8a-SNP | 0.9703 |
| TLR4a-SNP | vs. TLR4a-TLR7/8a-SNP | 0.9421 |
| TLR7/8a-SNP | vs. TLR4a-TLR7/8a-SNP | 0.5568 |

**Table S29.** *p* values from a general linear model (GLM) followed by Tukey's HSD multiple comparisons procedure for specific anti-gp120 endpoint IgG titers time points compared between different gp120 vaccines in male mice (referring to Figure S12c).

| Week 2<br>IgG titers [$log_{10}$] | | Adjusted *p* values |
|---|---|---|
| SNP | vs. TLR1/2a-SNP | **0.0436** |
| SNP | vs. TLR4a-SNP | **0.0079** |
| SNP | vs. TLR7/8a-SNP | 0.2080 |
| SNP | vs. TLR4a-TLR7/8a-SNP | **0.0103** |
| TLR1/2a-SNP | vs. TLR4a-SNP | 0.9716 |
| TLR1/2a-SNP | vs. TLR7/8a-SNP | 0.9548 |
| TLR1/2a-SNP | vs. TLR4a-TLR7/8a-SNP | 0.9839 |
| TLR4a-SNP | vs. TLR7/8a-SNP | 0.6784 |
| TLR4a-SNP | vs. TLR4a-TLR7/8a-SNP | 0.9999 |
| TLR7/8a-SNP | vs. TLR4a-TLR7/8a-SNP | 0.7317 |

| Week 3<br>IgG titers [$log_{10}$] | | Adjusted *p* values |
|---|---|---|
| SNP | vs. TLR1/2a-SNP | **0.0187** |
| SNP | vs. TLR4a-SNP | **0.0042** |
| SNP | vs. TLR7/8a-SNP | 0.0980 |
| SNP | vs. TLR4a-TLR7/8a-SNP | **0.0014** |
| TLR1/2a-SNP | vs. TLR4a-SNP | 0.9863 |
| TLR1/2a-SNP | vs. TLR7/8a-SNP | 0.9642 |
| TLR1/2a-SNP | vs. TLR4a-TLR7/8a-SNP | 0.9127 |
| TLR4a-SNP | vs. TLR7/8a-SNP | 0.7698 |
| TLR4a-SNP | vs. TLR4a-TLR7/8a-SNP | 0.9969 |
| TLR7/8a-SNP | vs. TLR4a-TLR7/8a-SNP | 0.5587 |

| Week 4<br>IgG titers [$log_{10}$] | | Adjusted *p* values |
|---|---|---|
| SNP | vs. TLR1/2a-SNP | **0.0188** |
| SNP | vs. TLR4a-SNP | **0.0032** |
| SNP | vs. TLR7/8a-SNP | **0.0296** |
| SNP | vs. TLR4a-TLR7/8a-SNP | **0.0007** |
| TLR1/2a-SNP | vs. TLR4a-SNP | 0.9752 |
| TLR1/2a-SNP | vs. TLR7/8a-SNP | 0.9998 |
| TLR1/2a-SNP | vs. TLR4a-TLR7/8a-SNP | 0.8324 |
| TLR4a-SNP | vs. TLR7/8a-SNP | 0.7911 |
| TLR4a-SNP | vs. TLR4a-TLR7/8a-SNP | 0.9999 |
| TLR7/8a-SNP | vs. TLR4a-TLR7/8a-SNP | 0.8831 |

| Week 6<br>IgG titers [$log_{10}$] | | Adjusted *p* values |
|---|---|---|
| SNP | vs. TLR1/2a-SNP | **0.0009** |
| SNP | vs. TLR4a-SNP | **0.0013** |
| SNP | vs. TLR7/8a-SNP | **0.0369** |
| SNP | vs. TLR4a-TLR7/8a-SNP | **0.0005** |
| TLR1/2a-SNP | vs. TLR4a-SNP | 0.9999 |
| TLR1/2a-SNP | vs. TLR7/8a-SNP | 0.7279 |
| TLR1/2a-SNP | vs. TLR4a-TLR7/8a-SNP | 0.9980 |
| TLR4a-SNP | vs. TLR7/8a-SNP | 0.7911 |
| TLR4a-SNP | vs. TLR4a-TLR7/8a-SNP | 0.9997 |
| TLR7/8a-SNP | vs. TLR4a-TLR7/8a-SNP | 0.8831 |

**Table S30.** *p* values from a one-way ANOVA followed by Tukey multiple comparison test for IFN-γ producing splenocyte counts compared between different RBD-NP vaccines (referring to Figure S13).

| Count | | Adjusted *p* values |
|---|---|---|
| CpG/Alum | vs. SNP | 0.4717 |
| CpG/Alum | vs. TLR1/2a-SNP | 0.8714 |
| CpG/Alum | vs. TLR4a-SNP | 0.9985 |
| CpG/Alum | vs. TLR7/8a-SNP | 0.9437 |
| SNP | vs. TLR1/2a-SNP | 0.0962 |
| SNP | vs. TLR4a-SNP | 0.6439 |
| SNP | vs. TLR7/8a-SNP | 0.8858 |
| TLR1/2a-SNP | vs. TLR4a-SNP | 0.7282 |
| TLR1/2a-SNP | vs. TLR7/8a-SNP | 0.4510 |
| TLR4a-SNP | vs. TLR7/8a-SNP | 0.9901 |

#### Supplemental Data

Source data presented in Figures 1-7, and supplemental figures are presented in the attached Excel Data file S1, “adn7187_Suppl. Excel_seq1_v1.xlsx.”
